# Bridging the Babble: A Method to Correlate Varying Nomenclature and Surface Marker Definitions Across the Human Splenic B Cell Literature

**DOI:** 10.64898/2026.11.15.766777

**Authors:** Molly E. Niska, Qian Zhou, W. Richard Burack, Mark Y. Sangster, Martin S. Zand

**Author notes:** Correspondence: Department of Medicine, Division of Nephrology, 601 Elmwood Ave - Box 675, Rochester, NY 14642, USA, martin.

## Abstract

**Introduction:** Flow cytometry-based immunophenotyping is widely used for identifying memory B cell subsets associated with disease or immune responses. For simplicity, multi-feature surface marker vectors are often reduced to a single subset label for cell sorting, functional tests, and understanding immune cell interactions. Across the literature however, the same label is often used with significantly different phenotype feature vectors. Unfortunately, this babelization of phenotype definitions, makes it difficult to compare experimental results across studies. This becomes a significant reproducibility issue when cells reside in multiple anatomical compartments, with location-dependent phenotypes and naming conventions.

**Methods:** Here, we report a broadly applicable, rigorous computational pipeline for analyzing flow cytometry data, with a novel implementation of the Jaccard Similarity Index (JSI) to compare gating schemes. The JSI is used to map the labels assigned to each cell when two gating schemes are applied to the same dataset. We designed a human splenic B cell flow cytometry panel comprising 29 B cell surface markers representative of the major gating schemes from 27 memory B cell studies, none of which used the same classification scheme. We then generated a cytometery dataset, replicated and mapped subset labels from 7 published gating schemes to each other and unsupervised clustering output.

**Results:** We found that different gating schemes produced inconsistent classifications for the same cell, and clusters from different gating methods with the same label had varying levels of similarity by JSI. Using the normalized variation of information metric, we find that the overall technical similarity between gating schemes varied from 0.16 (highly dissimilar) to 0.60 (moderately similar). We found that using JSI with unsupervised clustering, and further incorporating literature-based frameworks, can help disambiguate human splenic B cell clustering results, identifying subsets described in the literature, with additional intra-subset heterogeneity.

**Conclusions:** Cluster comparison metrics, combined with unsupervised clustering and in silico replication of published gating schema enables accurate cluster comparisons across studies, increasing rigor, reproducibility, and precision. Overall, these methods enabled detailed mapping of multiple literature-based B cell subsets onto a single classification map of human splenic B cells, harmonizing predefined definitions with in-depth immunophenotyping.

## 1 INTRODUCTION

A fundamental challenge in immunobiology is how to consistently classify cells based on biological markers. Such classification relies on a series of steps: feature specification, feature measurement, clustering of cells with similar features, and cluster label assignment. In cellular immunology, the correlates of this are marker selection (e.g., CD19, CD27, IgM), measurement of marker expression (e.g., flow cytometry, gene expression), manual (e.g., “gating”) or automated (e.g., machine learning) clustering, and phenotype label assignment (e.g., marginal zone B cells (MZBs), naive IgM+ B cells). This process reduces an *n*-dimensional feature vector of marker expression to a one-dimensional label. These labels are used throughout the literature as shorthand for a cell population and its associated functional properties (e.g., activation potential, immune memory, gene expression). Further, these labels are incorporated into experimental protocols, used to sort cell populations, and to create knowledge frameworks. A critical and pervasive issue across the literature is that investigators use different marker panels and clustering methods, but assign the resulting cells the same label. In some areas, like B cell immunobiology, this has led to babelization, where reports refer to population labels (e.g., MZBs) but the underlying cell populations cannot be mapped across studies using the same label. This is a profound issue for rigor and reproducibility, especially within the field of immunology, where flow cytometry is a foundational method for cell phenotyping, subset identification, and mapping populations to working ontologies.

The motivating case for this work is comprehensive memory B cell (MBC) flow cytometry phenotyping and mapping of MBC subsets in the human spleen. The spleen is the largest secondary lymphoid organ, maintaining a unique reservoir of *∼*1.2-2.5 x 10^9^ MBCs derived from *de novo* immune responses in the spleen or trafficking from other lymphoid tissues via the blood. (1, 2) The spleen also maintains populations, such as MZBs, that develop within the unique splenic microenvironment. (3) Much of the MBC literature, however, has relied on a wide variety of disparate flow cytometry phenotyping panels and markers applied to peripheral blood. Given the wide variety of splenic B cell responses and subsets, we hypothesize that the spleen retains phenotypes with a broader and richer range of differentiation states than those found in peripheral blood. Thus, to fully understand the development and persistence of B cell memory, it is critical to align phenotypic subsets across peripheral blood and spleen, and across the myriad of published technical phenotyping workflows.

We currently lack methods that enable rigorous cross-comparison of B cell clusters and their assigned labels across the literature. Many of the discrepancies in MBC subset phenotyping and classification in the literature stem from wide variations in flow cytometry panels, gating strategies, and analytical techniques, as well as a lack of standardized methods. This is a major issue in MBC immunology, since the same subset labels often refer to phenotypically heterogeneous cell populations. Indeed, when we reviewed 27 studies across the literature, we found that none use identical surface marker panels, flow cytometry gating, clustering strategies, or a standard ontology for cluster naming **(Supplemental Data 1)**. This issue of inconsistent classification goes beyond an exercise in ontology, as subset definitions are used to sort cells to test function, gene expression patterns, B cell receptor specificity, and repertoire distribution across memory compartments. Yet comparative studies using the same cell population with different gating or clustering strategies are almost entirely lacking.

Many attempts have been made to develop a unified B cell classification system, but none have been widely adopted, and we remain unable to consistently compare experimental results across studies. This is primarily due to an inability to determine precisely how cell clusters assigned the same label in two different studies correspond. The classical problem of cluster matching can be defined as follows: given two clusterings of the same cells obtained by two different methods (e.g., manual gating, unsupervised clustering), determine which clusters correspond to each other and to what extent. The degree of correspondence for clusters identified by each method can be calculated by generating a similarity matrix (e.g., Jaccard similarity index, overlap, F1 score) and comparing the membership of cells in each cluster in method A to each cluster in method B. (4) Unless both clustering approaches are constrained to result in the same number of clusters, there is no 1:1 correspondence to optimize. In addition, there is the biological question: how do clusters split or merge between published results or gating strategies, generally referred to by cluster labels (e.g., class-switched MBC, IgM+ MBC), which are used to infer cellular function, differentiation, and lineage.

Here, we describe a comprehensive approach to de-babelizing MBC phenotypic subsets identified using different manual gating or automated clustering methods, surface marker panels, and categorical labels (e.g., MZB). To this end, we have developed an extensive surface marker panel encompassing markers from 7 published studies on MBCs, as well as a broadly applicable computational analysis pipeline for human splenic B cell phenotyping and clustering. Finally, we implemented methods for cross-study subset comparison, label matching, and cluster disambiguation. This method provides a deeper understanding of the relationship between cell populations in experimental results across the literature.

Our approach mapped MBC phenotypes across multiple publications, revealing a one-to-many relationship among B cell clusters assigned the same label (e.g., MZB, double-negative (DN) B cell). Our analysis provides both an analytic pipeline for others to use and a coherent framework for identifying and mapping human splenic MBC phenotypes. Beyond de-bablization alone, we can now compare studies that have used differing phenotype definitions for hypothesis generation and to evolve the way we think about immune cell subset-specific phenotype and function.

## 2 RESULTS

We describe our results in four stages. First, we evaluate the motivating problem, the current babelization of MBC classification, with specific reference to human splenic MBC subsets. Second, we describe the analytic pipeline results, including mitigation of technical and analytic variation and methods for de-babelizing findings across multiple studies. We further describe the key finding that hierarchical gating, as applied in the literature, is not transitive; different hierarchical gating orders yield different populations that are often assigned the same phenotypic label. We then use the pipeline to map biologically relevant B cell subsets (e.g., MZBs, DN B cells) across multiple studies in the literature, and compare hierarchical manual gating with unsupervised clustering. Finally, we present a framework for integrating pipeline analytic results and cluster mapping with knowledge of B cell immunobiology to address the selection of the cluster number (i.e., *k*), assess biological context, and support hypothesis generation.

### 2.1 Motivating problem: Memory B cell subset classification variation

#### 2.1.1 Cross-study comparison is impeded by ambiguous MBC subset definitions and labels

We first evaluated marker usage and subset-naming conventions across 27 previously published studies (**Supplemental Figure 1; Supplemental Data 1**). (3, 5, 6, 7, 8, 9, 10, 11, 12, 13, 14, 15, 16, 17, 18, 19, 20, 21, 22, 23, 24, 25, 26, 27, 28, 29, 30, 31) We summarized these results in a rich factor bubble plot (**Figure 1A**), in which core marker usage and subset definitions were quantified across 27 studies, and expression patterns classified as positive, negative, or in disagreement. While phenotypic definitions of widely recognized populations, such as class-switched (CSW) MBCs, were largely similar, niche populations, such as atypical or MZBs, were more variable. Additionally, we found that even when not in direct disagreement, there was substantial variation in which core markers were used to define a subset (e.g., CSW MBC defined by IgD^-^ versus IgG/A^+^). For example, defining a DN label using the surface markers CD21 and CXCR5 versus CD21 and CD11c identifies partially overlapping, but not completely overlapping, subpopulations. In short, multiple different marker combinations have been used by different investigators to assign B cell subsets identical labels, without confirmation that the cell populations are the same. Critically, no study used exactly the same B cell classification, ontology, or surface markers, making direct comparisons virtually impossible.

**Figure 1.**
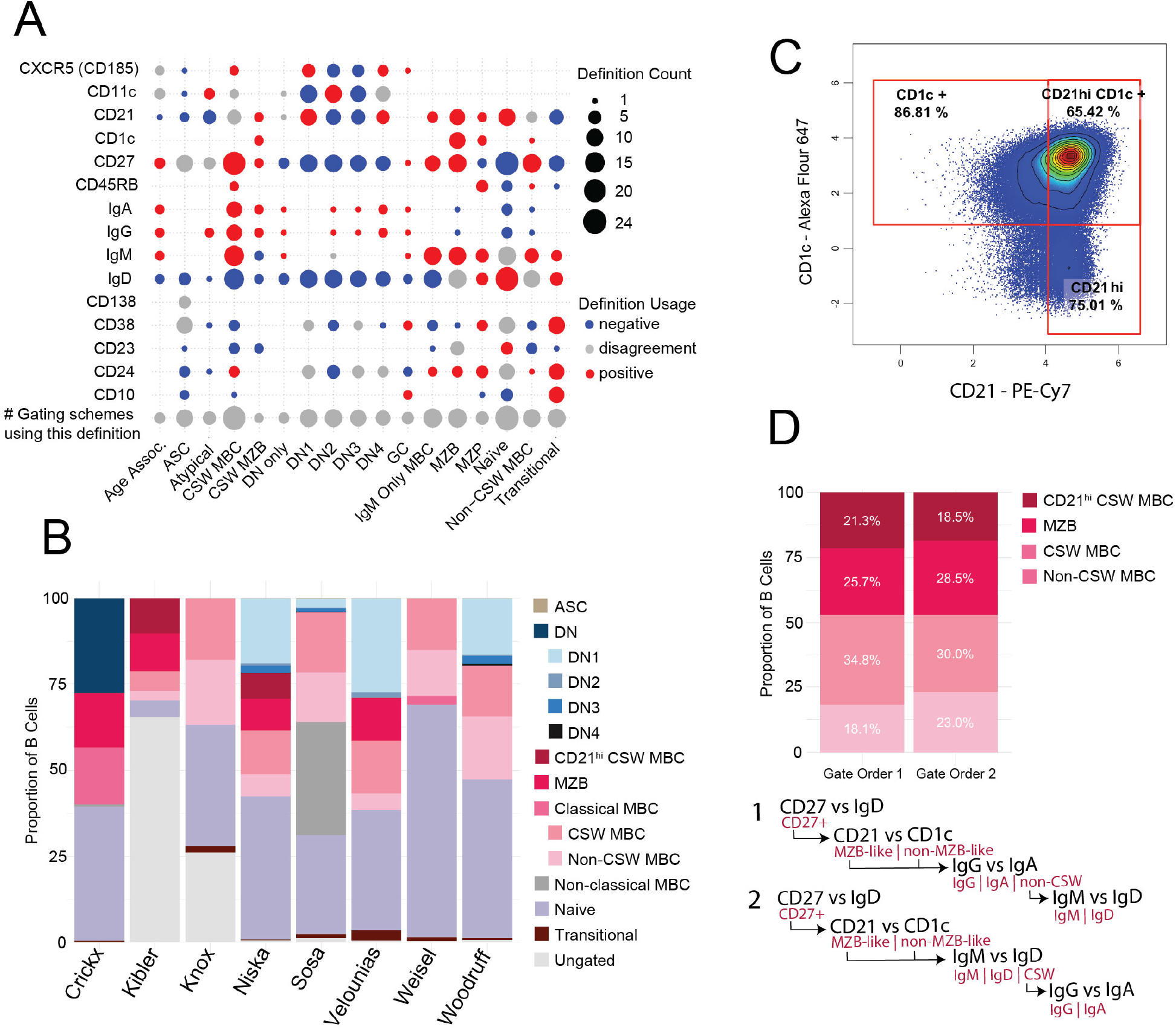
Variation in marker usage, subset definition, and gating results in cell phenotype assignment discrepancies. **(A)** Rich factor bubble plot comparing marker usage and subset definition across 27 studies’ gating schemes. **(B)** Proportion of cells from the same parent population assigned to each phenotype when gated across 8 different gating schemes (**Supplemental Figure 3)**. **(C)** Gating of total CD27^+^ IgM^+^ B cells using MZB definitions CD1c^+/hi^, CD21^hi^, or CD1c^+/hi^ CD21^hi^. **(D)** Proportions of cells identified as CD21^hi^CSW MBC, MZB, CSW MBC, or non-CSW MBC using the gating orders below, where black text indicates the plot axis and the pink indicates which population was gated (detailed in **Supplemental Figure 4**).

In addition to the problem of different marker definitions for similar subpopulations, two differently named subsets may actually contain overlapping cell populations that have been ascribed different functional relevance. For example, the subset age-associated B cells (CD11c/T-bet^hi^ CD27^+^) increase with age and have a phenotype overlapping with atypical B cells (CD11c/T-bet^hi^ CD27 ^-or^ ^+^) that are frequently associated with autoimmune diseases such as SLE, COVID-19 infection, or vaccination (**Figure 1A**). (32, 14, 15, 33) In these studies, CD11c/T-bet^hi^ CD27^+^ cells overlap phenotypically between the two subsets, and it is impossible to compare results across the two contexts because the terminology and associated phenotypes differ.

#### 2.1.2 Overlapping and inconsistent B cell classification arises when only select subsets are targeted for study

We next asked how the use of only select subsets impacted the resulting B cell classification. In **Figure 1A**, it is apparent that while most studies use common classifications like CSW MBC and naive B cells, many focus on a few select subsets instead of a comprehensive analysis of all subsets that may be present in a dataset. Thus, we wanted to determine whether disparities in subset usage led to differing classifications of the same cell. Briefly, we performed spectral flow cytometry on human splenic B cells using a panel of 29 surface markers (**Supplemental Table 1**). We then applied 7 published B cell gating schemes to the same cytometry data to compare the resulting B cell classification (**Supplemental Figures 2 and 3B-F**). (3, 15, 31, 34, 12, 28) Crucially, the panel included all markers used across the set of 7 published manuscripts, enabling us to replicate their gating strategies. In addition, we applied our own gating schema, which identified and mapped all splenic B cell populations to commonly used definitions (**Supplemental Figure 3A**). Gating was performed on the same B cell (CD19^+^ CD20^+^ CD3^-^ CD14^-^ CD16^-^) dataset and subset proportions compared across studies (**Figure 1B; Supplemental Figure 3**). Importantly, the greatest variation stemmed from a study’s intended focus: answering a specific research question directed towards a few MBC populations of interest, rather than identifying all MBC populations present. Studies with such a selective focus led to the grouping of individual B cells into different subsets with phenotypic and potentially functional differences (**Figure 1B**). For example, in studies that do not use the term DN (CD27^-^ IgD^-^), DN cells are often placed into the naive gate due to a lack of CD27. Furthermore, phenotypic classification of MBC populations in the human spleen is largely based on MBC subset definitions and gating strategies extrapolated from peripheral blood analyses, leading to the co-clustering of spleen-specific subsets, such as MZBs with classical MBCs.

#### 2.1.3 Lack of standardized gating schemes results in conflicting classification of the same cell

We next investigated how different gating strategies can lead to substantial variation in cell classification despite using the same cell surface marker phenotype definitions (**Figure 1B**). As expected, we found that the gate boundaries for markers with clear positive and negative populations showed a high degree of correspondence (**Supplemental Figure 2**). In contrast, gating schema using markers with broader biological distributions (e.g., broad or modest fluorescence signal peaks), such as those associated with signaling (e.g., FcRL5, CD85j), cell trafficking (e.g., CCR7, CXCR5), or differentiation state (e.g., CD21, CD69), often lacked clearly defined positive and negative populations, which confounded reproducibility of population assignments (**Supplemental Figure 2**).

Beyond marker separation, we investigated how variation in cell surface markers and gating schemes affected the classification of human splenic MZB subsets, a unique population residing within the splenic marginal zone (MZ) niche. Using the gating strategy from Kibler et al., MZBs are defined as CD21^hi^, whereas Velounias et al. gate on CD1c^+^ cells. (3, 30) While both authors recognize and show expression of the alternative marker within their gated population, each relies on only one marker for classification. We first gated CD27^+^ IgM^+^ B cells and then gated MZBs according to definitions used by Kibler et al. (CD21^hi^), Velounias et al. (CD1c^+^), or in combination (CD1c^+^ and CD21^hi^) (**Figure 1C**). We found 65.4% of cells overlap, both CD1c^+^ CD21^hi^, but when using only CD1c or CD21, an additional 21.4% (CD1c^+^) or 9.6% (CD21^hi^) of the total CD27^+^ IgM^+^ population was represented within the gate (**Figure 1C**). Consequently, while both studies evaluate MZBs, the results may not be generalizable between the two studies, since they were gated using differing criteria.

Finally, we evaluated how variation in the order of applying identical gates to the same cell population can affect the final proportion of a given subset. We began with pre-gated, non-transitional, non-ASCs, and then gated for MBCs (CD27^+^ CD21^-/+^) versus MZB cells (CD27^+^ CD21^hi^ CD1c^+^). These were separated into non-CSW and CSW MBCs using identical gates, but changing the sequence of the isotype gate **(Figure 1E**). The variance in CSW vs non-CSW between gate order 1 and 2 was 2.8% for the MZBs and 4.8% for the MBCs **(Figure 1E; Supplemental Figure 4**). The phenomenon is especially evident with co-expressed markers, here IgG and IgM, where the gating order can have a large impact on cell classification. This illustrates how disparate cell classification can arise and rapidly accumulate across the literature.

#### 2.1.4 A method to compare disjointed results across studies

While none of the above observations invalidate any given study, the above examples demonstrate how methodological variations can give rise to profound reproducibility issues. To our knowledge, this is the first rigorous study to demonstrate the considerable heterogeneity in MBC phenotyping and subset assignment across publications. We have demonstrated that using the same subset name with different definitions, or vice versa, results in different cell classifications. Importantly, by linking a subset label to function, disease- or age-related changes, or developmental pathways without identical phenotypic definitions, we risk misinterpreting results, highlighting the need for a rigorous method to compare classification results.

### 2.2 A rigorous and reproducible analytic pipeline for comparison of flow cytometry data

To comprehensively address reproducibility and classification issues with flow cytometry-based phenotyping, we developed a complete R-based pipeline that includes quality control, batch normalization, manual gating, clustering, and a toolbox of methods for data exploration and visualization **(Figure 2**). Additionally, we created a quantitative statistical tool for directly comparing two cell populations that have been gated or clustered, using an implementation of the Jaccard similarity index (JSI), that is broadly applicable across the field of immunology. We report here applications of our method (*see* Supplemental Material for R documentation) to a human splenic B cell flow cytometry dataset (see Methods) to 1) compare gating schemes for cell identification across studies and 2) demonstrate how we have implemented cross-study comparison to achieve better interpretation of machine learning clustering (MLC) results. Finally, we demonstrate how our method can provide insight into the phenotypic and functional implications of overlapping and diverging subset labels.

**Figure 2.**
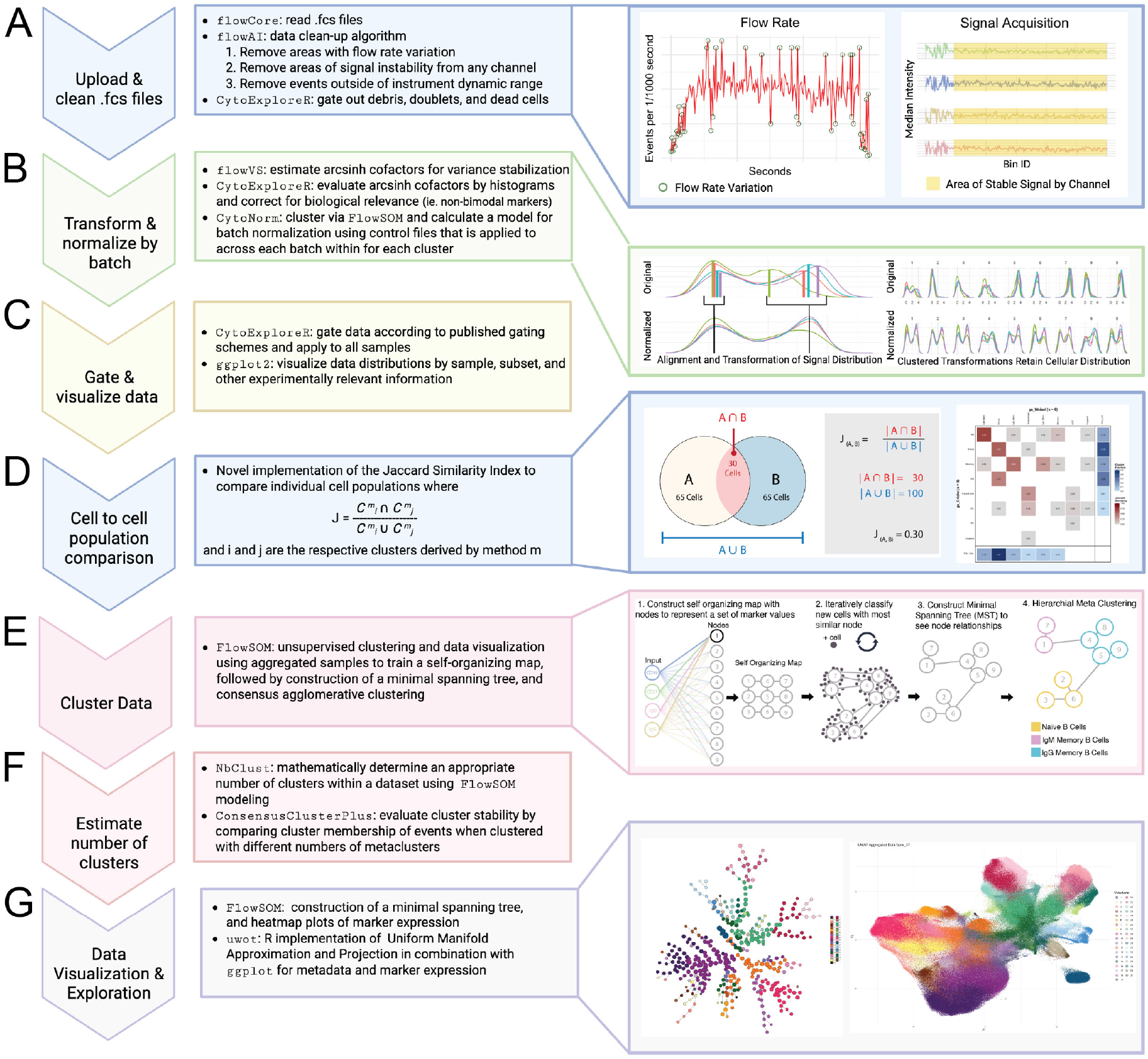
A robust R-based pipeline for rigorous analysis of flow cytometry data. Overviews of each step used in the flow cytometry pipeline, with R packages identified by `monospaced font` formatting. Primary steps include:**(A)** Initial uploading and quality control of .fcs files, **(B)** the transformation and batch normalization, **(C)** manual gating with visualization, **(D)** JSI based cell subsets comparison, **(E)** a summary of the `FlowSOM` algorithm steps, broken into substeps 1-4, **(F)** methodologies for estimating the number of clusters, and **(G)** visualization methods for data exploration.

#### 2.2.1 Data quality control

MLCs are often extremely sensitive to data anomalies, which can skew fluorescent shifts in batch normalization and lead to clustering driven by irrelevant outlier data points rather than biologically significant heterogeneity. Prior to analysis, we performed multiple quality-control checks to ensure the data was free of anomalies and artifacts that could cause downstream issues affecting the overall results. We used the R package `flowAI` to remove three types of common cytometry errors, including 1) variation in the cytometer’s flow rate, 2) signal instability, and 3) events above or below the instrument’s dynamic range **(Figure 2A**). (35) Lastly, data were gated using the `CytoExploreR` package to remove debris, doublets, and dead cells (**Supplemental Figure 2A**). (36) We found that performing these quality control steps significantly improved the results of algorithms used for batch normalization and clustering.

#### 2.2.2 Transformation and batch normalization

We next transformed the data using the inverse hyperbolic sine (*asinh*), which performs well in meta-clustering, with low intra-cluster variability and high inter-cluster variability. (37) Next, we determined the optimal cofactor for the transformation calculations for each fluorescent channel using Bartlett’s likelihood-ratio test to reduce variance within each channel, as implemented in the `flowVS` algorithm **(Figure 2B**). (37) To ensure optimal separation of each marker fluorescence, we then evaluated each cofactor using 2D pseudocolor plots and histograms of all live cells and pre-gated B cells to view the fluorescence distribution (**Supplemental Figure 5**). We found that while the algorithm is ideal for markers with bimodal distributions (e.g., CD3, CD19), further adjustment was required for markers with continuous or skewed unimodal distributions (e.g., CD21 or CD69; **see Methods R documentation**).

Batch normalization was performed using `CytoNorm` to allow for comparison and clustering across batches. The function uses a `FlowSOM` clustering step prior to normalization and normalizes within clusters, thereby preserving biologically relevant heterogeneity. (38) We first clustered the control samples using `FlowSOM` with cluster number values (*k*) ranging from 3 to 25 to determine the maximum number of clusters without clustering on batch effects **(Supplemental Figure 6**). The largest *k* value that kept the coefficient of variation CV*≤*1.75 across clusters was selected. Using the control samples, cluster-specific quartile distributions were calculated for each channel and aligned across batches, removing batch-specific variation while maintaining within-cluster variation. These were then applied to the entire dataset (**Figur**e 2B; Supplemental Figure 7).

#### 2.2.3 Gating and cell cluster comparisons

To compare MBC subsets as described across the literature, we pre-gated the live, single B cells (CD3^-^ CD14^-^ CD16^-^ CD19^+^ CD20^+^) and used the same dataset to replicate 7 gating schemes from the literature, and an additional gating scheme from the comprehensive staining panel (**Supplemental Figure 3**). The `CytoExploreR` R package provided a graphical user interface with flexibility to accommodate multiple manual gating schemes **(Figure 2C**). (36) Gating datasets in the R environment, we were then able to directly extract the gate assigned to each individual cell for each method for downstream analysis.

We analyzed cluster similarity between different gating schemes using JSI (see Methods). This method depends on applying two different clustering methods *m* to *the same primary flow cytometry data set*. Within each manual gating or unsupervised clustering schema, each cell was assigned membership in a cluster. For any two methods, we then compared the cluster assignments for all cells using the JSI **(Figure 2D**). If the clusters are identical *JSI* = 1 and if they have no cells in common *JSI* = 0. Thus, the JSI is the percentage of common cells between the two clusters relative to the total number of unique cells when A and B are combined. The resulting output is displayed as a heatmap, with each cluster pair showing a JSI between 0 and 1. Similarity between clustering methods, taking into account all cluster assignments, was assessed using the normalized variation of information (NVI; see Methods and Supplemental Material).

#### 2.2.4 FlowSOM clustering

While manual gating is a useful and arguably necessary tool for evaluating flow cytometry data, it is susceptible to human bias and highly dependent on the gating scheme and gate sequence used in two dimensions. Computational clustering algorithms are necessary to assess the richness of high-dimensional datasets, as they can look beyond 2D gating and analyze patterns of similarity within the data across all markers simultaneously. We have used the hierarchical `FlowSOM` clustering algorithm. Briefly, from a representative aggregate file, a grid is generated with a user-defined size (*n* x *y*), resulting in *n ∗ y* nodes that represent points in *d*-dimensional space, where *d* is the number of fluorescent parameters used in the clustering **(Figure 2E-1**). (39) Through an iterative process, new cells are assigned to phenotypically similar nodes. Upon each cell addition, the grid is reorganized so that similar nodes are placed closer together, known as a self-organizing map (SOM) **(Figure 2E-2**). (39) Once all cells have been assigned to a node, a minimal spanning tree (MST) is constructed to define the nodal relationships in the SOM. (39) The Euclidean distance formula was used to connect nodes via the shortest path, as nodes closer together are assumed to be more similar **(Figure 2E-3)**. Finally, a hierarchical meta-clustering step groups nodes into *k* clusters as defined by the user **(Figure 2E-4**).

Specifying the number of clusters that a data set partitions into is a fundamental issue for machine learning algorithms. For analyses using unsupervised algorithms, we rigorously determined the number of clusters (*k*) in the dataset using a consensus-based algorithmic approach. We first used the package `NbClust` which uses 30 different methods of cluster approximation to return a list of candidate *k*s selected by 1 or more methods to test *k* values from 5 to 50 **(Figure 2F**). (40) We then evaluated the resulting candidate *k*s to evaluate cluster stability using the `ConsensusClusterPlus` package **(Figure 2F)**. (41) Finally, we used the JSI to both identify the gates that a given `FlowSOM` cluster mapped to and determine how metaclusters with increasing *k* values map to one another. Final *k* values were chosen based on a combination of stability and biologically relevant population identification. These workflows are discussed in further detail in the following sections, with detailed code available in the **Supplemental Method R documentation**.

#### 2.2.5 Data visualization and cluster exploration

To compare marker fluorescence intensity distributions across metaclusters, we projected individual marker expression in a heatmap format onto the nodes of the MST to visualize within-cluster heterogeneity **(Figure 2G; Supplemental Method R documentation)**. For visualization, clustering results were projected onto a UMAP using the `uwot` package **(Figure 2G**). (42) UMAP with the Euclidean distance measure has the advantage of preserving global data relationship structure; similar clusters are closer together in the projection, facilitating trajectory analysis.

### 2.3 Mapping manual gating to unsupervised clustering improves rigor and reproducibility

Increased depth (i.e., number of markers) and complexity (i.e., understanding of developmental stages) in B cell phenotyping with flow cytometry has greatly broadened our understanding of the human B cell response. However, diverging phenotypic definitions and gating strategies lead to contradictory B cell classifications. Publications using a multitude of different markers and manual gating schemes designed to focus on very specific subsets have made it difficult to determine whether two studies using the same label for their clusters are actually discussing identical phenotypic cell subsets. Therefore, before considering phenotypic associations with B cell function, differentiation state, age, or disease, it is crucial to map the populations identified across different studies to one another. To this end, we used the previously described B cell dataset, with gating schemes replicated from the literature, and applied the JSI to directly identify biologically relevant points of convergence and divergence in B cell classification (**Figure 3; Supplemental Figure 3**).

#### 2.3.1 B cell clusters identified using the same label but derived using different markers and manual gating are not identical

First, we compared classification of MZBs using two published gating schemes defining MZB as either CD21^hi^ B cells (Kibler et al.) or IgM^+^ CD1c^+^ (Velounias et al.) (**Figure 1C and 3A-B**). (3, 30, 5, 43) Human MZBs have classically been described as a specialized MBC that resides in the splenic MZ, with a microenvironment-induced phenotype and a role in bacterial immunity. (1, 3) Kibler et al. describe the MZBs as a component of the spleen’s MBC archive (3), while Velounias et al. define two distinct populations of MZB distinguished by CCR7 expression. (30) CCR7^-^ MZB-1 are derived from a distinct IgM^hi^ marginal zone precursor (MZP) population, thought to be educated in the gut and biased towards anti-microbial immunity **(Figure 3B**). (5, 30) CCR7^+^ MZB-2 are thought to have a MZB phenotype that is induced by DLL1-NOTCH2 signaling within the splenic MZ, biased towards an anti-viral memory. (5, 30) This distinction has important implications for functional assays. Here, two studies have evaluated MZBs using different marker definitions; thus, the anti-bacterial or anti-viral MZB function inferred from the term MZB may actually be functionally heterogeneous.

To understand the overlap of these populations, we replicated each author’s gating and used the JSI to compare. The JSI is presented in a matrix format, comparing each defined subset within a given gating scheme, where a value closer to 1 (darker red) indicates greater similarity between the two gates **(Figure 3C**). We found that the 64% of the of the IgM MZB (sMZ) cells defined by Kibler et al., were grouped into Velounias’s MZB-2 (J = 0.45) or MZB-1 (J = 0.19), with a minority falling into the IgM-only MBC (J = 0.10) or non-CSW MBC (J = 0.05; **Figure 3C**). Alternatively, of Velounias et al.’s MZB-1 and MZB-2, a proportion fell into Kibler’s non-CSW MBC gate (J = 0.09 and 0.04; **Figure 3C**). This could result in substantial variability in any downstream functional assay readout. For example, Kibler et al. sorted CD21^hi^ cells and found that upon T-independent and T-dependent stimulation, MZBs had higher expression of activation markers and more proliferation than MBCs. (3) Based on the JSI results, we could hypothesize that the more anti-microbial-biased MZB-1s would respond to T-independent stimulation, while the anti-viral-biased MZB-2 would respond to T-dependent stimulation. Changes in the proportions of MZBs and MZB subsets across different panels or subjects would yield different results.

We next evaluated the population overlap between two gating schemes designed to identify different MBC subsets. Focused phenotyping panels omit markers for B cell subsets outside the study’s main focus. This may lead to increased heterogeneity within the subsets of interest, as other phenotypes become invisible due to a lack of markers. For this analysis, we compared the gating strategies of two studies, Velounias et al. and Woodruff et al., both of which focus on MBCs. Velounias et al. focused on the MZB subset and Woodruff et al. focused on the DN MBCs **(Figure 3B, D**). (30, 15) Each used a subset of MBC markers specific to their phenotype of interest. We replicated the gating strategies on the same dataset **(Figure 3E**). We found the JSI score was high for IgM MBCs, naïve, DN1, and CSW MBCs. As expected, the phenotypes labeled MZPs and MZBs by Velounias et al. are both included with the labeled naïve and non-CSW MBC populations of Woodruff et al. **(Figure 3E**). Though MZBs are associated with the spleen, Velounias et al. suggest they are also a circulating subset that downregulate CD21, while maintaining CD1c expression, meaning MZBs, specifically anti-viral biased MZB-2s, could be important in Woodruff et al.’s analysis of the peripheral blood response to COVID-19. (30, 15, 5) However, the gating strategy of Woodruff et al. did not separate these two populations or measure how their proportions varied across subjects. While Woodruff et al.’s primary focus was on the pathogenic role of DNs in severe COVID-19, a deeper analysis of anti-viral-like MZB-2s may have allowed investigation of their potential role in less severe COVID-19.

**Figure 3.**
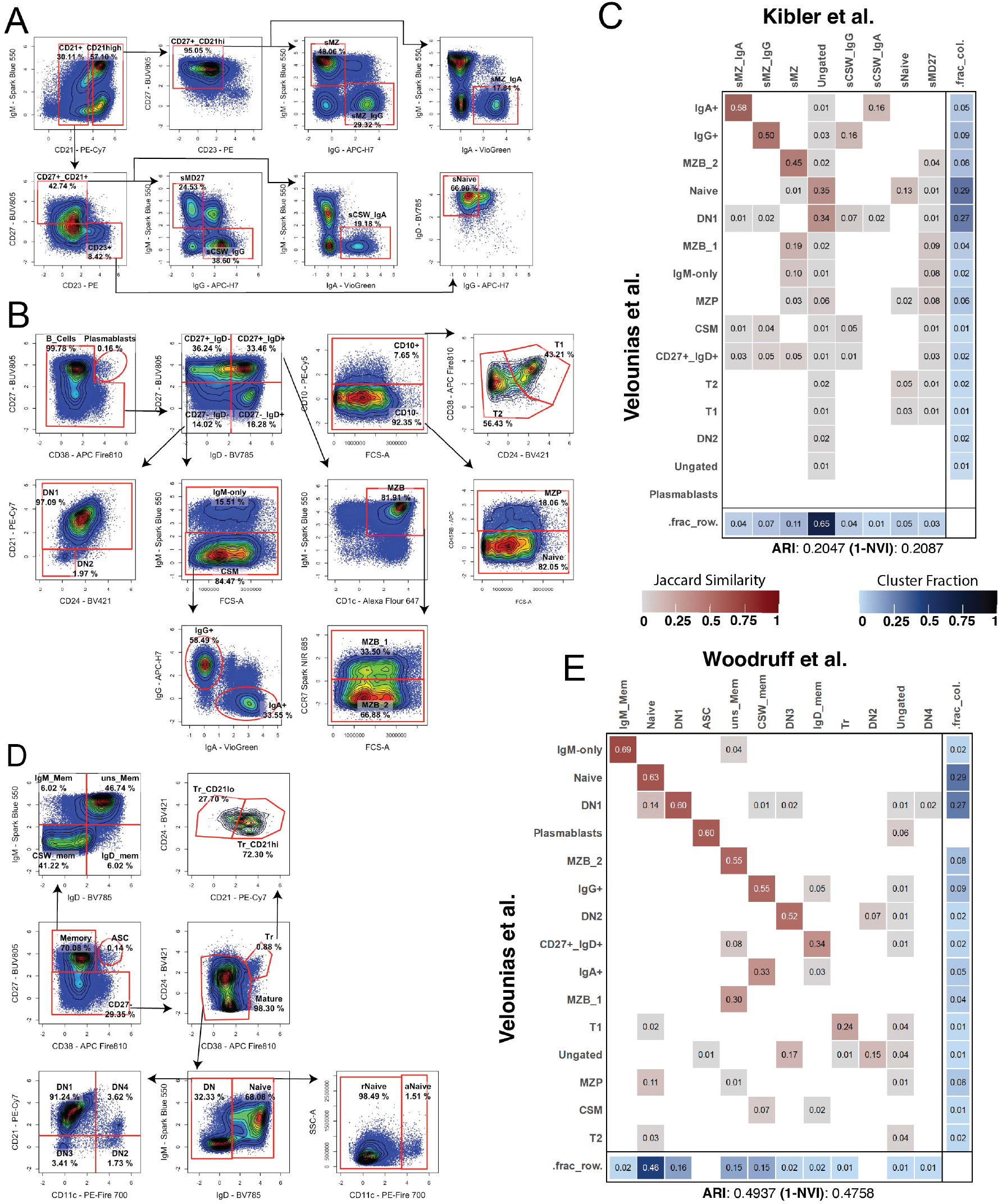
Comparison of marginal zone B cell populations classified by two different studies. Human splenocytes were isolated and stained with a 29 marker flow cytometry panel. This allowed direct and rigorous comparison of cluster membership assignment using different published gating schemes for marginal zone B cells (MZB). **(A)** Manual gating schema of Kibler et al., for clustering MZB. **(B)** Gating schema of Velounias, et al. for manually clustering MZB, including separation of MZB-1 and MZB-2 subpopulations. **(C)** Heatmap of cluster membership comparison using the Jaccard Similarity Index. Cells contain JSI scores for pairwise comparisons between gating methods on a red gradient, with blank areas indicating JSI *<* 0.01. The blue row and column indicate the percentage of cells in each cluster by method. For example, the DN1 cluster identified by Velounias’s gating comprises 27% of the total cells and shares cells with the two clusters of Kibler’s gating (Ungated and Naive). **(D)** Gating schema of Woodruff, et al. focused on manually clustered MBC and DN populations. **(E)** JSI mapping of cluster membership when either the Velounias or Woodruff gating schemes were applied to the same cytometry data. Note the high correspondence of the ASC (Woodruff) and plasmablast (Velounias) populations.

These two reports also provide further examples of how the same labels (e.g., DN1, DN2) were applied to cell clusters derived from different markers and gating strategies. Woodruff et al. defined DN1 as CD21^+^CD11c^-^, whereas Velounias et al. defined DN1 as CD21^+^ CD24^+^ and DN2 as CD21^-^ CD11c^-^ (Woodruff) versus CD21^-^ CD24^+^ (Velounias) (**Figure 3B, D, E**). (15, 30) Interestingly, discussion of the DN1 and DN2 MBC clusters assumed that they were identical across the literature. Our analysis showed that Velounias et al. did not include markers for the less frequently studied DN3 or DN4 subsets, which were then assigned to the DN1 and DN2 clusters by Woodruff et al. This is clearly visible in our comparative analysis using the JSI score (**Figure 3B, D, E**).

These are not isolated examples. We performed pairwise comparison of 8 gating and clustering schemes in the MBC literature using the JSI to assess overlap in membership of each identified (**Supplemental Figure 8**). These results demonstrated wide variability in cluster overlap between published investigations, making it difficult to compare results across the literature. We next used a modified version of the normalized variation of information (NVI) metric to assess how well the methods compared across all identified clusters (**Table 1**). The metric 1-NVI = 1 if all the clusters identified between two methods are identical, and *<*1.0 for lesser degrees of cluster agreement. For these 8 studies, (1-NVI) ranged from 0.16 - 0.60. Of the studies with high agreement (1 *− NV I ≤* 0.5), all but one used gating for MBC and DN cells in their panels. In contrast, Kibler (3) used a gating rubric that split CSW MBC into MBC and MZB and had a low NVI range (0.183 - 0.292), unlike all other studies that did not. These results demonstrate that the specific phenotype focus of a study (e.g. MZB, DN), and thus the surface marker panel composition and sample origin (e.g. spleen, blood) matters for intra-study subset comparison.

**Table 1.** Pairwise similarity of published B-cell phenotyping classifications, quantified as 1 *−* NVI (normalized variation of information). Higher values indicate greater similarity. Dashes denote self-comparisons or pairs not repeated in the upper-triangular matrix.

| **B-cell populations focus** | **Study** | **Niska** | **Kibler** | **Velounias** | **Knox** | **Sosa** | **Woodruff** | **Crickx** | **Weisel** |
|---|---|---|---|---|---|---|---|---|---|
| Broad spleen | Niska | — | 0.2629 | 0.4147 | 0.4343 | 0.3825 | 0.4504 | 0.3993 | 0.3508 |
| MZB + CSW MZB | Kibler | — | — | 0.2087 | 0.2230 | 0.1588 | 0.2061 | 0.1831 | 0.2924 |
| DN 1/2 + IgM MZB 1/2 | Velounias | — | — | — | 0.6041 | 0.4520 | 0.4758 | 0.5390 | 0.3925 |
| Tr + MBC | Knox | — | — | — | — | 0.4043 | 0.5303 | 0.5524 | 0.3822 |
| DN 1-4, MBC, Tr, ASC | Sosa | — | — | — | — | — | 0.5043 | 0.5595 | 0.3265 |
| Tr, DN 1-4, MBC | Woodruff | — | — | — | — | — | — | 0.5531 | 0.3937 |
| MZB, MBC, GC, PC, DN (no subset) | Crickx | — | — | — | — | — | — | — | 0.4080 |
| Tr, MBC, ASC | Weisel | — | — | — | — | — | — | — | — |

#### 2.3.2 Unsupervised clustering minimizes bias from surface marker and manual gating

To remove human bias in gate placement, subset selection, and surface marker usage, many studies now use automated clustering algorithms. The greatest hurdle is determining the number of clusters (*k*), as there is no defined ground truth or standard convention. Cluster identification is a key step in both evaluating final clustering results and determining an appropriate number of clusters by identifying where further subgrouping becomes biologically irrelevant **(Figure 2D)**. Thus, we will first discuss how we identified clusters and then how we determined the number of clusters to use for this analysis. We have used the JSI to aid in cluster identification by mapping clusters to multiple gating schemes to evaluate cluster composition. For this example, we will use *k* = 14, intentionally under-clustering the data, which resulted in clusters with intra-cluster heterogeneity.

We clustered an aggregated dataset comprising 9,000 cells from each FCS file, across all batches, with more than 9,000 B cells to ensure inclusion of all relevant populations. The FlowSOM model was trained on a 20 x 20 grid, yielding 400 unique nodes, which were grouped into 14 metaclusters **(Figure 2E**). In our initial analysis, we used the previously described gating schemes **(Figure 3; Supplemental Figure 3)** to map each cluster to the previously defined populations. The comparison of multiple gating schemes versus a single clustering result using the JSI further illustrated the inherent bias in cell classification when restricted to a pre-defined manual gating scheme and how unsupervised machine learning clustering can improve upon gating-defined classifications **(Figure 4A-B; Supplemental Figure 9**).

Next, we compared two clustering schemes to the unsupervised clustering one that includes a phenotyping marker panel that broadly covers all B cell subsets described in the literatu re (Niska) and a manual gating scheme focused on classical and DN MBCs (Woodruff et al.) to illustrate how clustering can help overcome gating-induced bias (**Supplemental Figure 3A and D**). Using Niska vs *k* = 14, we can see that non-CSW MZB has JSI = 0.49 with cluster 5 and a JSI = 0.21 with non-CSW MBCs **(Figure 4A**). A high *JSI* value in 2 clusters suggests under-clustering, as the markers used to define gating of the non-CSW MBC and MZB subsets did not separate as expected based on manual gating. Of the 3 other gating schemes that included MZBs, each had a JSI between 0.51 and 0.64 for cluster 5, validating this cluster contains MZBs, but did not perfectly match any other gating strategy (**Supplemental Figure 9**). Alternatively, using Woodruff’s scheme (no MZB), cluster 5 is largely composed of non-CSW MBCs and IgM-only MBCs **(Figure 4B**). Importantly, even without Woodruff’s inclusion of MZB in the gating scheme, a MZB population was identified. This emphasizes the value of using clustering over manual gating, as it identified subsets that may not have been initially evaluated, signaling possible biologically relevant heterogeneity within a manually defined subset.

We then analyzed clusters based on surface marker expression to better understand the key components defining each cluster. Using FlowSOM, we extracted the minimal spanning tree to examine nodal cluster assignments with mean marker expression by node (**Figure 4C-F**). We found that the MST maintains biologically relevant structure and allows for clear separation between core markers, such as IgD/M in the bottom branches, while IgG/A make up the upper branches of the tree (**Figure 4D**). There was a concentration of the memory marker CD27 and, to a lesser extent, CD45RB in the outer branches, primarily on the right side of the tree, thereby identifying these portions as memory (**Figure 4E**). Subsets such as DN were also evident near the center of the tree, where there was no overlap of IgD and CD27, but clear expression of IgG/A (**Figure 4D-E**). Interestingly, while DN2 (cluster 13), DN3 (cluster 12), and DN4 (cluster 8) were separated into individual branches, cells with DN1 phenotypes (cluster 1) were mixed into a large, more naïve-like cluster (**Figure 4D**). This aligns with the hypothesis in the literature that the DN1 population functions as a precursor to the CSW MBC compartment. (27) Whereas manual gating allows identification of the DN1 and naive subsets, the clustering results and minimal spanning tree analysis revealed an intermediate population between the subsets, enabling a deeper evaluation of the biological processes.

Beyond the general structure, we also used the MST to examine phenotypic heterogeneity within a given cluster. For example, we have identified cluster 5 as broadly MZB-like, as it contains cells with high levels of CD21 and CD1c (**Figure 4D-F**). But if we look at the individual nodes within the cluster, while the majority were CD1c^hi^, only one of the 2 branches that make up cluster 5 was CD21^hi^ (**Figure 4F**). These findings reiterate the advantage of data evaluation beyond manual gating. Unsupervised clustering reveals biologically relevant heterogeneity that may be missed using a manual gating scheme. Additionally, evaluating all markers, not just select core markers for identification, allows assessment of the heterogeneity and states within a subset (detailed clustering information available in **Supplemental Figures 10-12)**.

**Figure 4.**
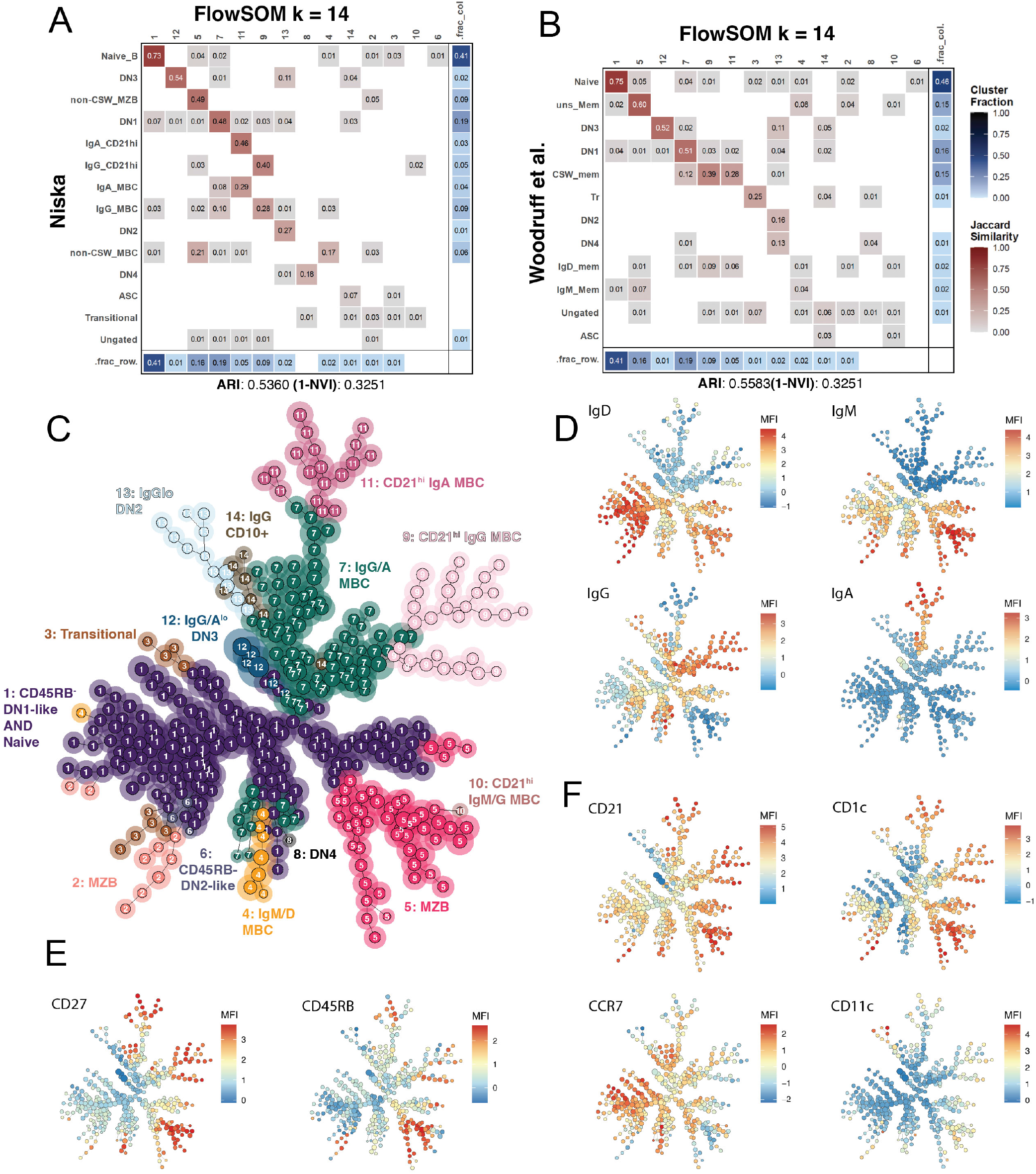
FlowSOM cluster identification can be assisted with the use of the JSI and MST nodal heatmaps. **(A)** JSI comparing FlowSOM clustering results for *k* = 14 and the Niska gating scheme. **(B)** JSI comparing FlowSOM clustering results for *k* = 14 and the Woodruff gating scheme. **(C)** MST with color-coded cluster identification, where each cluster node is identified by both the corresponding color and cluster number in white. **(D)** Heatmap visualization of the MST for immunoglobulin isotype. **(E)** Heatmap visualization of the MST for primary surface markers used to identify memory populations. **(F)** Heatmap visualization of the MST for primary surface markers used to differentiate MZB and DN subpopulations.

### 2.4 Incorporating prior biological knowledge improves cluster labeling and hypothesis generation combining mathematical evidence with biological insight

#### 2.4.1 Finding ***k***: Rational selection of cluster number using minimal spanning trees

We next investigated how MSTs, combined with biological context, could improve the selection of cluster number *k*. As a starting point, we used `NbClust`, which indicated the *k* values 5, 14, 24, 27, 31, 37, 43, and 50 qualified as the best choice in at least one of 30 indices tested, but statistics alone lack biological insight. To determine an appropriate *k*, we used MSTs to help classify clusters and identify where clusters branch into new subclusters, JSI to evaluate changes in cluster memberships, and a series of mathematical measures of cluster stability (**Figures 4 and 5; Supplemental Data 2**).

**Figure 5.**
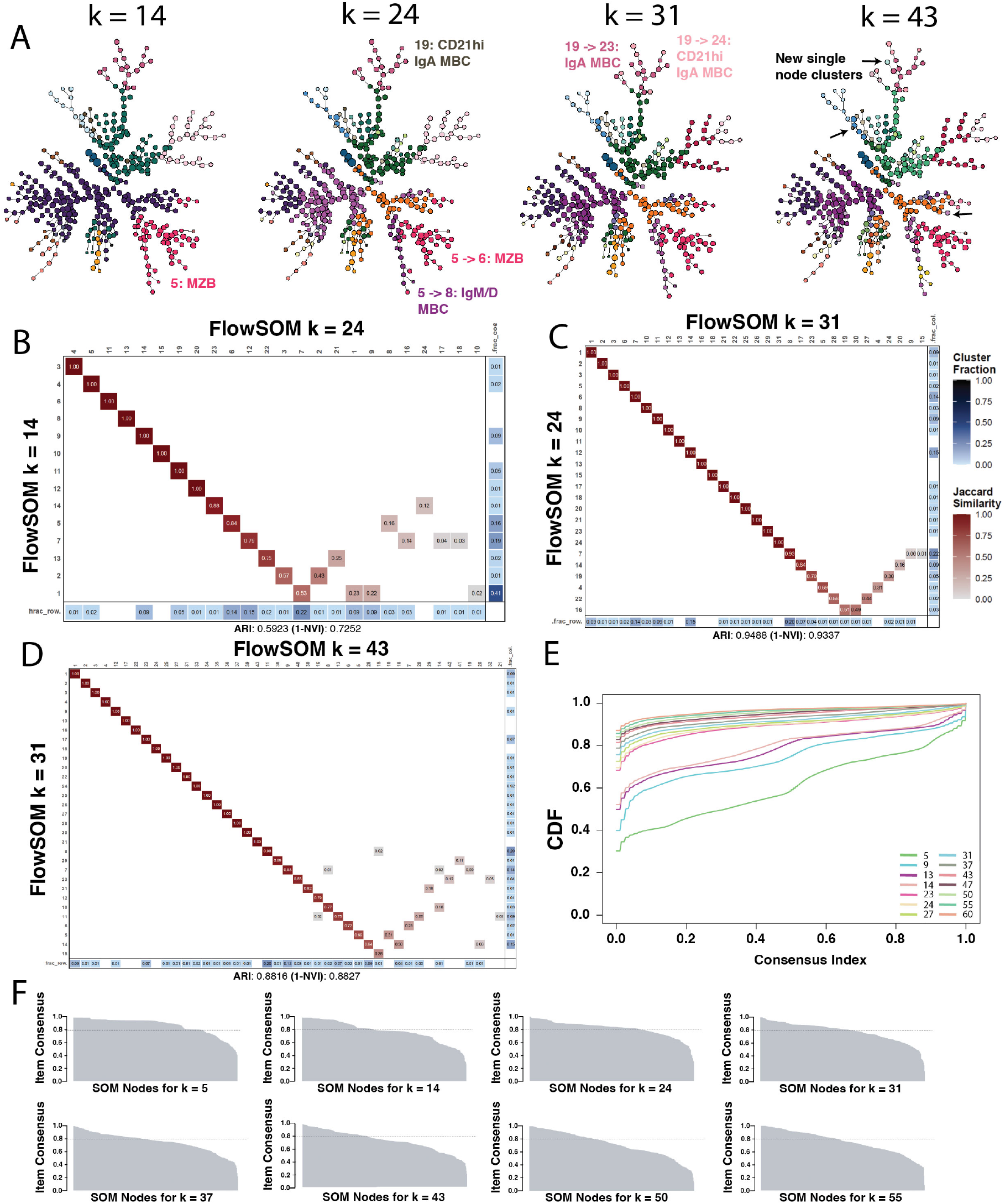
Identification of the best *k* value for clustering relies upon both biological context and mathematical determination of cluster stability. **(A**) MSTs at *k* values 14, 24, 31, and 43 with clusters color-coded to indicate splitting of clusters with increasing *k*. Key changes are annotated in the figure, including the original and new cluster numbers and classification. **(B)** JSI for comparison of clustering results for *k* = 14 and *k* = 24. **(C)** JSI for comparison of clustering results for *k* = 24 and *k* = 31. **(D)** JSI for comparison of clustering results for *k* = 31 and *k* = 43. **(E)** Concensus cummulative distribution function for clustering results at varying *k*s. **(F)** Item consensus scores for *k* values ranging from 5 to 55.

First, we visualized where a cluster split on the MST. For example, in **Figure 4F**, when *k* = 14, cluster 5 appeared heterogeneous, with a CD21^+^ MBC branch and a CD21^hi^ MZB branch. When we increased *k* from 14 to 24, the branches that made up cluster 5 were divided into 2 clusters (**Figure 5A**). Similarly, an increase from *k* = 24 to *k* = 31 resulted in splits of larger populations that we expect to see based on gating, such as IgA MBCs into a CD21^hi^ and CD21^+^ cluster. As we continued to increase *k*, we found that values beyond *k* = 37 began to split small clusters into multiple single-node clusters, representing very few cells **(Figure 5A**). While rare populations with smaller clusters exist, at high *k* values this is less likely and may reflect overclustering. At *k* = 14, we observed 2 clusters, each consisting of a single node, that persisted at every subsequent increase in *k*, representing rare subpopulations**(Figures 4C and 5A**). Of these, cluster 8 was DN4 B cells, which is expected to be very small, and cluster 10 was a MZB-like population with co-expression of IgD, IgM, and IgG **(Figures 4D and 5A**).

To evaluate cluster distribution at the cell level, we used the JSI to compare cell assignment at varying *k* values. In the top-left portion of the heatmaps, we observed that many cells remain within the same cluster (J = 1), whereas in the lower-right portion, clusters that were split into new clusters begin to fragment (**Figure 5B-D**). The cluster fraction on the x- and y-axis of JSI indicates the proportion of all cells represented by any given cluster. We found that at lower *k* values, further clustering often breaks up larger clusters. In the comparison of 14 vs 24 clusters, 14-1 is a large cluster representing 41% of B cells, but when *k* is increased to 21, it splits into 4 new clusters representing 22%, 9%, 9%, and 2% of all B cells **(Figure 5B**). A similar pattern is observed as *k* increases from 24 to 31 clusters **(Figure 5C**). When comparing *k* =31 vs *k* =43, many clusters from *k* = 31 are very small, representing only 1 to 2% of all cells, and are further divided into sub-clusters that represent less than 1% of cells at *k* = 43 (no value shown; **Figure 5D**). Further increases in *k* did not separate out novel phenotypic subsets; instead, they separated a small group of cells with changes in perhaps only one marker. While this could be useful for identifying subset phenotypic states within a larger cluster label, clustering beyond k = 37 did not reveal any novel subsets.

Next, we investigated clustering stability, how often two nodes are grouped in the same cluster, as a function of *k*. Using the `ConsensusClusterPlus` package, we calculated individual consensus scores for pairwise comparisons of 2 nodes, where 0 indicates never clustered together, and 1 indicates always clustered together. (41) The cumulative distribution function (CDF) was visualized over a range of *k* values. We found that *k ≤*14, a large proportion have low cluster stability scores (0.2-0.8; **Figure 5E**). Ideally, the score should be close to 0, indicating that nodes are either dissimilar and never assigned to the same cluster, or close to 1 when nodes are similar and always assigned to the same cluster. (41) The curve begins to flatten with *k ≤*27 approaching 0 at *k ≤*60, with most node consensus scores either 0 or 1 (**Figure 5E**).

Lastly, we evaluated item consensus (IC), the average probability that a node co-clusters with the same metacluster across iterations. This measure assesses cluster label stability as a function of *k*, where 0 indicates frequent changes in cluster assignment, and 1 indicates strong meta-cluster membership. (41) Overall, we saw that the IC will decrease as *k* increases, because separations between clusters are more granular and a given node may sit somewhat between two cell states **(Figure 5F**). Our analysis showed that the optimal value was *k* = 37 for this dataset, yielding a biologically balanced identification of primary subsets and intra-subset heterogeneity. Due to variability in sample processing, source, disease state, and different donors, *k* must be determined for each combined dataset to ensure separation of all unique populations and gain the desired granularity in intra-subset heterogeneity.

#### 2.4.2 Cluster label disambiguation within an evidence-based framework

We next applied our pipeline to address fundamental issues in splenic memory B cell immunobiology. This analysis used unsupervised, algorithmic clustering analysis for both data disambiguation and hypothesis generation. We used Uniform Manifold Approximation and Projection (UMAP) to visualize the findings at multiple levels of detail. UMAPs preserve both the local and global structure of the data; cells of the same subset tend to be near one another, often correlating with cell differentiation reflected by variation in feature (i.e., surface marker) expression levels.

First, we displayed the expression of surface immunoglobulin isotypes by overlaying a heatmap color gradient **(Figure 6A, all markers Supplemental Figure 13)**. Isotype is tied to B cell differentiation, and the map shows a clear progression of naïve IgD^hi^ IgM^lo/+^ in the bottom left, progressing upwards towards IgM^hi^ MZBs and IgM^+^ MBCs. Similarly, moving to the right on the UMAP, increased IgG and IgA with decreased IgM/D indicate CSW MBCs. We then displayed a simplified view of the main subsets described in the literature by merging clusters that were part of similar subsets (**Figure 6B**). This overlay mimicked the pathway inferred by isotype, with naïve cells towards the bottom-left corner, progressing upwards with MZBs and IgM/D memory cells, and to the right with CSW MBCs (**Figure 6B**).

**Figure 6.**
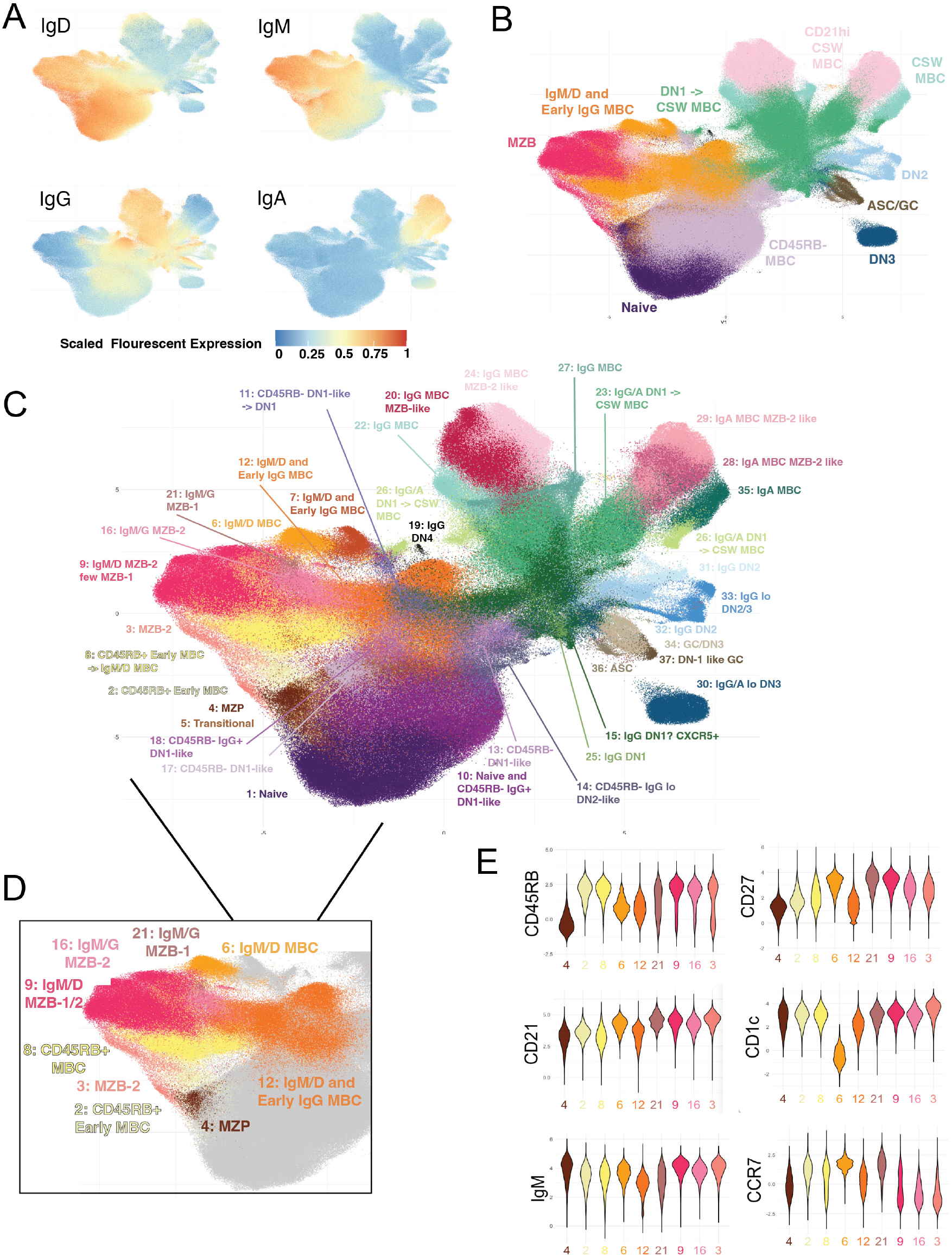
Heterogeneity with defined B cell subsets is resolved using high-dimensional clustering with FlowSOM and our proposed pipeline. **(A)** UMAP of splenic B cells with heatmap projections of B cell isotype expression. **(B)** UMAP with simplified clustering results, produced by merging all clusters that fell within a primary B cell subset. **(C)** Detailed UMAP with 37 color-coded clusters. **(D)** Magnification of UMAP with isolated MZB, IgM/D MBC, and MZP clusters. **(E)** Violin plots of isolated MZB, IgM/D MBC, and MZP clusters expression of CD45RB, CD27, CD21, CD1c, IgM, and CCR7.

##### Multidimensional unsupervised clustering reveals populations missed by manual gating

We next mapped all 37 clusters from unsupervised algorithmic clustering onto the UMAP, demonstrating substantial heterogeneity within the main B cell literature-defined subsets **(Figure 6C**). Since the human spleen is a lymphoid tissue capable of de novo B cell responses and a reservoir of B cells with unique microenvironments, we hypothesized that we would find greater heterogeneity than in the blood. Indeed, we observed many instances of clusters with phenotypes that had partial characteristics falling between two commonly defined subsets. This suggested that the human spleen may serve as a site of B cell differentiation, beyond its long-recognized function as a B cell repository. For example, in **Figure 6B**, the UMAP display shows a mixed population of DN1 and CSW MBCs in green. This larger cluster was composed of DN1 (clusters 15 and 25) and multiple clusters with phenotypes that suggested they were in transition from a DN1-like state to a CSW MBC state (clusters 23 and 26; **Figure 6C**). Our analysis identified a CD45RB^-^ subset (IgM/D^lo/+^ IgG^+^ CD45RB^-^ CD27^-^) that was not identified with manual gating. Comparing the unsupervised clustering and manual gating using Jaccard similarity revealed that these cells primarily fell into our naïve manual gate, a population we would have missed without unsupervised clustering (**Supplemental Figure 8**). Additionally, we found that among the CD45RB^-^ clusters, five (clusters 10, 11, 13, 17, and 18) had phenotypes similar to the DN1 population, and one (cluster 14) resembled the DN2 subset. This finding suggested that these clusters could be precursor populations undergoing downregulation of IgD (**Figure 6C**).

##### Identifying intra-subset heterogeneity

To evaluate intra-subset heterogeneity in greater depth, variation in marker expression is required. Here, we isolated clusters related to the MZP, MZB, and IgM/D MBC subsets, which were similar based on the global structure of the data and known B cell biology (**Figure 6D**). (3, 15, 31, 34, 12, 28) The MZP is an IgM^hi^ transitional subset, distinguished by lack of CD45RB or CD27 expression indicative of prior antigen experience, but elevated CD1c (**Figure 6E**). (6) The yellow clusters (clusters 2 and 8) were mapped **Supplemental Figure 1** as memory precursors and early memory, characterized by CD45RB^hi^, CD27^int^ expression. Interestingly, although not CD21^hi^, both clusters have CD1c levels similar to those of the MZP and MZB clusters (**Figure 6E**), suggesting the hypothesis that these may be intermediate stages requiring signaling within the MZ niche for differentiation into MZB or MBC (**Figure 6E**). The orange MBC clusters (clusters 6 and 12) were clearly separated from the MZB clusters and show differing expression patterns. Where cluster 6 is CD21^hi^ CD1c^-^ CD27^+^ MBC population and cluster 12 is CD21^+^ CD1c^+^ with an early memory-like phenotype CD45RB^+^ CD27^lo^ (**Figure 6E**). These hypotheses regarding intermediate MBC differentiation states in the spleen will need further experimental confirmation, but these examples demonstrate the power of quantitative unsupervised clustering and projected visualization for identifying previously unknown phenotypes in multi-dimensional space.

##### Hypothesis generation

Using these 6 markers (CD1c, CD24, CD27, CCR7, CD45RB, IgM/IgD), we were able to map each cluster to literature consensus labels. Importantly, however, the use of additional markers enhances hypothesis generation and identifies relevant markers for functional experiments. For example, clusters 16 and 3 appear nearly identical in **Figure 6E**, but cluster 3 is IgD^+^ CD24^-^, whereas cluster 16 is CD24^+^ IgD^+^, indicating that cluster 16 may represent a more mature MZB population (**Supplemental Figure 13 and 14**). Further, Hanson et al. have shown that in murine spleen, CD62L^-^ cells (cluster 3) are more recent and closer to the follicle and germinal center, whereas CD62L^+^ (cluster 16) reside in the MZ and have enhanced ability to differentiate into ASCs. (44) They also evaluated human PBMCs and found increased CD62L expression in SARS-CoV-2-specific cells at day 210 compared with day 35 post-vaccination. (44) These results, combined with our data, suggest important functional distinctions between these phenotypes relevant to further understanding the development of long-lived human B cell memory.

## 3 DISCUSSION

This study addresses an issue of rigor and reproducibility plaguing the field of flow cytometry-based phenotyping: the labels and definitions of cell subsets across the literature are inconsistent, overlapping, and were previously unresolvable. We have presented a broadly applicable method for de-babelization of subset definitions across the literature and have applied it to human splenic B cells. The pipeline we describe begins with quality control steps and progresses through each step of B cell classification, including manual gating, clustering, estimation of *k*, and visualization, with the addition of the JSI to compare gates and clusters at each step. While we initially developed this implementation of the JSI to compare cell populations using different gating schemes, we have demonstrated additional capabilities for evaluating the number of clusters, classifying B cell clusters, and generating hypotheses from cross-comparisons of data. While this method addresses inhomogeneity across broad categorical labels and improves upon the semi-subjective selection of *k*, it does not eliminate the need to select *k*. To date, despite multiple different metrics, selection of the number of clusters remains semi-subjective. If widely adopted, these methods could be pivotal for understanding B cell immunobiology at a systems level, using results from prior studies to evaluate the interconnected network underlying B cell development and memory persistence.

The advent of single-cell sequencing has been pivotal across the field of immunology, providing extensive detail for cell characterization and pathway prediction, but it is not a substitute for functional testing. Subset-specific functional testing requires cell sorting, based on surface marker expression. This is where the ability to compare subsets, as we have done with the JSI, becomes crucial. There is an abundance of functional testing data in the literature, but because subsets and clusters lack standard naming conventions and surface marker definitions, it is difficult to compare findings across studies. Thus, the methods of comparison we have presented here are necessary to understand areas of overlap and divergence to design future studies aimed at understanding B cell biology at a systems level.

Our analysis revealed substantial variation in B cell classification **(Figure 1B)**, for example, incomplete overlap between MZB population definitions in two seminal papers **(Figure 3C)**. (3, 30) These studies differed in marker definitions, methodologies, and objectives. Kibler et al. examined MZB/MBC phenotypes, responsiveness, and aging, proposing that circulating MBCs enter the spleen to form a systemic archive and transiently acquire a hyper-responsive MZB phenotype. Supporting this model, DLL1 stimulation can induce CD21 upregulation. (3) In contrast, Spencer’s group investigated MZBs in SLE, emphasizing gut-associated IgM^hi^ precursors linked to MZB-1 and antibacterial immunity. (30, 5) Both MZB subsets were reduced in SLE; reduced MZB-1 abundance was proposed to contribute to pneumococcal susceptibility. (30, 5) Although both studies inferred clonal relationships from BCR sequencing, neither confirmed receptor reactivity.

Future studies could compare CD1c^+^, CD21^hi^, and double-positive populations, subdivided into MZB-1 and MZB-2, using antigen-specific probes to assess reactivity and age-related accumulation across disease states. Our JSI results indicate overlapping definitions, but the relationship between CD1c^+^ CD21^+ (not^ ^hi)^ and CD21^hi^ CD1c^-^ populations remains unresolved, limiting the generalizability of single-marker studies. Distinct antiviral versus antibacterial reactivity profiles would further test the proposed models. Our method provides a basis for hypothesis generation and comparative studies that clarify MZB classification and function.

This study has provided insight into how variations in phenotypic descriptions of identically named subsets can easily lead to confusion when comparing results. Though the field has made immense progress in understanding the B cell response, to reach a systems-level understanding we must expand upon the findings presented here with comparative functional follow-up.This work uses both multi-dimensional clustering in high-dimensional space and two-dimensional projection for visualization. It is critical to note that the similarity of measured cell states in a two-dimensional projection (e.g., FlowSOM, UMAP) cannot be used on its own to infer MBC developmental trajectories. Notably, several different biological processes can produce the same visual continuum. These complementary methods can reveal heterogeneity obscured by conventional manual gating, identify populations worth sorting or profiling further, or detect co-regulation of surface marker expression. A future large, but worthwhile study using technologies such as CITE-seq could use these approaches to harmonize flow cytometry and sequencing results. The resulting reference dataset could be used to guide subset identification, sorting strategies, and methods of targeting particular genotypic profiles for functional assay in many future studies.

We have provided extensive R Markdown documentation of the entire pipeline and made the entire code accessible on Github. Additionally, because the entire pipeline is implemented in the open-source R environment, it can be reproduced and applied to any dataset. However, to perform this comparison, a reference dataset must exist in which all surface markers contained in the comparison studies are present. While this requires a large flow cytometry panel, once these datasets are available and made public, researchers can reuse them to evaluate data presented in the literature. If this method proves valuable, future endeavors may be undertaken to create an accessible web-based app that integrates multiple public datasets to enable quick comparisons using the JSI or other cluster membership comparison metrics. Lastly, although it is possible to evaluate different clusterings of the same dataset, we have not yet addressed the issue of comparing clustering results across studies, as these are not easily reproduced without the original dataset. To allow sorting or replication of clustering results across studies using 2D gating, algorithms such as `GateFinder` can be used to identify the best path to a clustered population with 2D gates. (45)

## 4 METHODS

### 4.1 Human Subjects Protection

This research study was a secondary analysis of existing de-identified samples from human spleen and peripheral blood samples, and was approved by the University of Rochester Research Subjects Review Board RSRB STUDY0000159.

### 4.2 Flow Cytometry Staining

Cryopreserved human splenocytes were thawed in warmed complete RPMI-1640 supplemented with 10% FCS, 55 µM 2-ME, 1% Anti-Anti, 10 mM HEPES, 1 mM Sodium Pyruvate, and 1% MEM NEAA and centrifuged at 300 × g for 10 minutes. Cells were then resuspended in 2 mL complete RPMI with 200 µL of DNase (1 mg/mL) and incubated for 15 minutes at room temperature. After incubation, 10 mL of complete RPMI with 5 mM EDTA was added to deactivate DNase and prevent clumping in downstream steps, and the mixture was centrifuged at 300 × g for 10 minutes. Cells were then resuspended in 3 mL of complete RPMI + EDTA, filtered through a 70-µm filter, and counted with Trypan Blue. Cells were brought to a concentration of 1e7 cells/mL.

For staining, cells were aliquoted at 200 µL per well into a v-bottom 96-well plate and centrifuged at 800 x g for 3 minutes at 4 °C. Media was carefully decanted, and cells were resuspended in 200 uL of staining buffer (1X PBS supplemented with 5 mM EDTA and 0.1% BSA and centrifuged at 800 x g for 3 minutes at 4 °C. All remaining steps were performed on ice. The staining buffer was decanted, and cells were resuspended in 50 µL staining buffer with 5% v/v normal mouse serum (Jackson ImmunoResearch Laboratories, 015-000-001 and incubated for 10 minutes. After incubation, 150 µL of staining buffer was added to each well, and the wells were centrifuged at 800 × g for 3 minutes at 4 °C. The wash step was then repeated with 200 µL of staining buffer. Cells were then stained with Live/Dead Blue diluted 1:200 in 50 µL of staining buffer for 20 minutes. After incubation, 150 µL of staining buffer was added to each well and centrifuged at 800 × g for 3 minutes at 4 °C. Cells were then resuspended in the antibody cocktail (see **Supplemental Table 2**) and incubated for 20 minutes. After incubation, 150 µL of staining buffer was added to each well and centrifuged at 800 × g for 3 minutes at 4 °C. Cells were then fixed with Cytofix/Cytoperm™ Fixation/Permeabilization Solution Kit (Becton Dickinson, 554714) according to the manufacturer’s protocol. Fixed cells were resuspended in 200 µL of staining buffer and kept at 4 °C until run on the Cytek Aurora 5-laser flow cytometer. Additionally, data were annotated with keywords in SpectraFlow for downstream R analysis using sample metadata.

### 4.3 Flow Cytometry Analytic Pipeline

Cells were acquired using a Cytek Aurora 5-laser flow cytometer at the University of Rochester flow cytometry core. Compensation controls were prepared using splenocytes, ArC™ Amine Reactive Compensation Bead Kit (Invitrogen, A10346), or UltraComp eBeads™ Plus Compensation Beads (Invitrogen, 01333341), by adding 1 µL of antibody to 1 drop of beads or 50 µL of cells, washed with 150 µL of staining buffer was added to each well and centrifuged at 800 × g for 3 minutes at 4 °C, and resuspended in 200 µL of staining buffer. Data were acquired and unmixed using SpectroFlo software.

#### 4.3.1 Data Pre-processing

Unmixed data was imported into R (Version 2026.01.0+392) using the `flowCore` package. (46) All files were cleaned for quality control using the package `flowAI`, after which new .fcs files containing only high-quality events were exported. (35) Using the package `CytoExploreR`, data were gated by batch to remove doublets and dead cells and exported as new .fcs files. (36)

#### 4.3.2 Transformation and Batch Normalization

After quality control, the cleaned .fcs files were imported, and a subsample from each batch were aggregated (3,900,000 total cells) to estimate the cofactor for transformation using the `flowVS` package. (37) Additionally, the aggregate file was transformed with the `transFlowVS` function with the cofactors 100, 200, 400, 600, 800, 1000, 2000, 3000, 4000, and 5000. The `flowVS` and manual transformations were compared using density plots with the `ggridges` package. (37) Cofactors were selected based on the expected marker distribution on density plots and with 2D pseudocolor flow plots (using the package `ggplot2`) for a list of commonly combined markers at the total splenocyte and B cell level.

Batch normalization was performed by aggregating the control files from each batch and applying the cofactor-optimized arcsinh transformation function. The aggregate files were then gated using `CytoExploreR` to remove any extreme values introduced by the transformation that negatively impact the overall normalization function calculation. (36) The files were then inverse transformed (`CytoNorm` uses untransformed data) to train the `CytoNorm` model. (38) Prior to training the model, the appropriate number of clusters to use for the `FlowSOM` aspect of the algorithm was selected using the `prepareFlowSOM` and `testCV` functions, and the highest *k* value (number of clusters) with a CV below 1.7 was selected. (39) Next, the `CytoNorm.train` function was used to train on the control samples run in each batch. The transformation function was then applied to the entire dataset and evaluated using the `CytoNorm plotDensities` function, with cofactors adjusted if visible issues were present in the density plot or in 2D pseudocolor plots for each marker by batch, to ensure similarity across batches.

#### 4.3.3 Manual Gating

Manual gating was performed on normalized, transformed data using the `CytoExploreR` package. (36) Manual gating for all gating schemes was performed on pre-gated to that excluded CD3, CD14, and, CD16 positive cells, and then gated on CD20 and CD19 positive cells. The gating schemes were all applied to this initial B cell dataset.

#### 4.3.4 Unsupervised Clustering

Clustering was performed using the `FlowSOM` algorithm. To determine the appropriate number of clusters for this dataset, we used the packages `NbClust` and `ConsensusClusterPlus`. (40, 41)Using all fluorescent channels, a `FlowSOM` SOM was generated with the parameters nClus = 5, seed = 27, silent = TRUE, xdim = 20, ydim = 20, rlen = 260, alpha = c(0.01, 0.001)). The SOM was extracted, and a set of 28 tests was performed with `NbClust` to determine candidate *k* values, using an initial *k* range of 5-50 and repeating with a variety of ranges to account for algorithms that consistently under- or overestimate the number of clusters. The stability of candidate *k*s was evaluated using the `ConsensusClusterPlus` package. Once selected, the final clustering was performed using `FlowSOM` with the parameters nClus = 37, seed = 27, silent = TRUE, xdim = 20, rlen = 260, alpha = c(0.01, 0.001)). The training dataset consisted of an aggregate file containing 9,000 cells from each sample with at least 9,000 B cells. The trained model was then applied to all samples.

#### 4.3.5 Data Visualization via UMAP

The UMAP was generated using the `uwot` package using all fluorescent channels with the parameters n neighbors = 50, min dist = 0.3, n components = 2, n trees = 100, seed = 27, ret model = TRUE using the aggregate dataset used for training the `FlowSOM` model. (42) The model was then applied to the entire dataset. UMAP results were exported into a `data.frame`. To project different data onto the UMAP, the gating and clustering results were extracted and stored in a `data.frame`, with each cell represented by a row containing columns for each fluorescent marker expression, clustering results, and gate assignment across all gating schemes. The resulting UMAPs were plotted using `ggplot2`.

### 4.4 Similarity measures

#### 4.4.1 Jaccard similarity index

Similarity between individual clusters was assessed using the Jaccard similarity index (JSI). Please note that this method depends on applying two different clustering methods *m* to *the same primary cytometry data set*. Within each manual gating or unsupervised clustering schema, each cell was assigned membership in a cluster *C_m,n_*, where *m* is the gating or clustering method and *n* is the cluster number. For any two methods, we then compared the cluster assignments for all cells, using the JSI to specify the overlap between clusters:

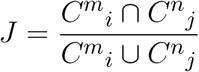

where *i* and *j* are the respective clusters derived by methods *m* (e.g., manual gating vs. self-organized maps), and 0 *≤ J ≤* 1 **(Figure 2D**). If the clusters are identical *J* = 1 and if they have no cells in common *J* = 0. Thus, the Jaccard index (J) is the percentage of common cells between the two clusters relative to the total number of unique cells when A and B are combined. The resulting output is a heatmap where each cluster has a J *{*0,1*}* displayed for each gating being compared.

#### 4.4.2 Normalized variation of information

Similarity measurement between clustering *methods* was assessed with the normalized variation of information (NVI) metric. NVI quantifies how much information is gained or lost when moving from one clustering method to another(47). It is especially useful when there is no ground truth labeling (e.g., the “true” number of cell subsets is unknown), and when different gating or clustering methods differ in the number or structure of the clusters. Raw VI can give values *≤ V I ≥ H*(*C*) + *H*(*C^′^*), where each entropy *H* depends on the number and size of clusters, with the resulting *V I >* 1. Changing the number of clusters or the dataset size changes the maximum possible VI. We used normalization to ensure that the metric is bounded between [0, 1] and interpretable. Values of 0 indicate that the resulting clusterings are identical between methods, while larger values indicate greater dissimilarity between all of the clusters generated by the two methods.

NVI between clusterings *C* and *C^′^* on *n* points (e.g., cells) is defined as:

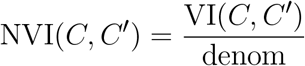

where the numerator is the variation of information:

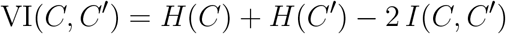

and the denominator denom depends on the chosen normalization method. We used a modified version (1-NVI) so it would scale similarly to JSI, 0 indicating no concordance between clustering methods being compared, and 1 indicating complete identity. The resulting (1-NVI) can be interpreted as:

- (1-NVI) *≤* 0.2: Substantial disagreement between clustering methods
- (1-NVI) 0.2–0.5: Moderate differences between clustering methods
- (1-NVI) *≥* 0.5: Very high agreement (over 80% of the combined structure is the same).

## CONFLICT OF INTEREST STATEMENT

The authors declare that the research was conducted in the absence of any commercial or financial relationships that could be construed as a potential conflict of interest.

## AUTHOR CONTRIBUTIONS

Conceptualization: [M.E.N., M.Y.S., M.S.Z.], Data curation: [M.E.N., M.S.Z.]. Formal analysis: [[M.E.N., M.Y.S., M.S.Z.]. Funding acquisition: [M.E.N., M.Y.S., M.S.Z.]. Investigation: [M.E.N., M.Y.S., M.S.Z., Q.Z., W.R.B.]. Methodology: [M.E.N., M.Y.S., M.S.Z.]. Resources: [M.S.Z., W.R.B.]. Software: [M.E.N., M.S.Z.]. Supervision: [M.Y.S., M.S.Z.]. Validation: [M.E.N.]. Visualization: [M.E.N., M.S.Z., M.Y.S.]. Writing, original draft: [M.E.N., M.S.Z.]. Writing, review and editing: [M.E.N., M.Y.S., M.S.Z., Q.Z., W.R.B.]. All authors contributed to the article and approved the submitted version.

## FUNDING

This work was partially funded by the National Institutes of Health Heart, Lung, and Blood Institute training grant T32 HL171029 (M.E.N.) and the National Institute of Immunology, Allergy, and Infectious Diseases grant R21 AI185253 (M.Y.S. and M.S.Z) and NIAID Centers of Excellence for Influenza Research and Response (CEIRR) contract 75N93021C00014/Option18C (M.Y.S.).

## ACKNOWLEDGMENTS

The authors would like to acknowledge the assistance of the University of Rochester Clinical and Translational Science Institute’s Informatics Branch with data management and storage. We thank The University of Rochester Medical Center’s Flow Cytometry Shared Resource, which receives financial support from the New York First Grant, especially Matthew Cochran, M.S., and Steven Polter, for their support and maintenance of the Cytek 5-Laser Aurora Spectral Cytometer.

## DATA AVAILABILITY STATEMENT

The cytometry datasets for this study will be available on request.

## Supplementary Material

### 1 SUPPLEMENTARY DATA

**Table S1.** Description of flow cytometry panel including markers, fluorophore, and supplier information. The optimized dilution was determined using 1e6 human splenic B cells in 50 uL of staining buffer, using a Cytek Auora 5 laser flow cytometer.

| Marker | Flour | Clone | Supplier | Cat # | Optimized Dilution |
|---|---|---|---|---|---|
| Live/Dead | Blue | – | Invitrogen | L23105 | 1:200 |
| CD19 | BUV395 | SJ25C1 | BD Horizon | 563549 | 1:200 |
| IgM | Spark Blue 550 | MHM-88 | BioLegend | 314555 | 1:200 |
| IgG | APC-H7 | G18-145 | BD Pharmingen | 561297 | 1:200 |
| CD73 | BUV615 | AD2 | BD Opti-Build | 751369 | 1:200 |
| CD20 | eFlour450 | 2H7 | Invitrogen | 48-0209-42 | 1:200 |
| IgA | VioGreen | IS11-8E10 | Miltenyi Biotec | 130-113-481 | 1:200 |
| CD3 | BV570 | UCHT1 | BioLegend | B171767 | 1:200 |
| CD14 | BV570 | M5E2 | BioLegend | B372136 | 1:200 |
| CD16 | BV570 | 3G8 | BioLegend | B379013 | 1:200 |
| IgD | BV785 | IA6-2 | BioLegend | 348242 | 1:200 |
| CD21 | PE-Cy7 | B-Ly4 | BD Pharmingen | 561374 | 1:200 |
| CD10 | PE/Cy5 | HI10a | BD Pharmigen | 563509 | 1:200 |
| CD71 | BV650 | CY1G4 | BioLegend | 334116 | 1:100 |
| CD24 | BV421 | ML5 | BioLegend | 311122 | 1:100 |
| CD185 | BV480 | RF882 | BD Horizon | 566142 | 1:100 |
| CD95 | BV605 | DX2 | BioLegend | 305628 | 1:100 |
| CD62L | NovaFlour Blue 585 | DREG56 | Invitrogen | H009T03B04 | 1:100 |
| CD184 (CXCR4) | RY586 | 12G5 | Invitrogen | 569646 | 1:100 |
| CD85j | PE/Dazzle 594 | ILT2 | BioLegend | 33716 | 1:100 |
| CD183 | PE/Fire810 | G025H7 | BioLegend | 353760 | 1:100 |
| CD1c | Alexa Flour 647 | L161 | BioLegend | 331510 | 1:100 |
| CD197 | Spark NIR 685 | G043H7 | BioLegend | 353258 | 1:100 |
| CD38 | APC/Fire810 | HIT2 | BioLegend | 303550 | 1:100 |
| CD27 | BUV805 | M-T271 | BD Opti-Build | 742012 | 1:67 |
| CD11c | PE/Fire700 | Bu15 | BioLegend | 337248 | 1:67 |
| CD138 | PerCP-eFlour710 | MI15 | Invitrogen | 46-1388-42 | 1:67 |
| CD307e | BUV563 | 509F6 | BD Opti-Build | 749598 | 1:50 |
| CD69 | BUV661 | FN50 | Invitrogen | 376-0699-42 | 1:50 |
| CD45RB | APC | MEM-55 | Invitrogen | MA1-19461 | 1:40 |
| CD23 | PE | M-L233 | BD Pharmingen | 555711 | 1:10 |

**Figure S1.**
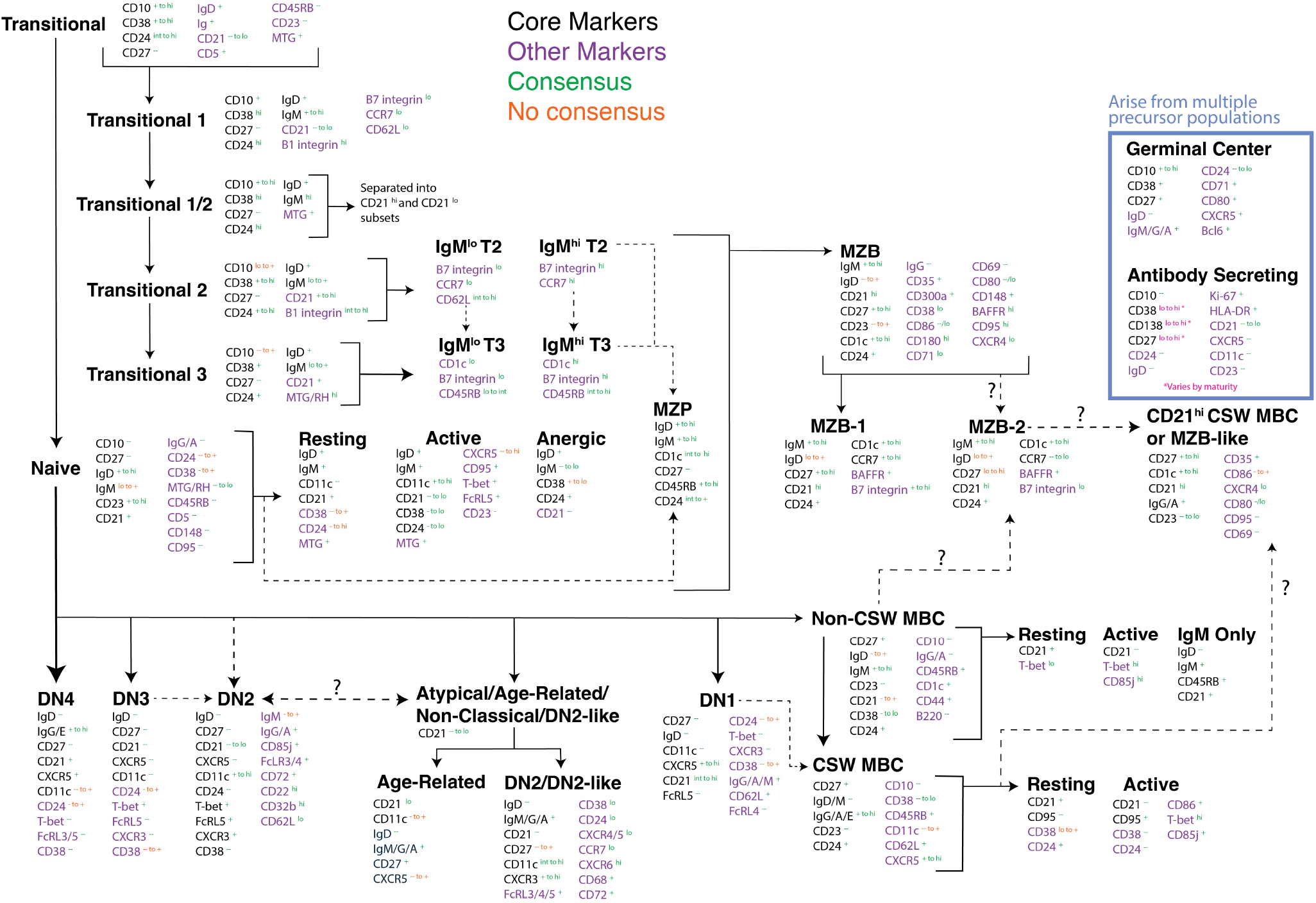
Diagram of B cell classification across 27 published studies (see Supplemental Table 1). Subset names are in bold black text. Core markers (black text) are used to identify a subset by 1 or more studies, and other markers are associated with a phenotype but are not used in isolation from the subset. Superscript in green indicates the marker expression is agreed upon by all who use that given marker. Superscript in orange indicates that at least 2 manuscripts differ in the usage of the marker. All solid black arrows indicate a known differentiation pathway, while dashed arrows indicate a potential or hypothesized pathway of differentiation.

**Figure S2.**
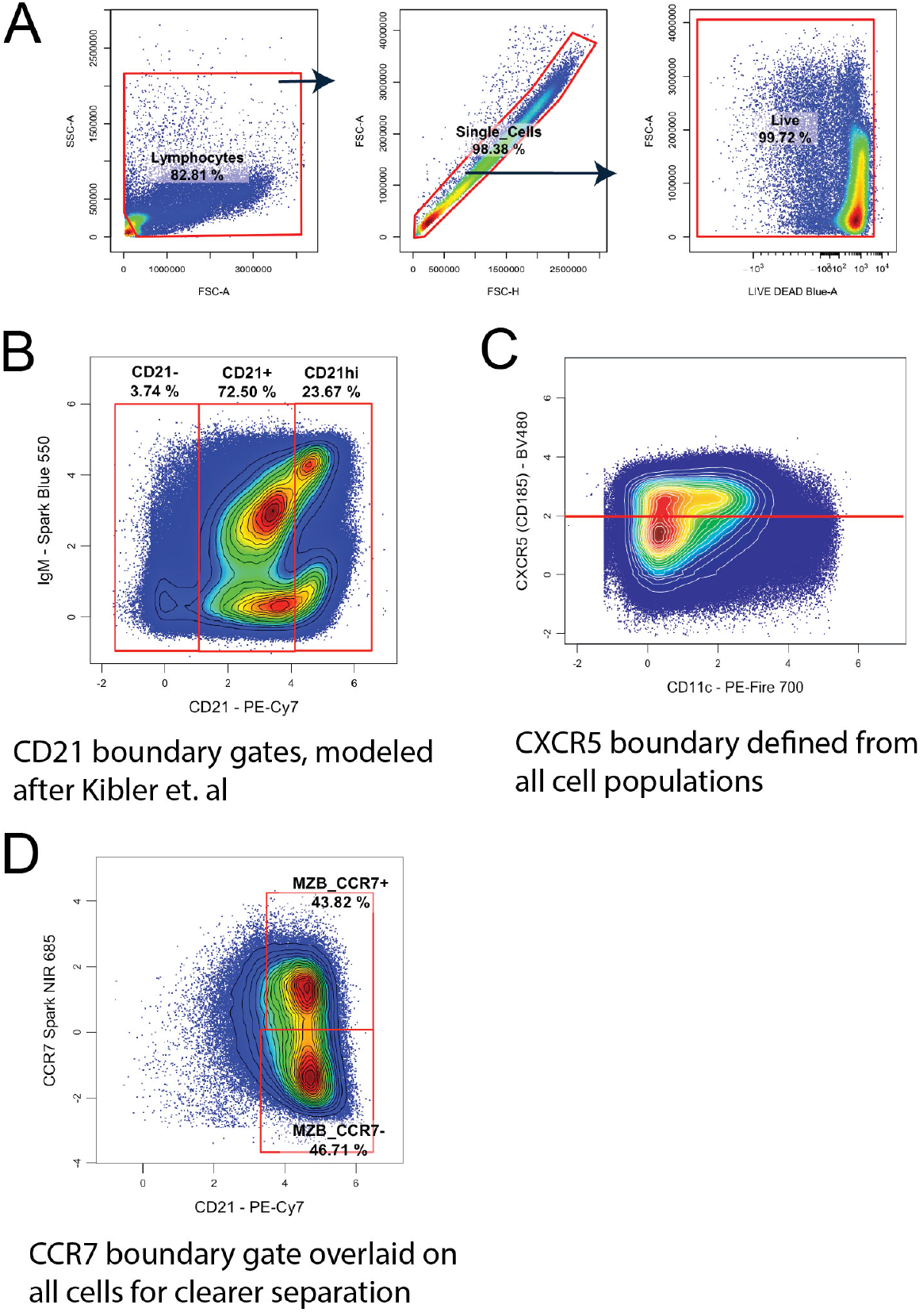
(**A**) Initial gating was applied to all samples to isolate lymphocytes, single cells, and live cells. (**B**) Boundary used to set CD21^hi^ versus CD21^+^ gate as described by Kibler et al. using all CD19^+^ CD20^+^ B cells. (3) (**C**) Boundary used to separate CXCR5 (CD185) positive and negative cells, applied to all live, single cells. (**D**) CCR7 positive and negative gates applied to all live, single cells.

**Figure 3a.**
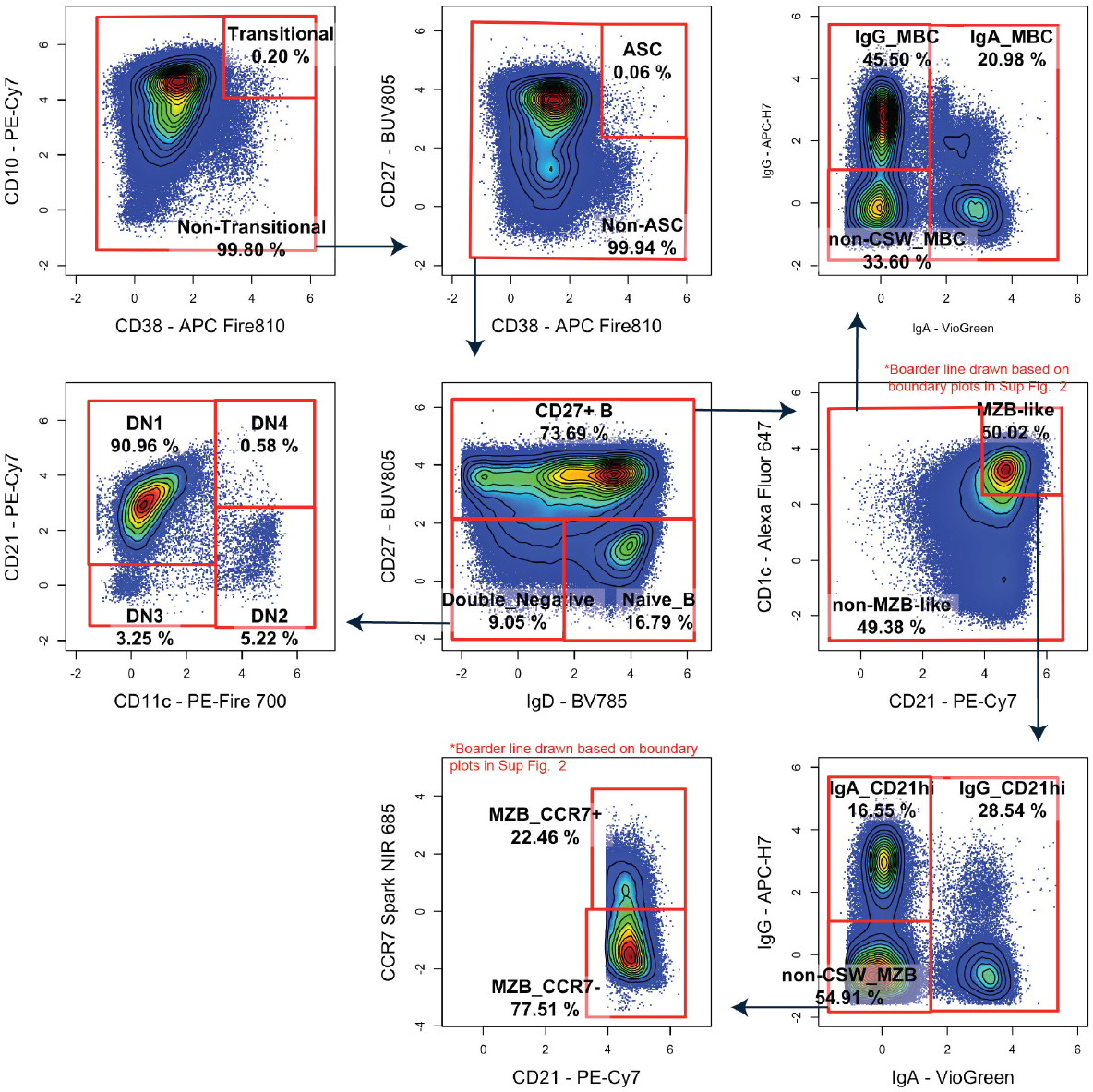
Niska gating scheme applied to pre-gated live, single B cells (CD3^-^ CD14^-^ CD16^-^ CD19^+^ CD20^+^).

**Figure 3b.**
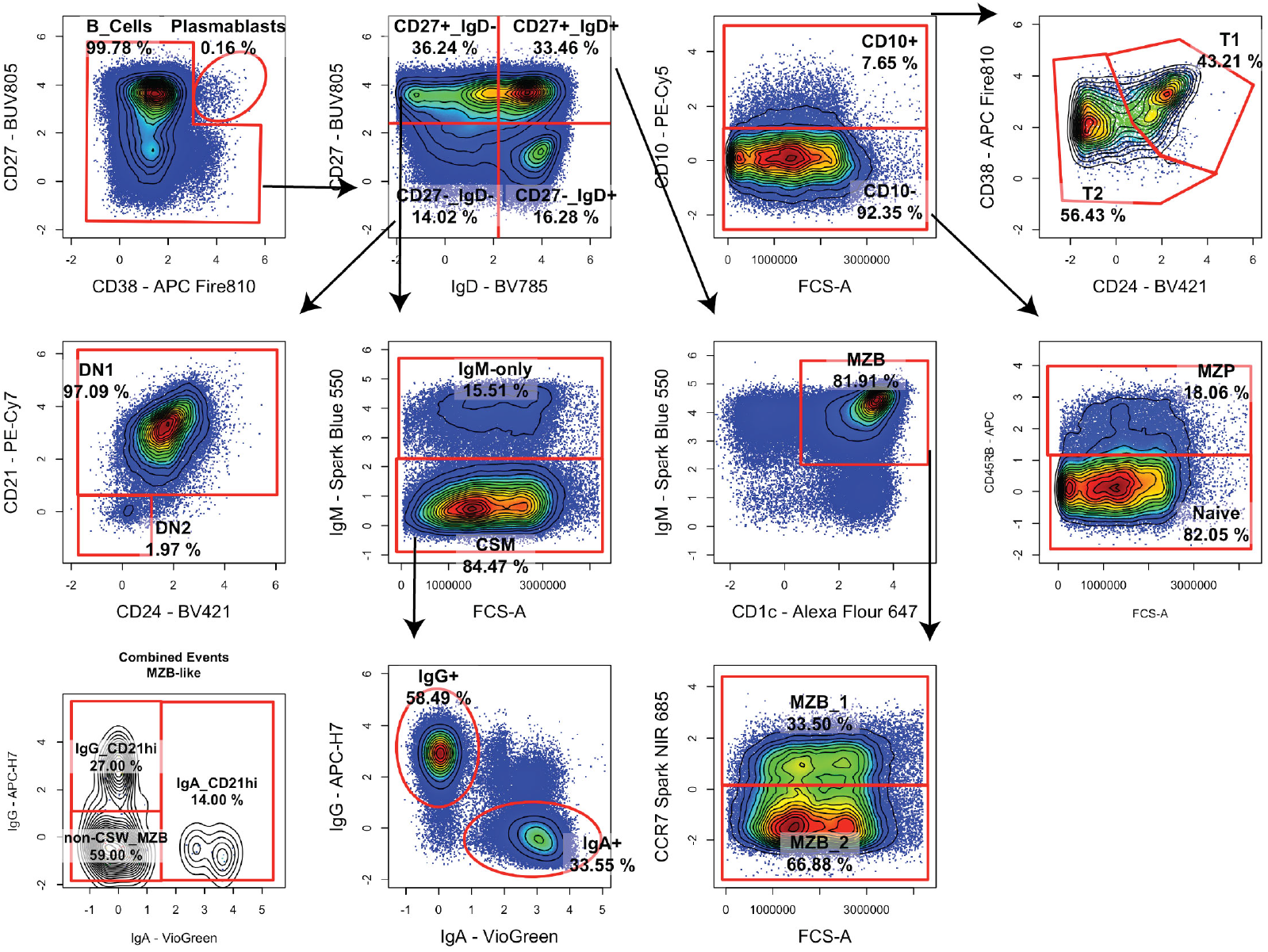
Velounias et al. gating scheme applied to pre-gated live, single B cells (CD3^-^ CD14^-^ CD16^-^ CD19^+^ CD20^+^). (30)

**Figure 3c.**
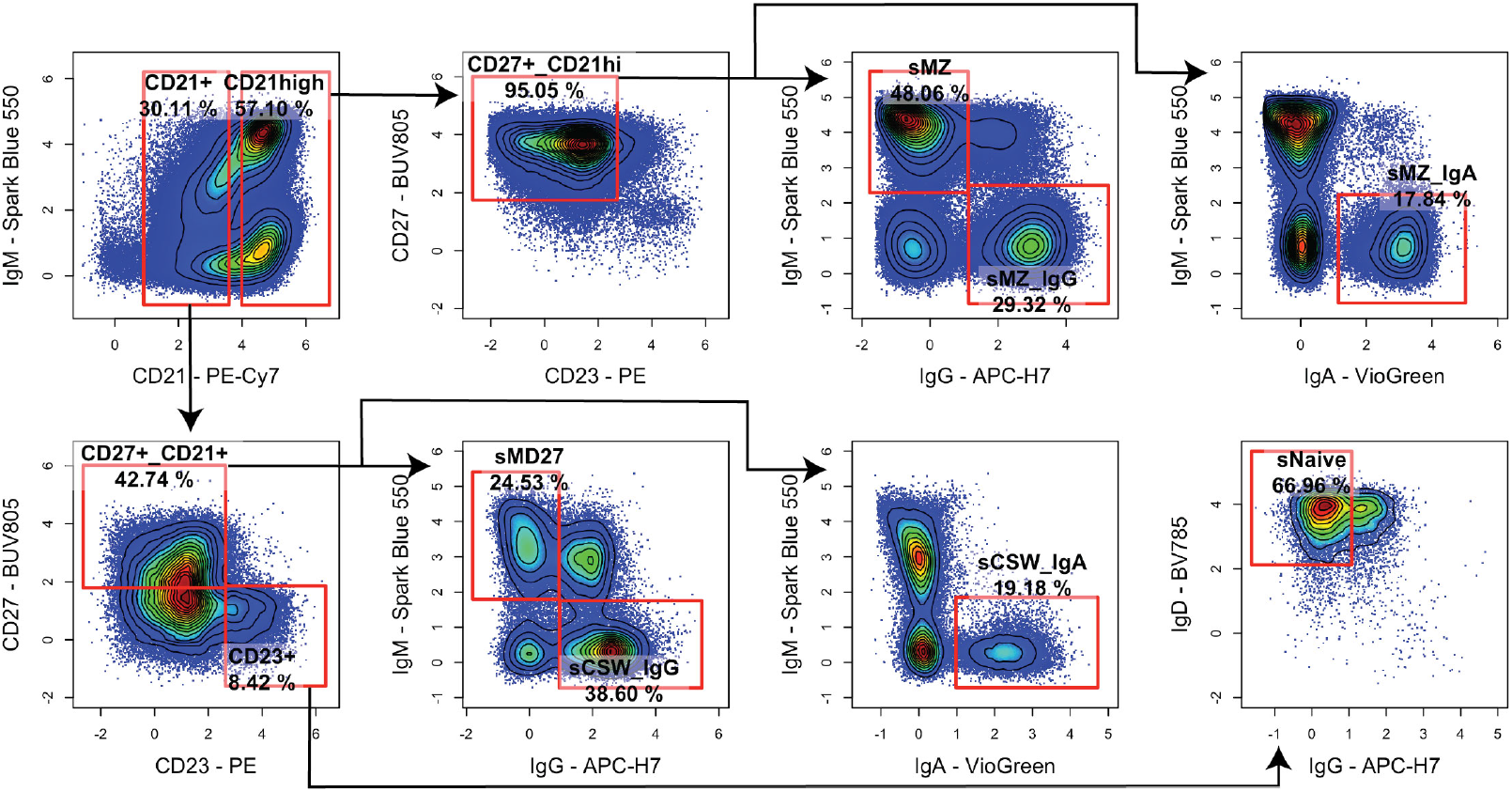
Kibler et al. gating scheme applied to pre-gated live, single B cells (CD3^-^ CD14^-^ CD16^-^ CD19^+^ CD20^+^.(3)

**Figure 3d.**
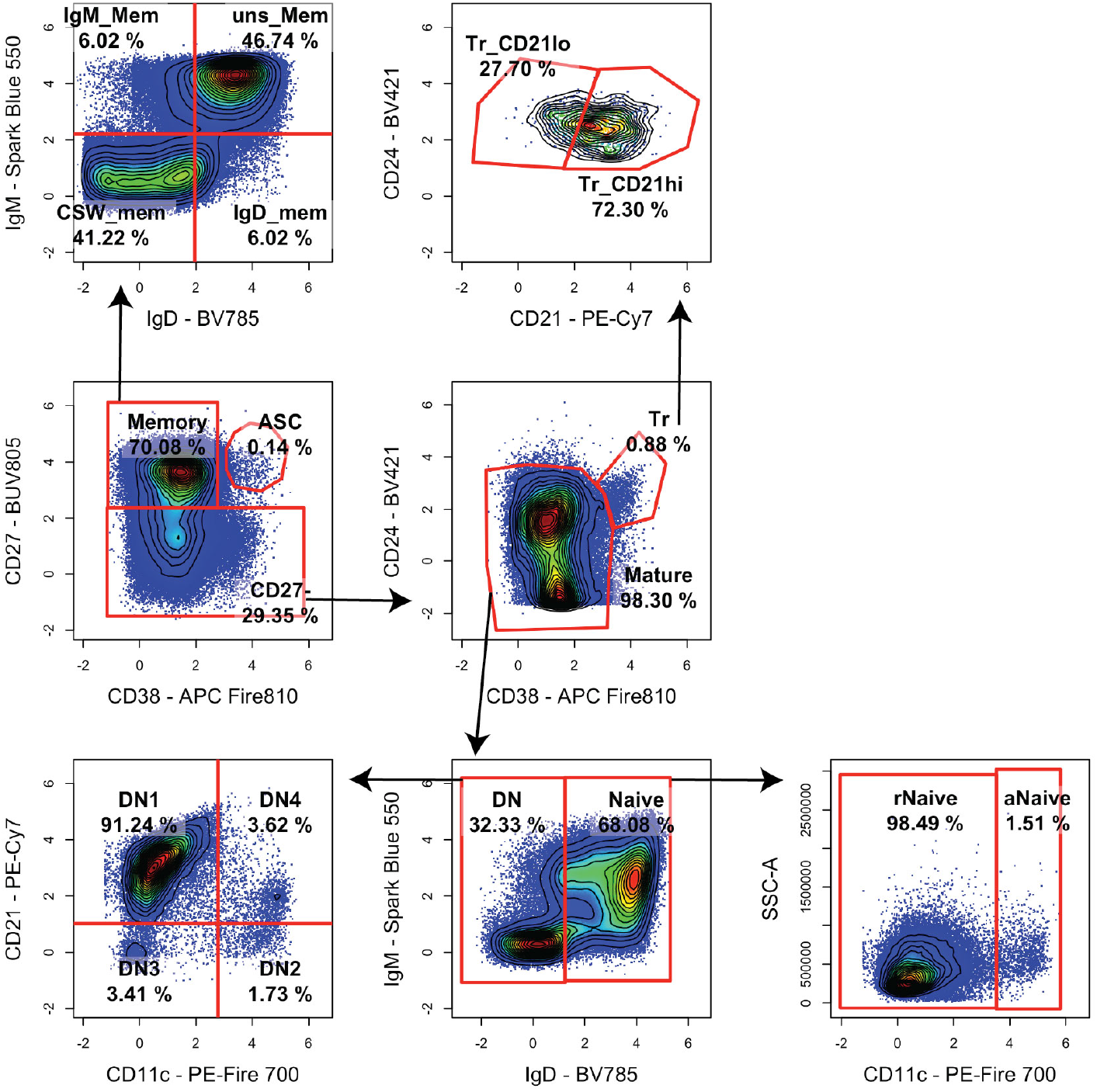
Woodruff et al. gating scheme applied to pre-gated live, single B cells (CD3^-^ CD14^-^ CD16^-^ CD19^+^ CD20^+^). (15)

**Figure 3e.**
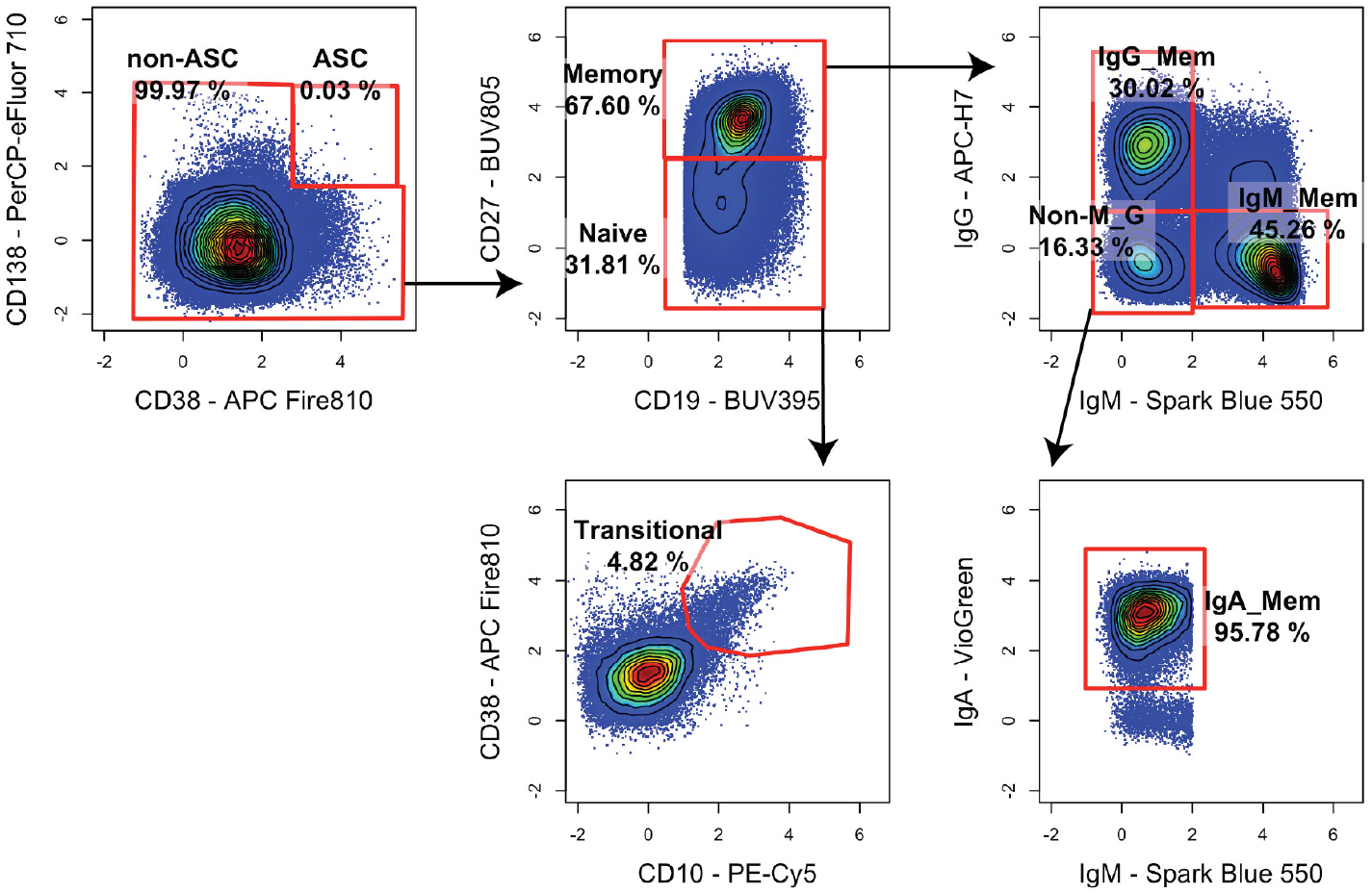
Weisel et al. gating scheme applied to pre-gated live, single B cells (CD3^-^ CD14^-^ CD16^-^ CD19^+^ CD20^+^). (31)

**Figure 3f.**
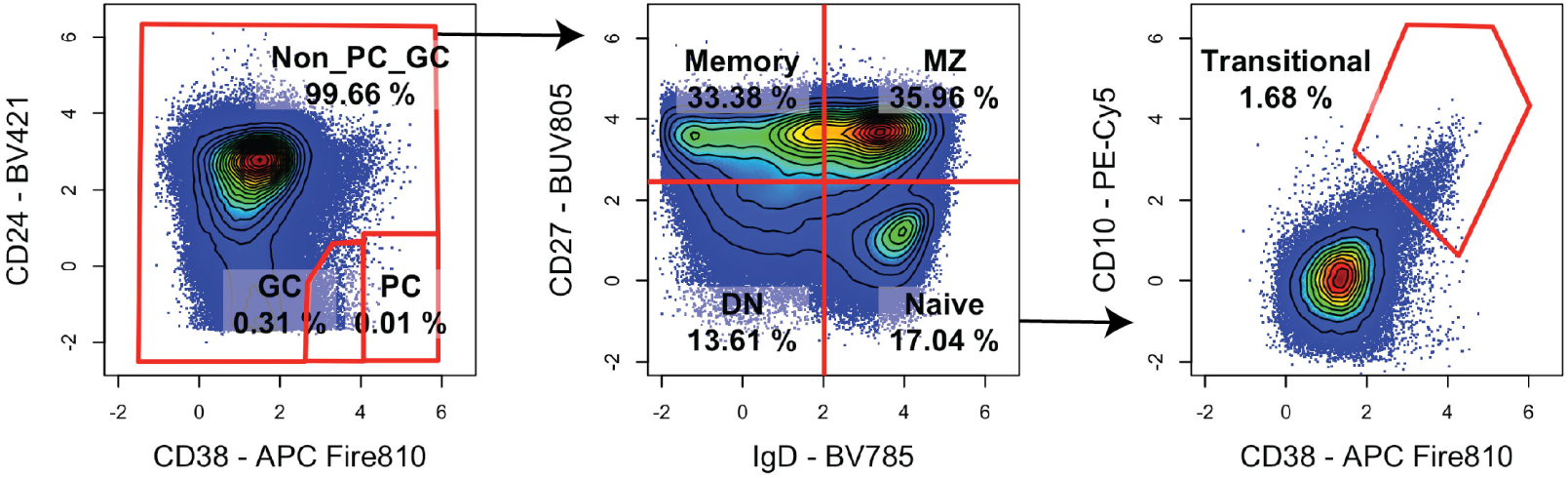
Crickx et al. gating scheme applied to pre-gated live, single B cells (CD3^-^ CD14^-^ CD16^-^ CD19^+^ CD20^+^). (34)

**Figure 3g.**
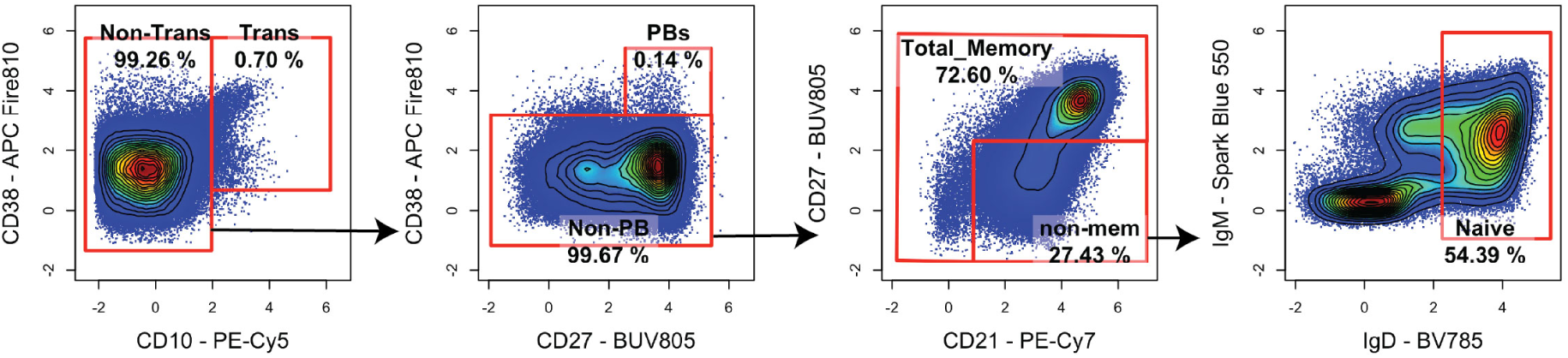
Knox et al. gating scheme applied to pre-gated live, single B cells (CD3^-^ CD14^-^ CD16^-^ CD19^+^ CD20^+^). (12)

**Figure 3h.**
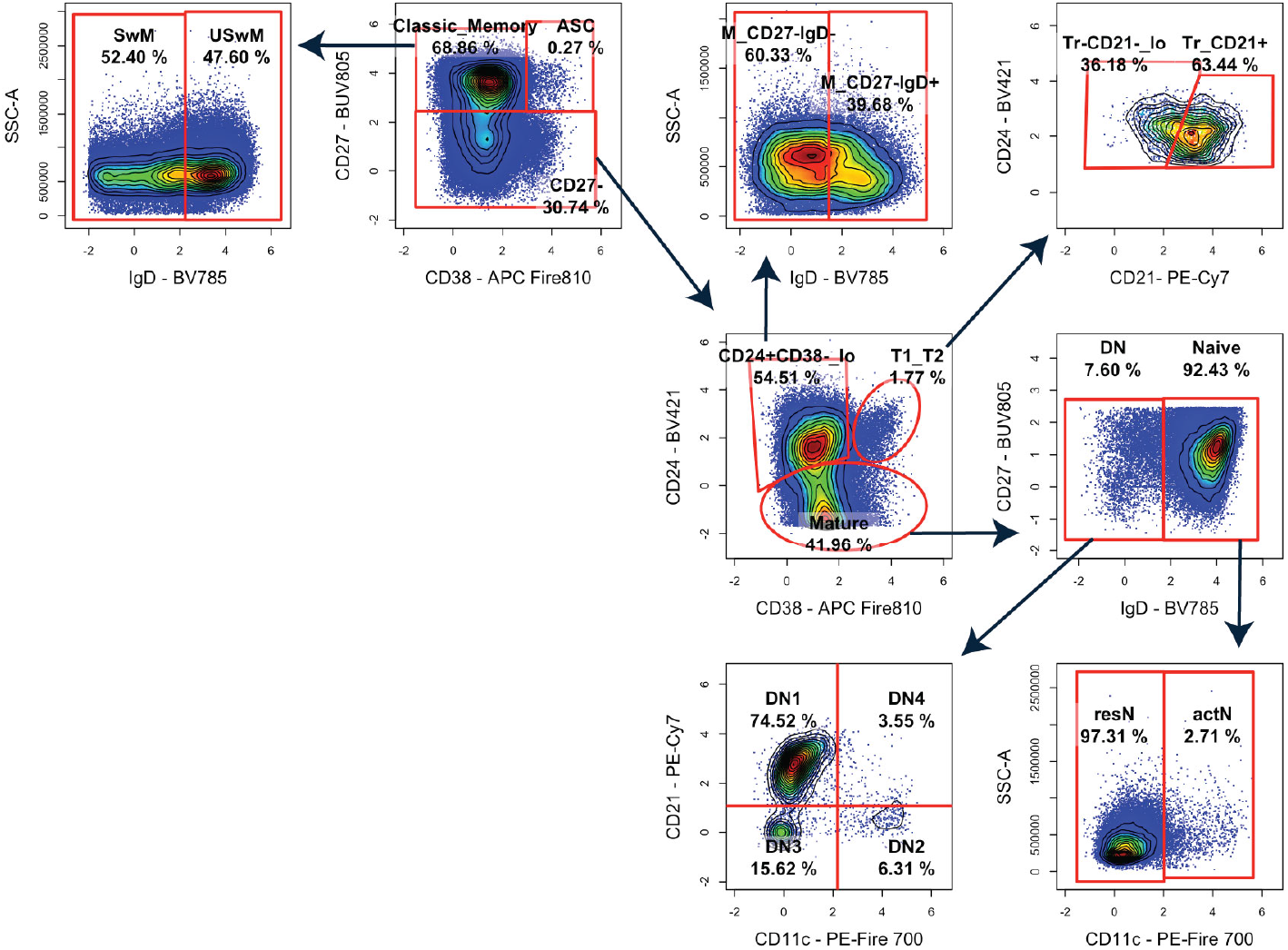
Sosa-Hernandez et al. gating scheme applied to pre-gated live, single B cells (CD3^-^ CD14^-^ CD16^-^ CD19^+^ CD20^+^). (28)

**Figure S4.**
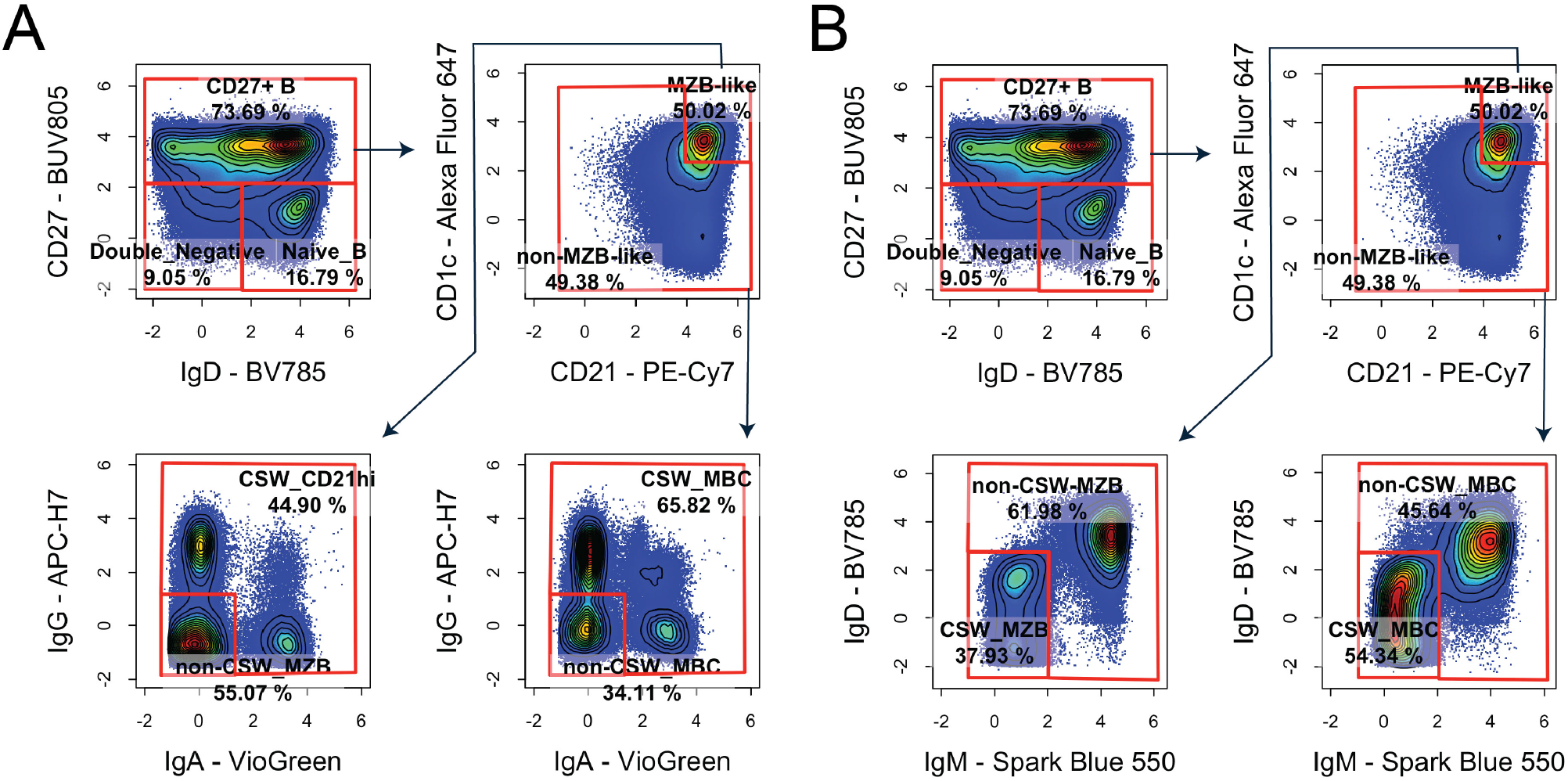
Gating with IgG/A for isotype applied to pre-gated live, single B cells (CD3^-^ CD14^-^ CD16^-^ CD19^+^ CD20^+^) by (**A**) defining CSW based on IgG/A expression or (**B**) defining CSW based on lack of IgD/M expression.

**Figure 5a.**
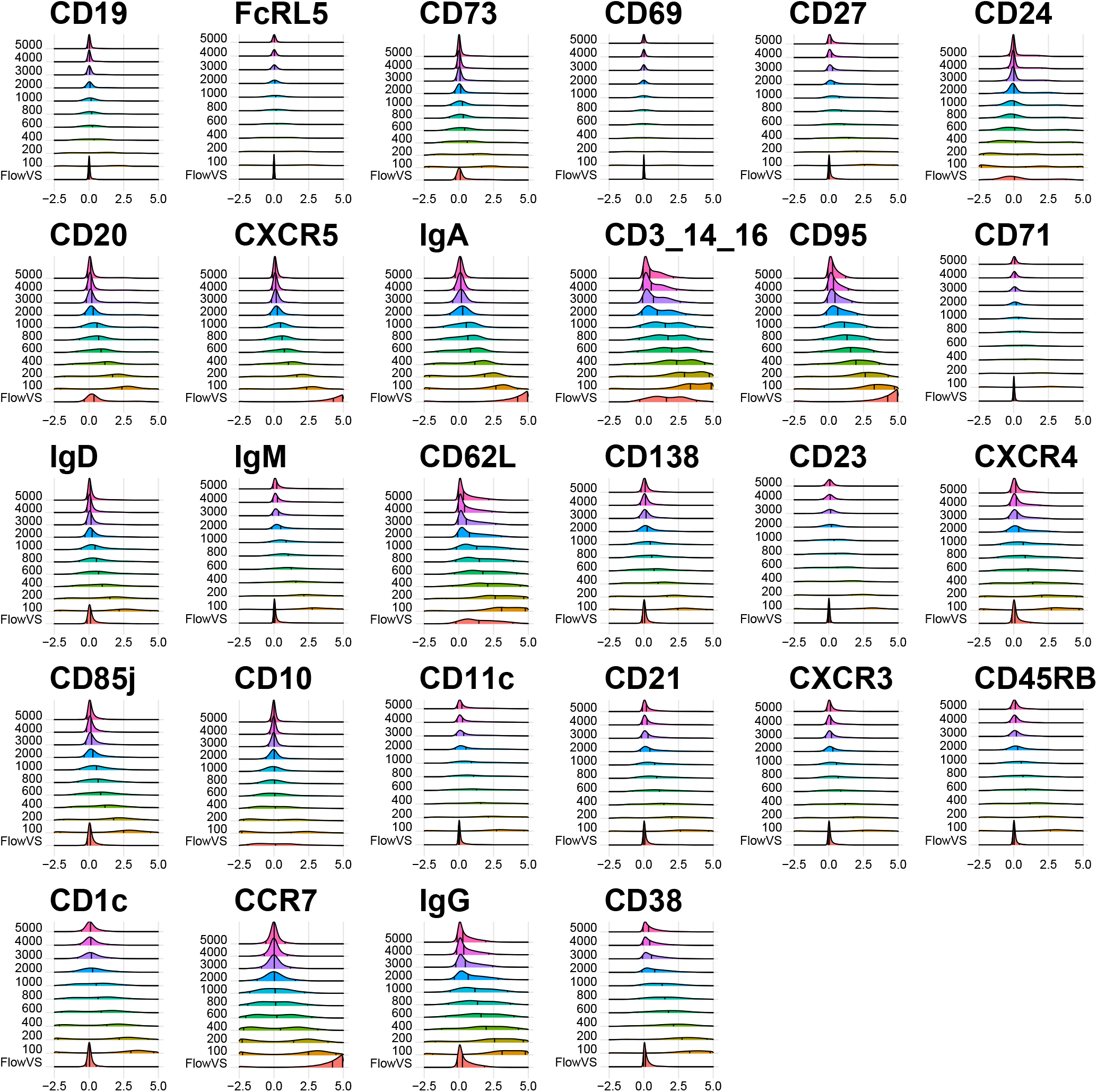
FlowVS optimized and selected cofactors on all live, single-cell splenocytes

**Figure 5b.**
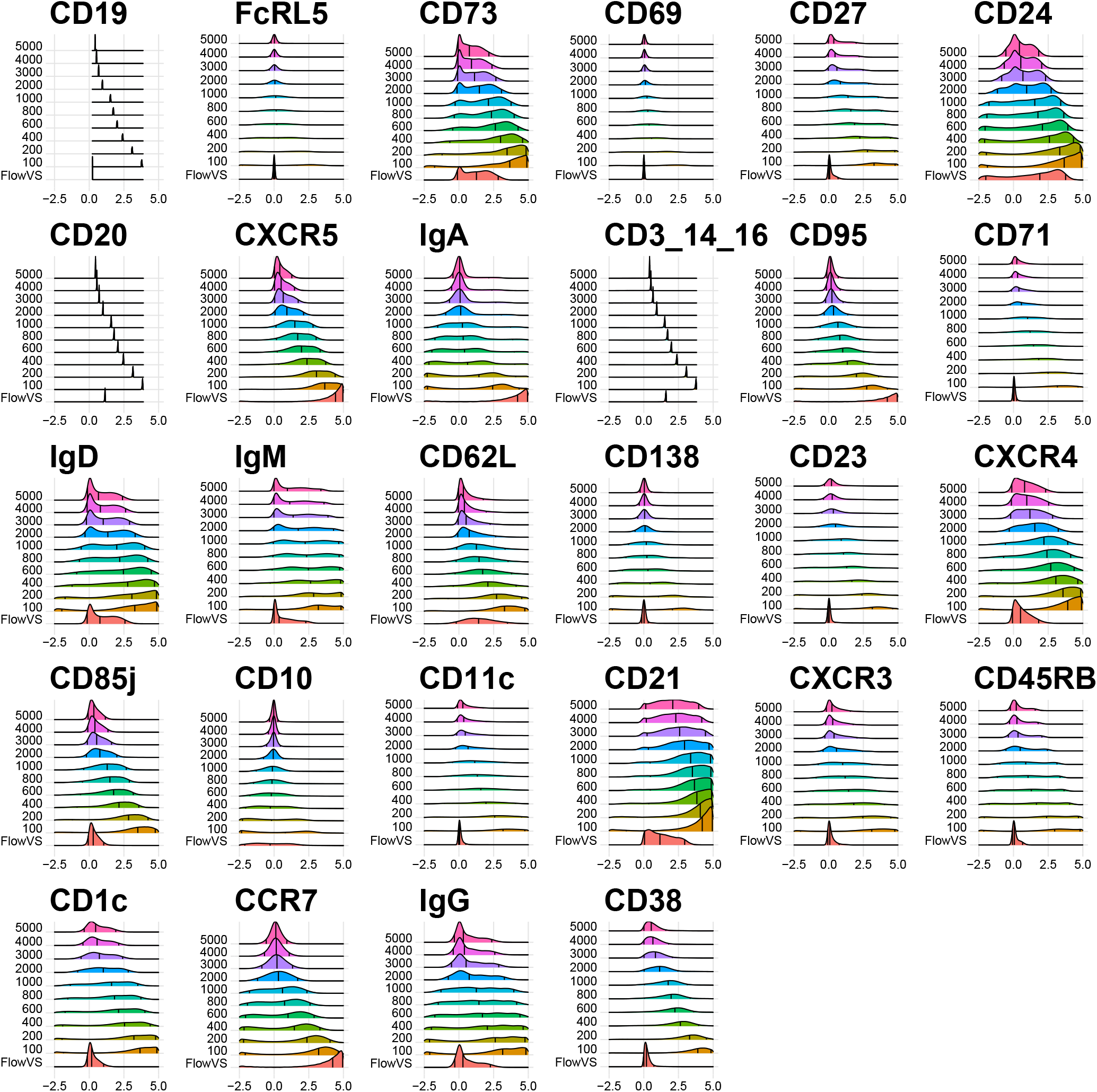
FlowVS optimized and selected cofactors on CD19^+^ CD20^+^ B cells

**Figure S6.**
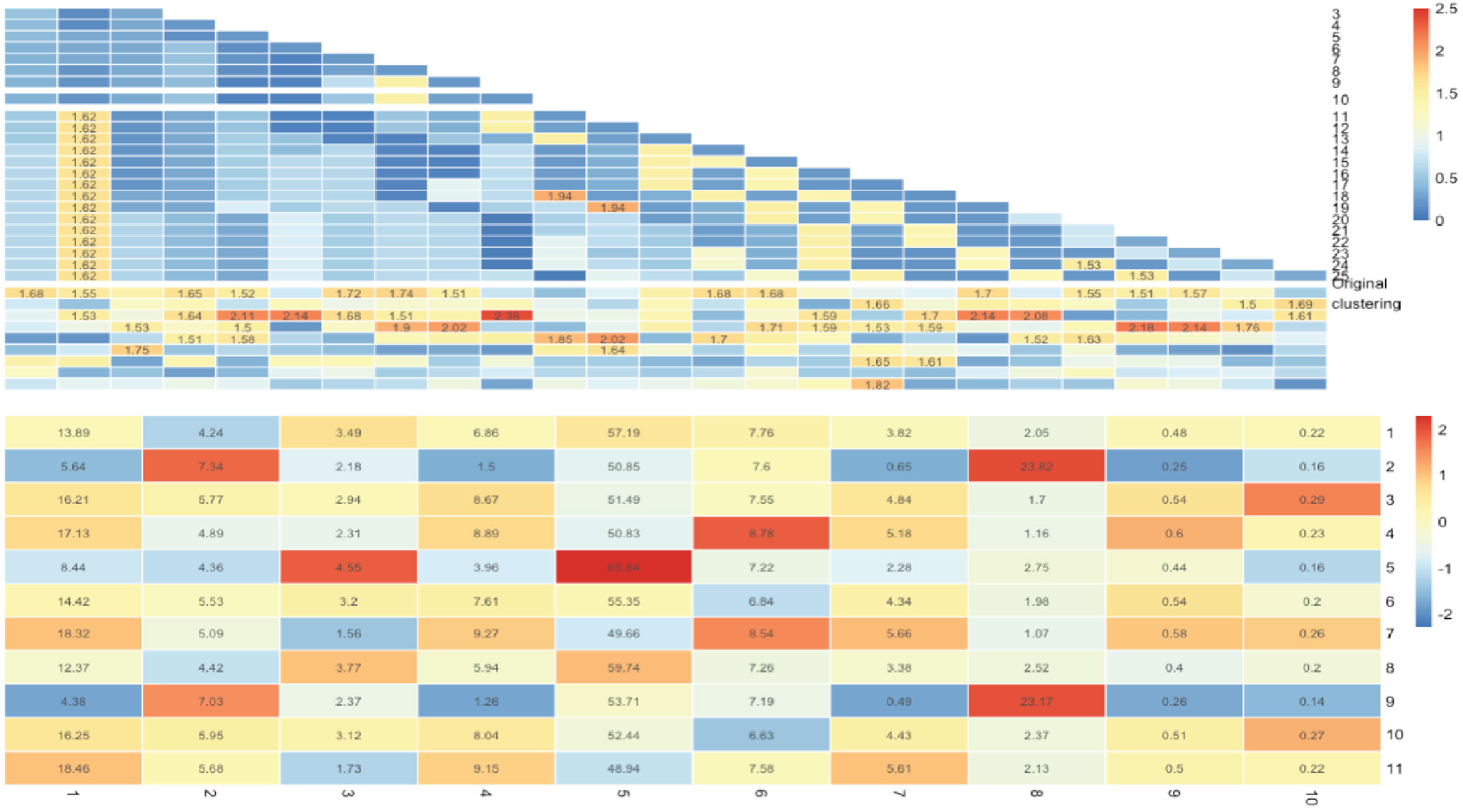
`CytoNorm` output for determining the number of clusters that exist within the dataset using the CV to show differences between batches. Cluster numbers (right sid(**E**) for which all CVs are below 1.8 are less likely to be clustered based on the batch effect.

**Figure 7a.**
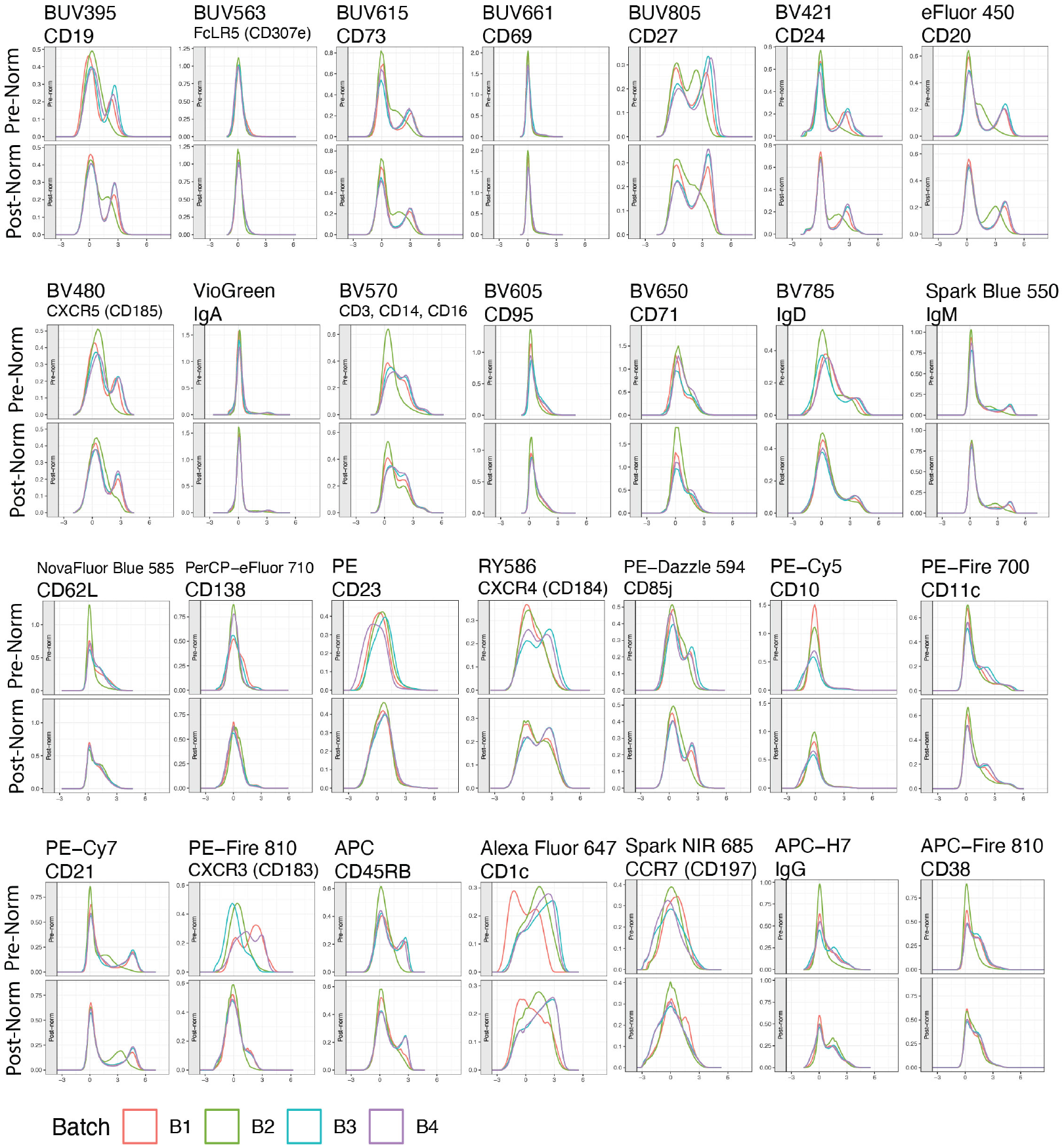
Histogram of fluorescence by channel pre- and post-normalization, where each batch is represented by color according to the key on the bottom.

**Figure 7b.**
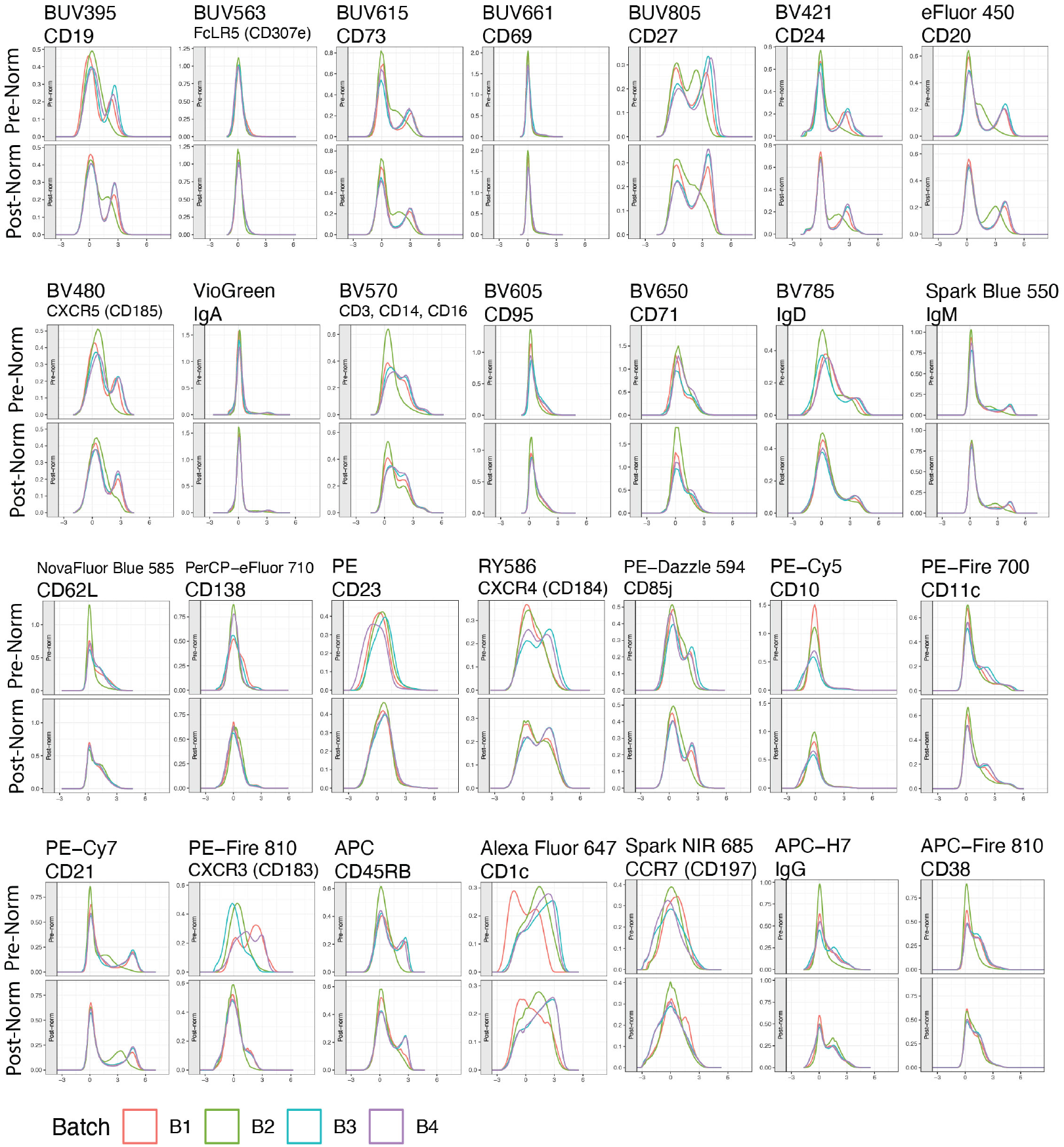
2D plots of all live, single cells with optimized cofactor for the markers displayed on the x- and y-axis.

**Figure 8a.**
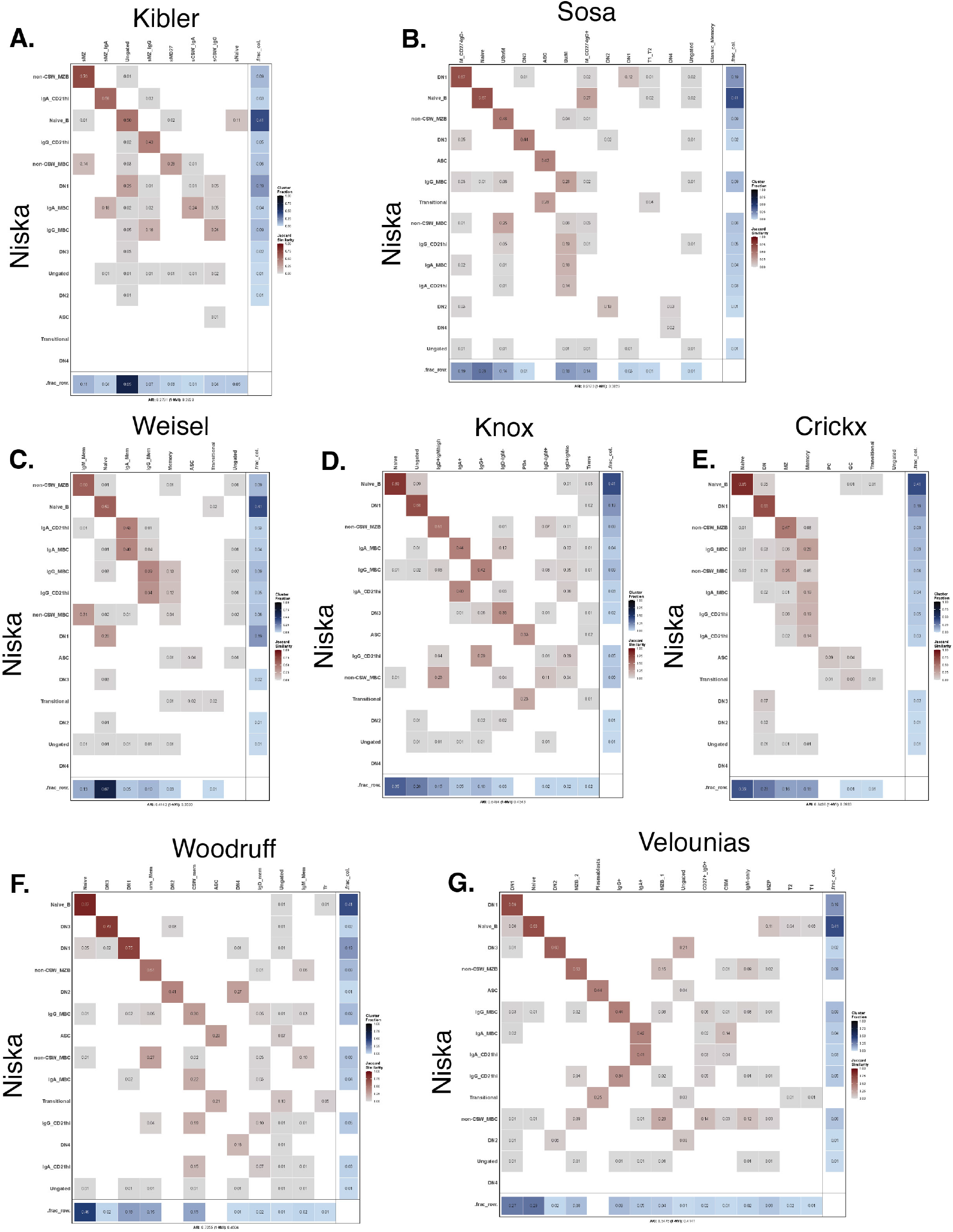
JSI for the Niska gating scheme versus (**A**) Kibler, (**B**) Sosa-Hernandez, (**C**) Weisel, (**D**) Knox, (**E**) Crickx, (**F**) Woodruff, and (**F**) Velounias gating schemes as shown in **Supplemental Figure 3**.

**Figure 8b.**
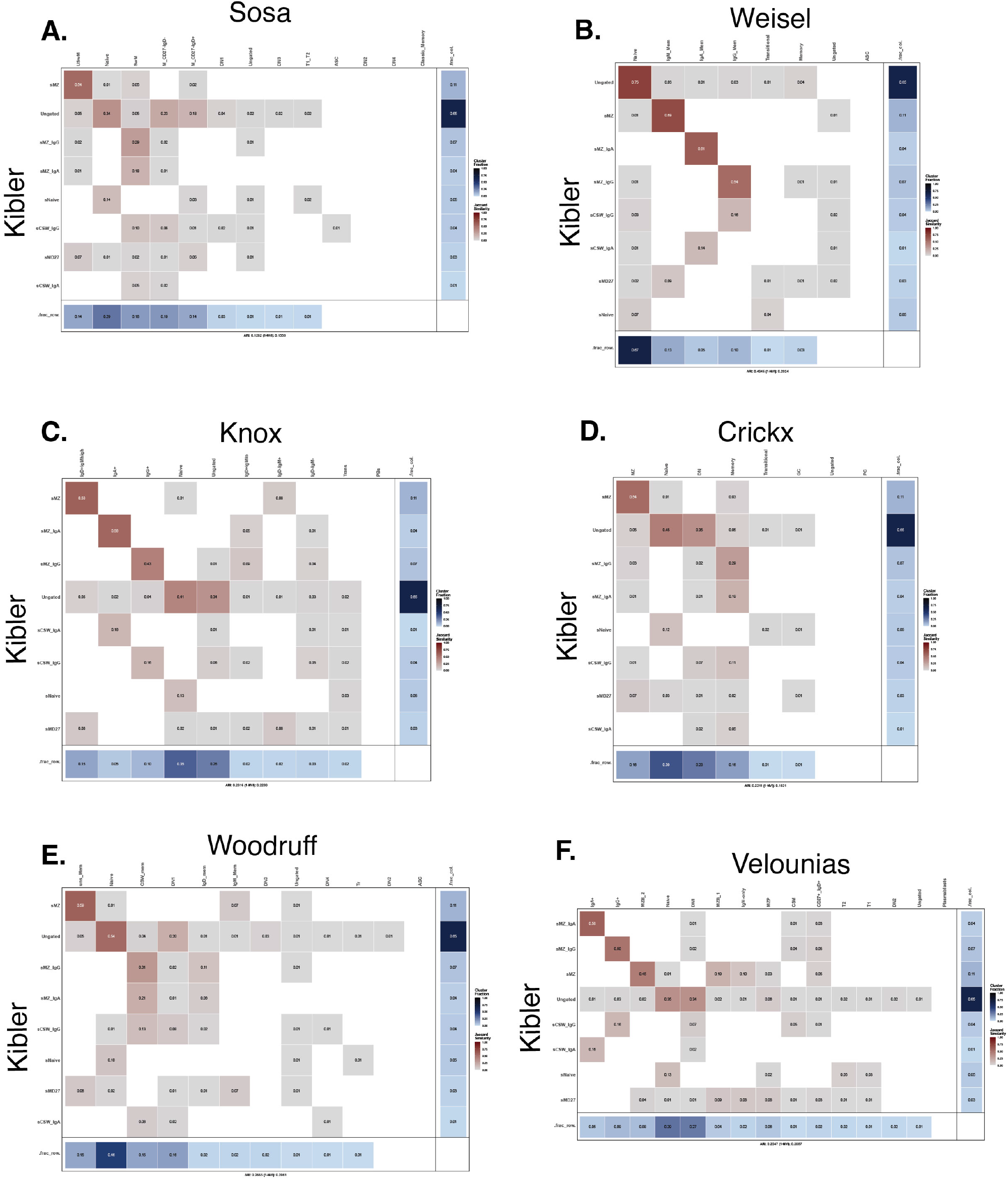
JSI for the Kibler gating scheme versus (**A**) Sosa-Hernandez, (**B**) Weisel, (**C**) Knox, (**D**) Crickx, (**E**) Woodruff, and (**F**) Velounias gating schemes as shown in **Supplemental Figure 3**.

**Figure 8c.**
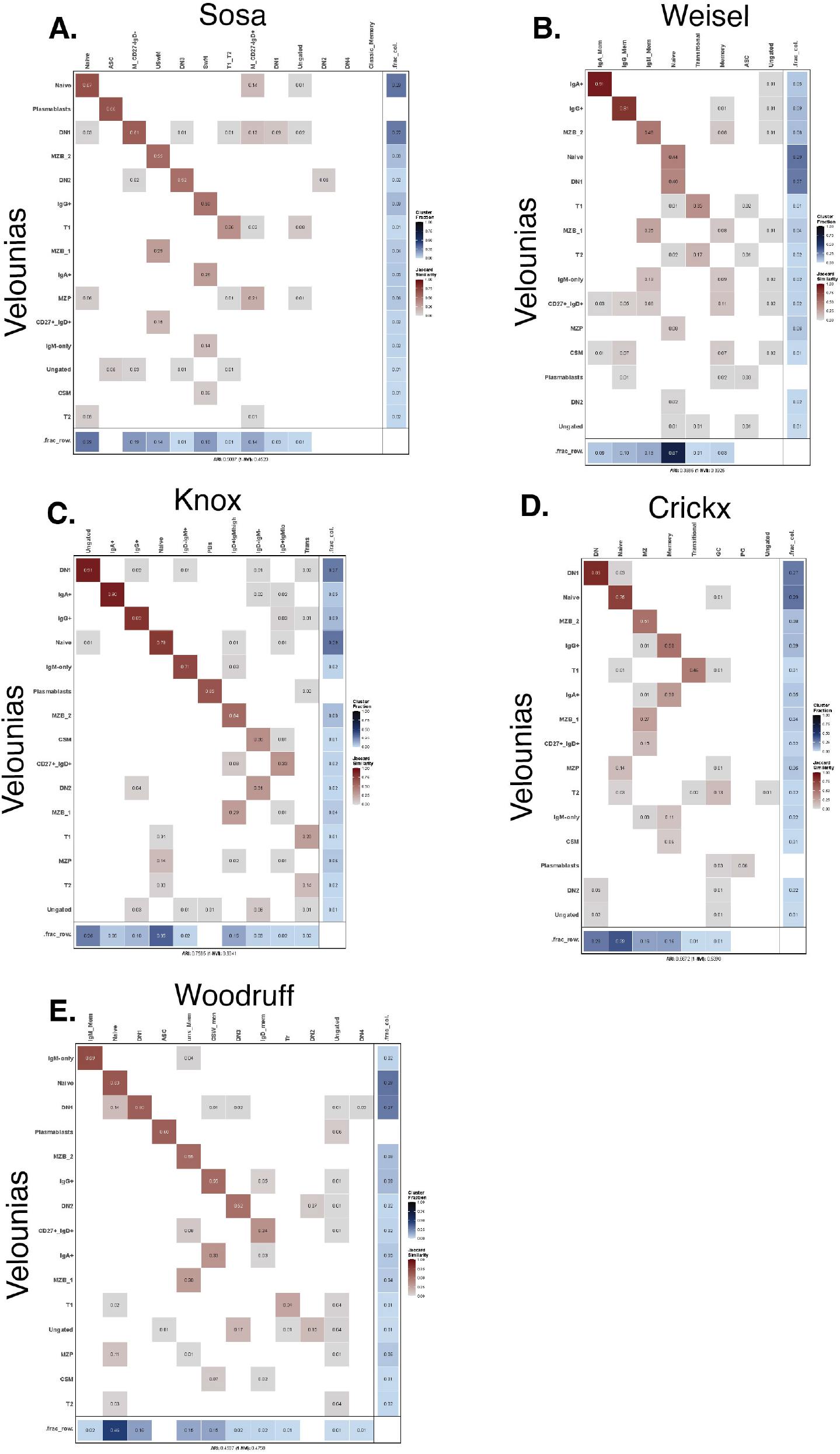
JSI for the Velounias gating scheme versus (**A**) Sosa, (**B**) Weisel, (**C**) Knox, (**D**) Crickx, and (**E**) Woodruff gating schemes as shown in **Supplemental Figure 3**.

**Figure 8d.**
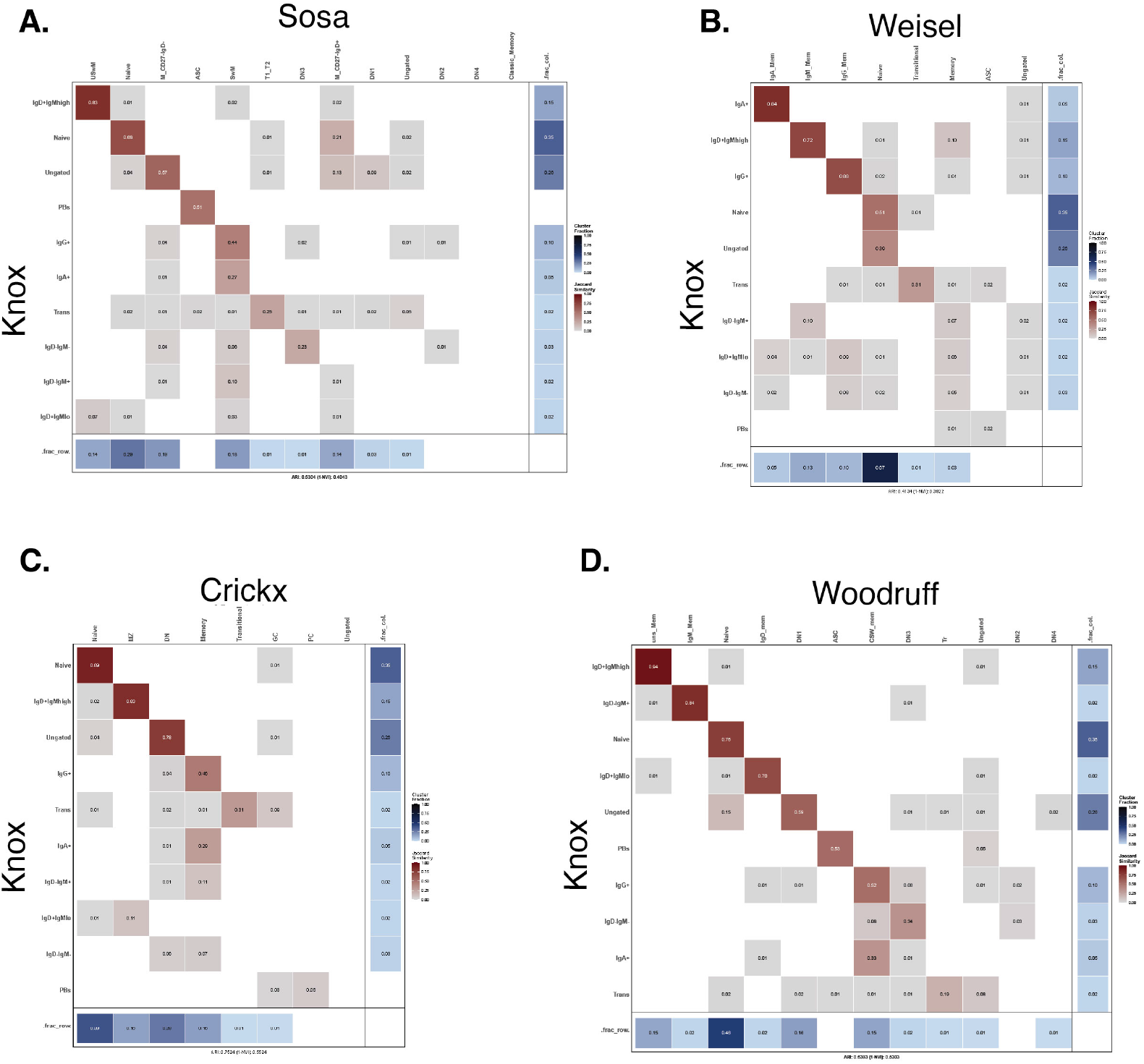
JSI for the Knox gating scheme versus (**A**) Sosa-Hernandex, (**B**) Weisel, (**C**) Crickx, and (**D**) Woodruff gating schemes as shown in **Supplemental Figure 3**.

**Figure 8e.**
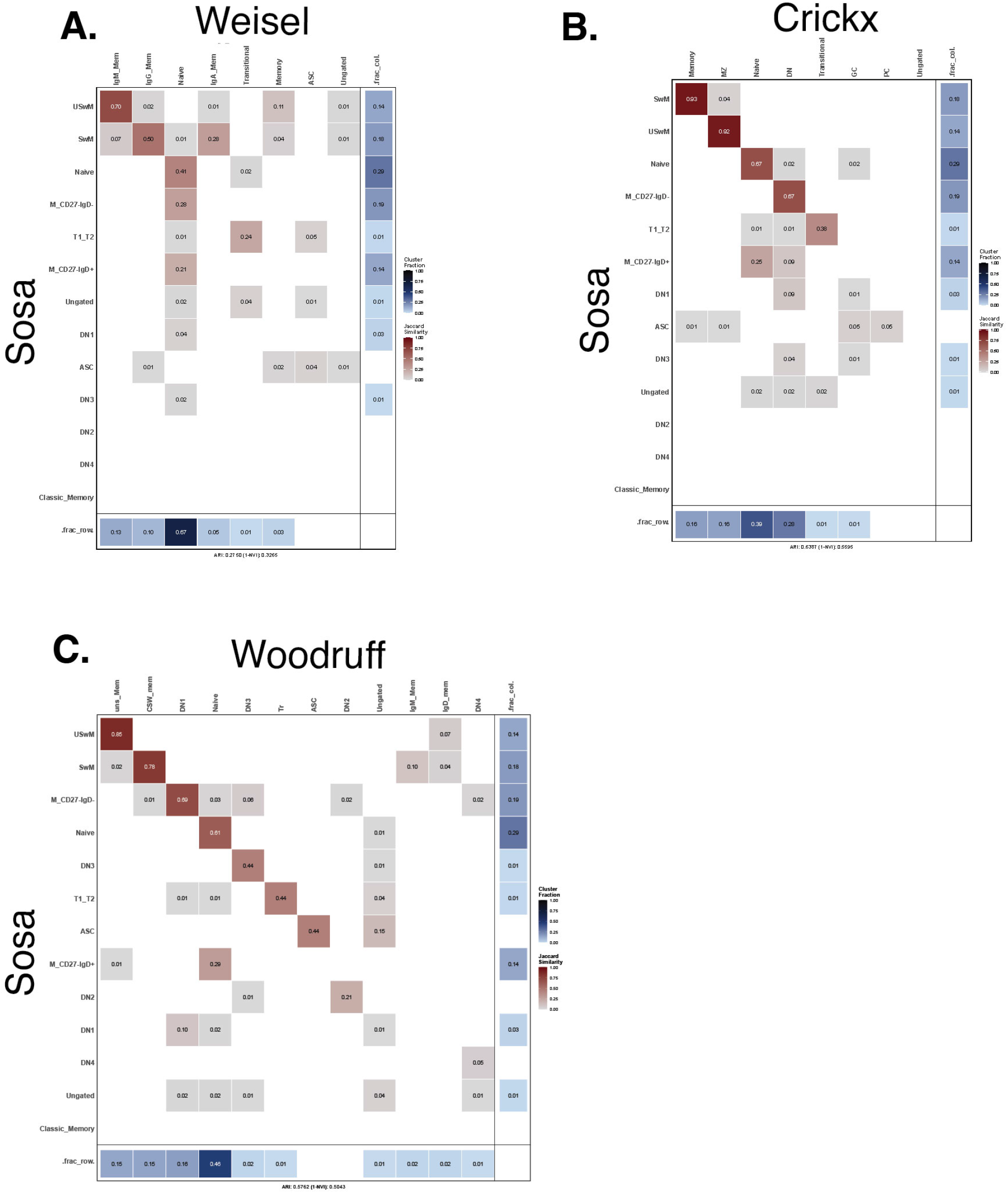
JSI for the Sosa-Hernandez gating scheme versus (**A**) Weisel, (**B**) Crickx, and (**C**) Woodruff gating schemes as shown in **Supplemental Figure 3**.

**Figure 8f.**
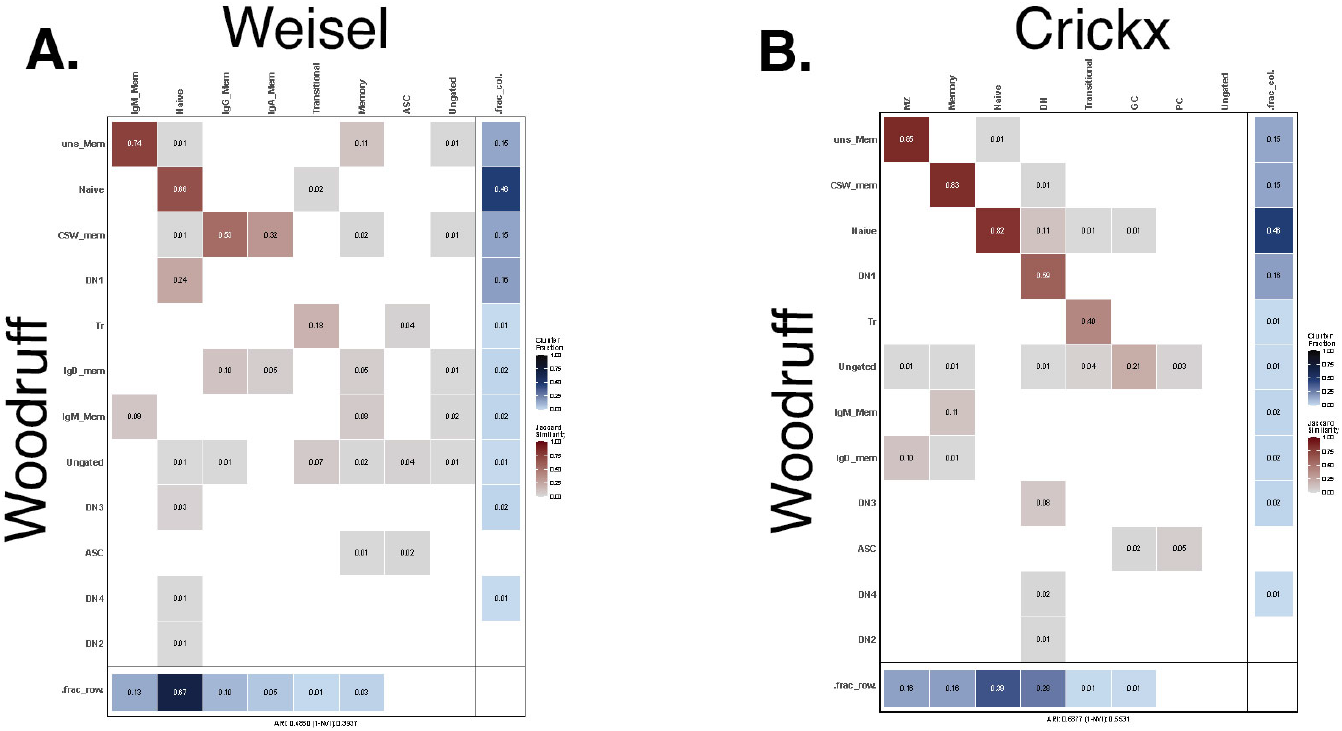
JSI for the Woodruff gating scheme versus (**A**) Weisel and (**B**) Crickx gating schemes as shown in **Supplemental Figure 3**.

**Figure 8g.**
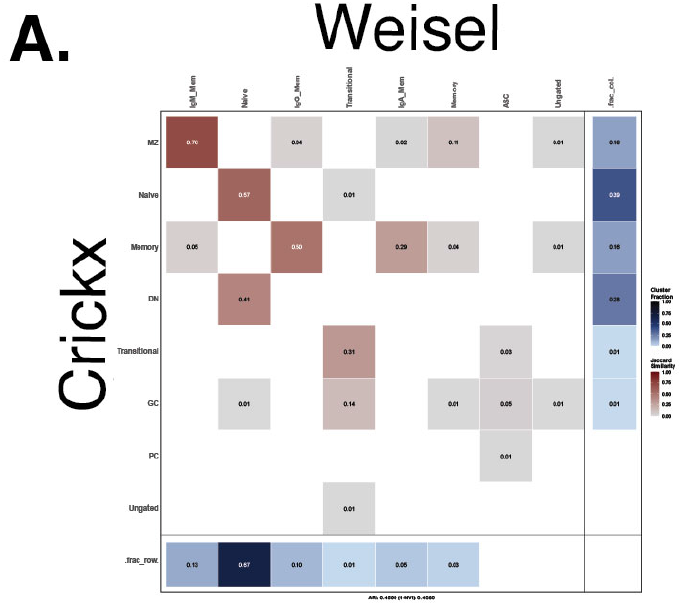
JSI for the Crickx gating scheme versus Weisel gating schemes as shown in **Supplemental Figure 3**.

**Figure 9a.**
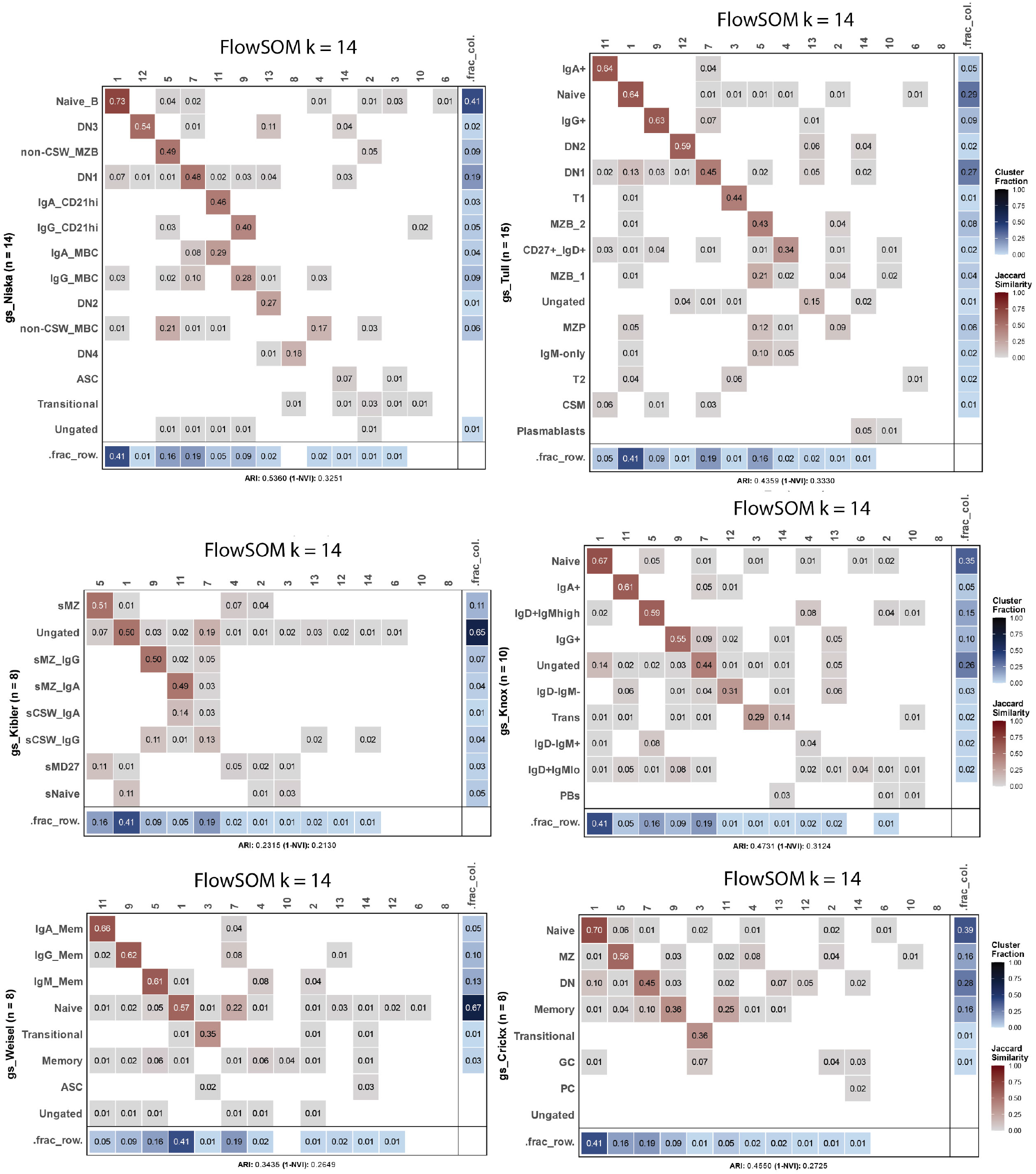
JSI for FlowSOM with 14 metaclusters (*k* = 14) versus gating schemes illustrated in **Supplemental Figure 3**.

**Figure 9b.**
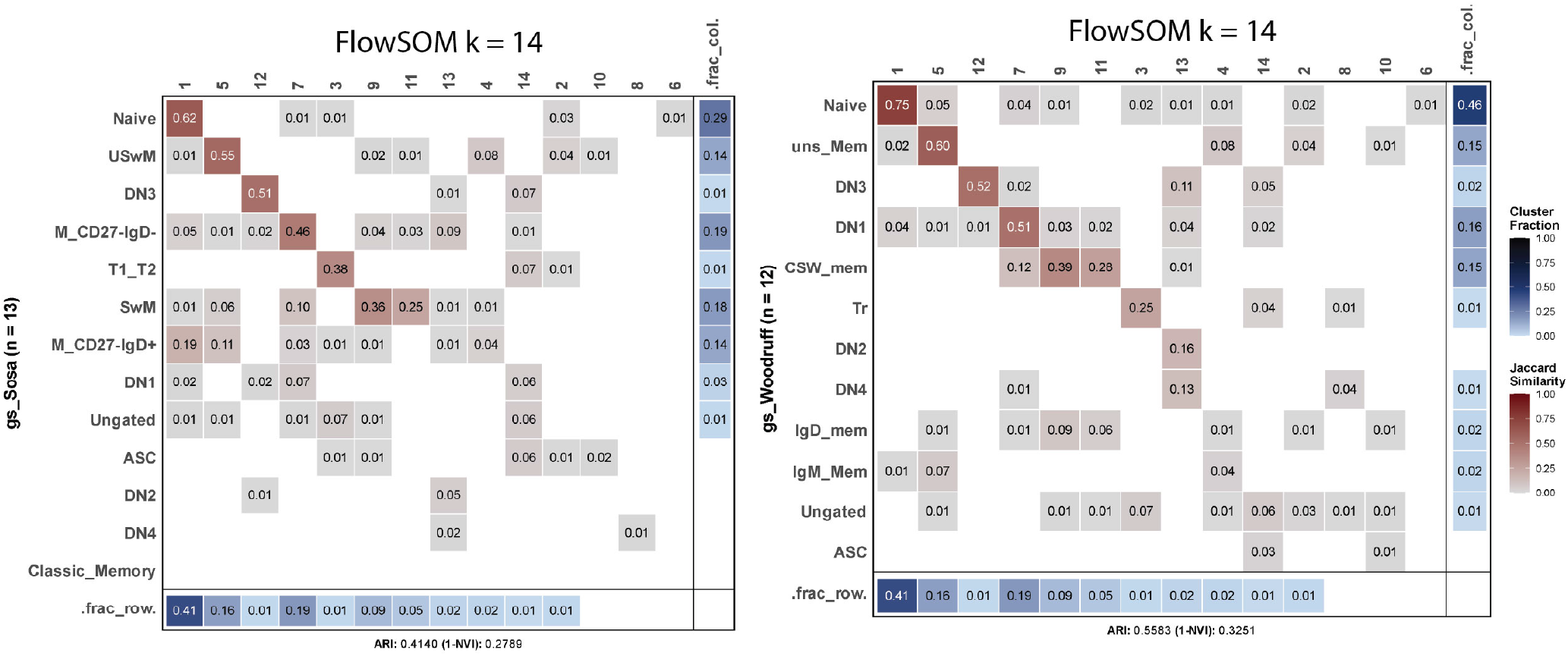
JSI for FlowSOM with 14 metaclusters (*k* = 14) versus gating schemes illustrated in **Supplemental Figure 3**.

**Figure 9c.**
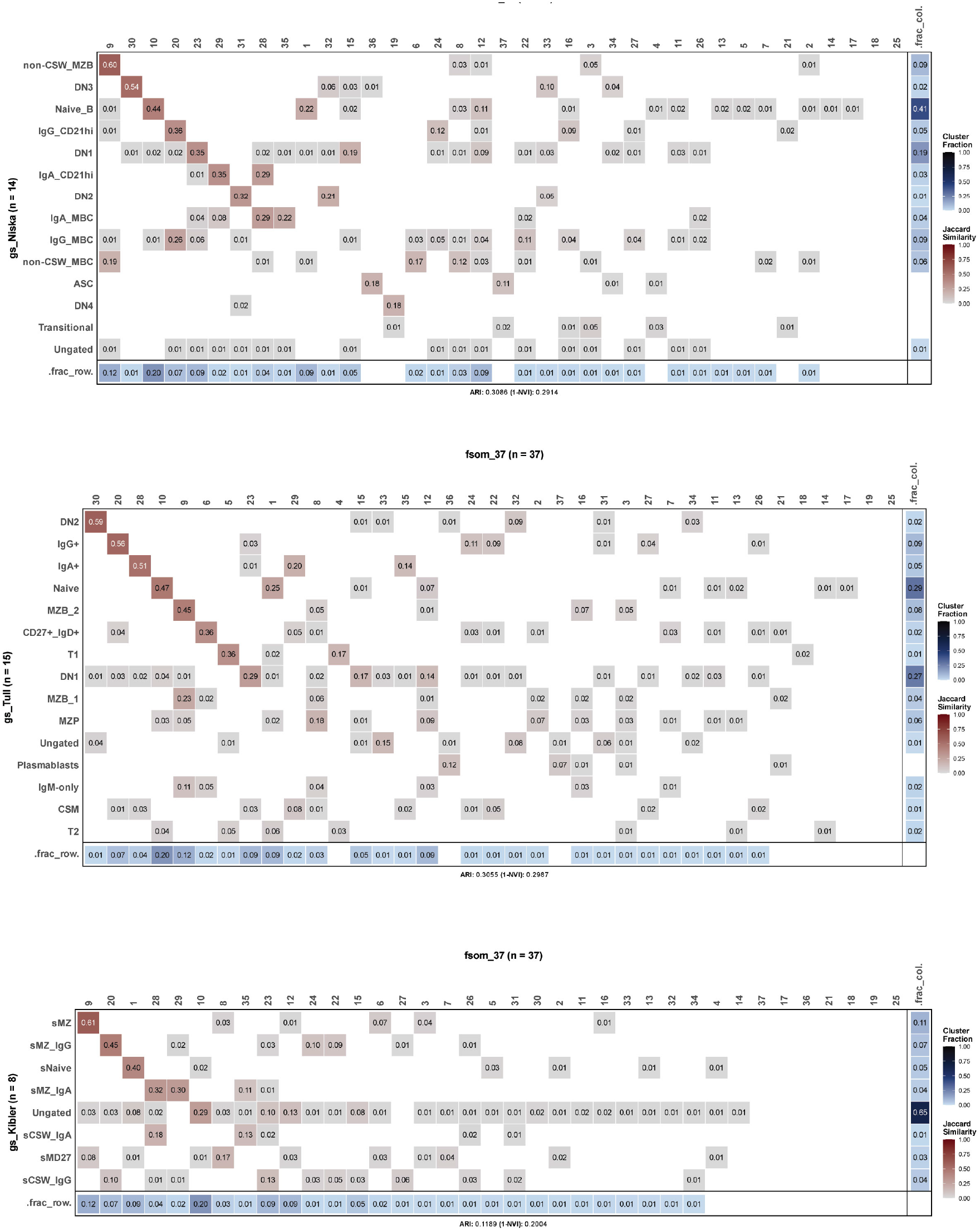
JSI for FlowSOM with 37 metaclusters (*k* = 37) versus gating schemes illustrated in **Supplemental Figure 3**.

**Figure 9d.**
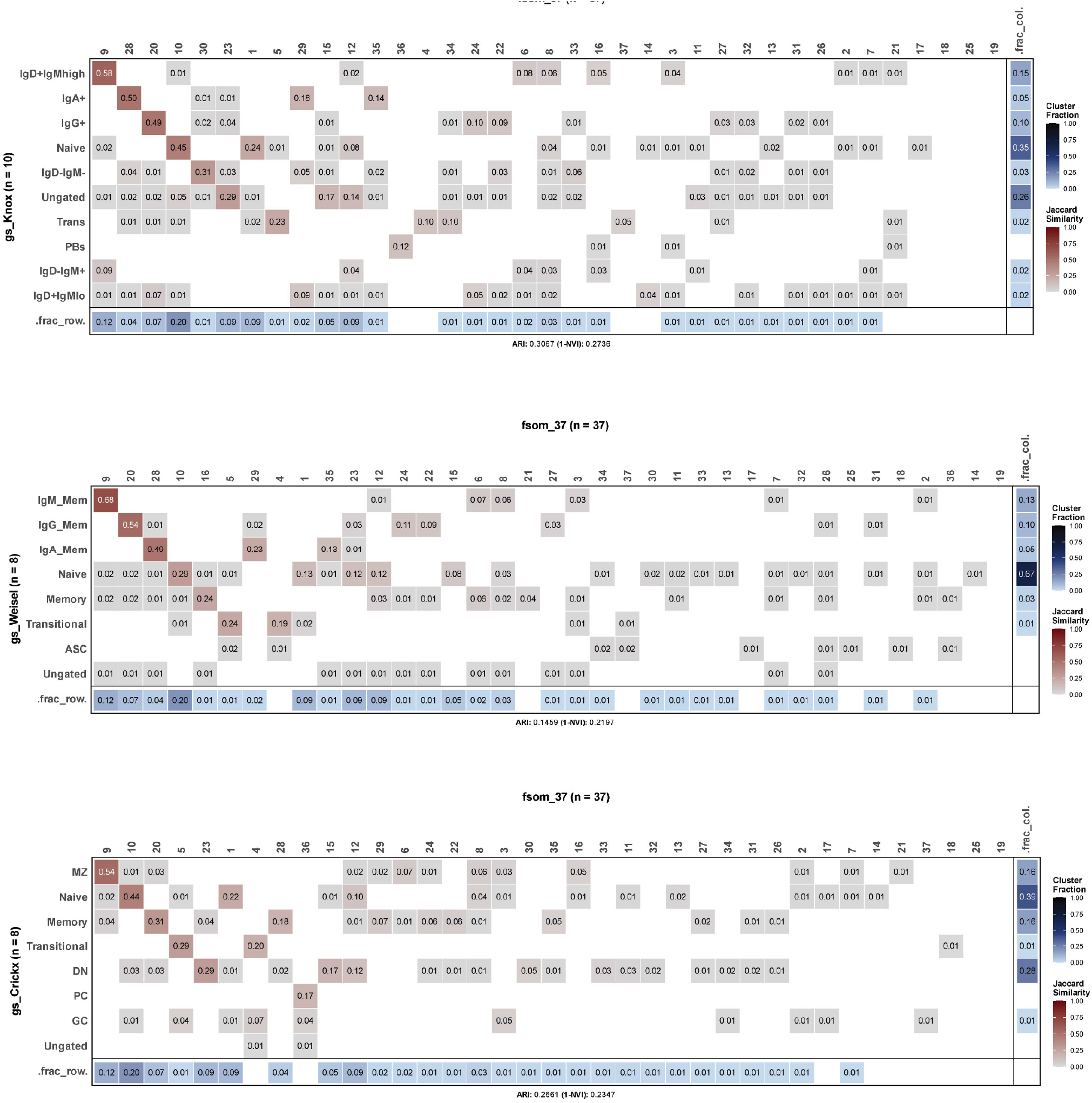
JSI for FlowSOM with 37 metaclusters (*k* = 37) versus gating schemes illustrated in **Supplemental Figure 3**.

**Figure 9e.**
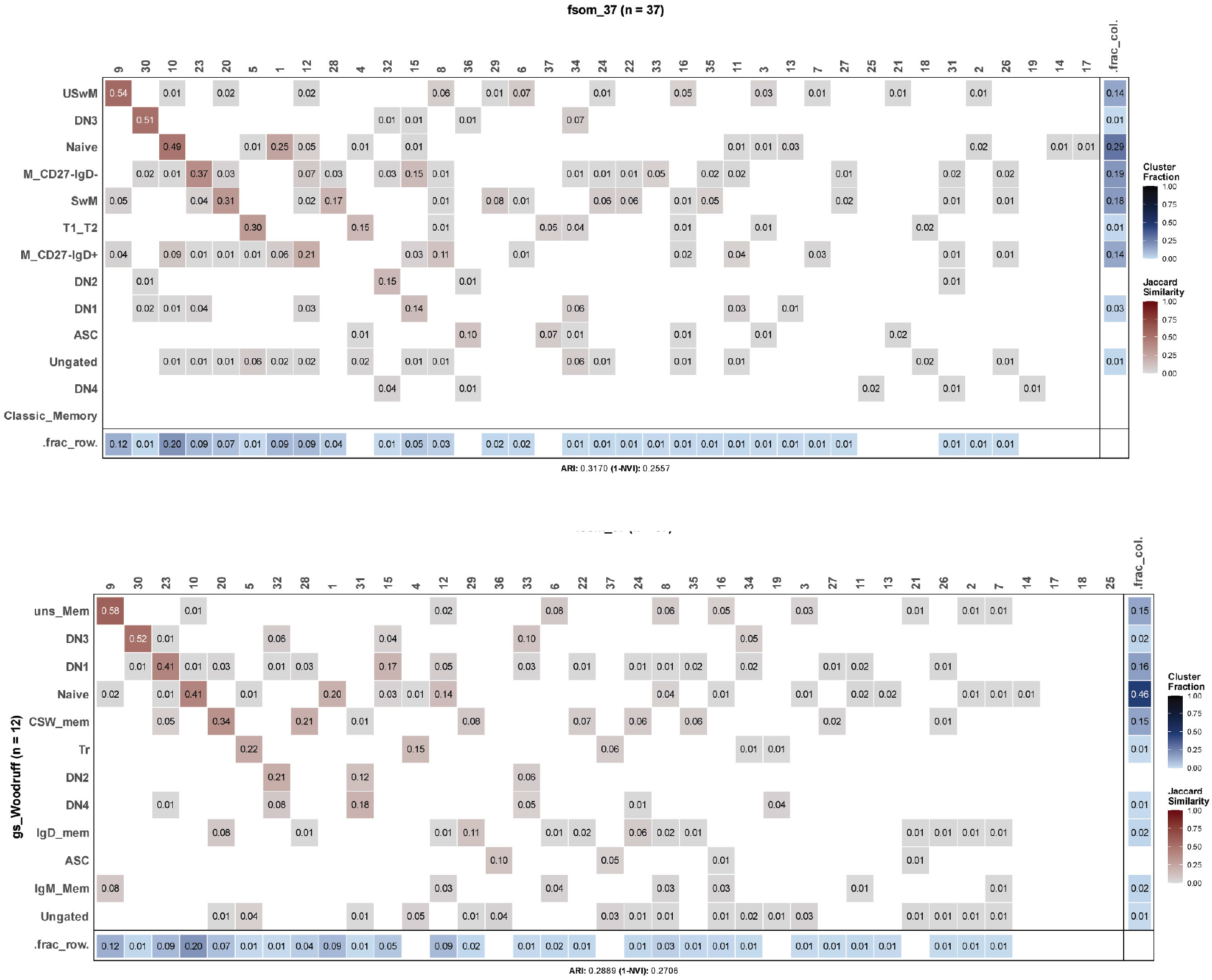
JSI for FlowSOM with 37 metaclusters (*k* = 37) versus gating schemes illustrated in **Supplemental Figure 3**.

**Figure 10a.**
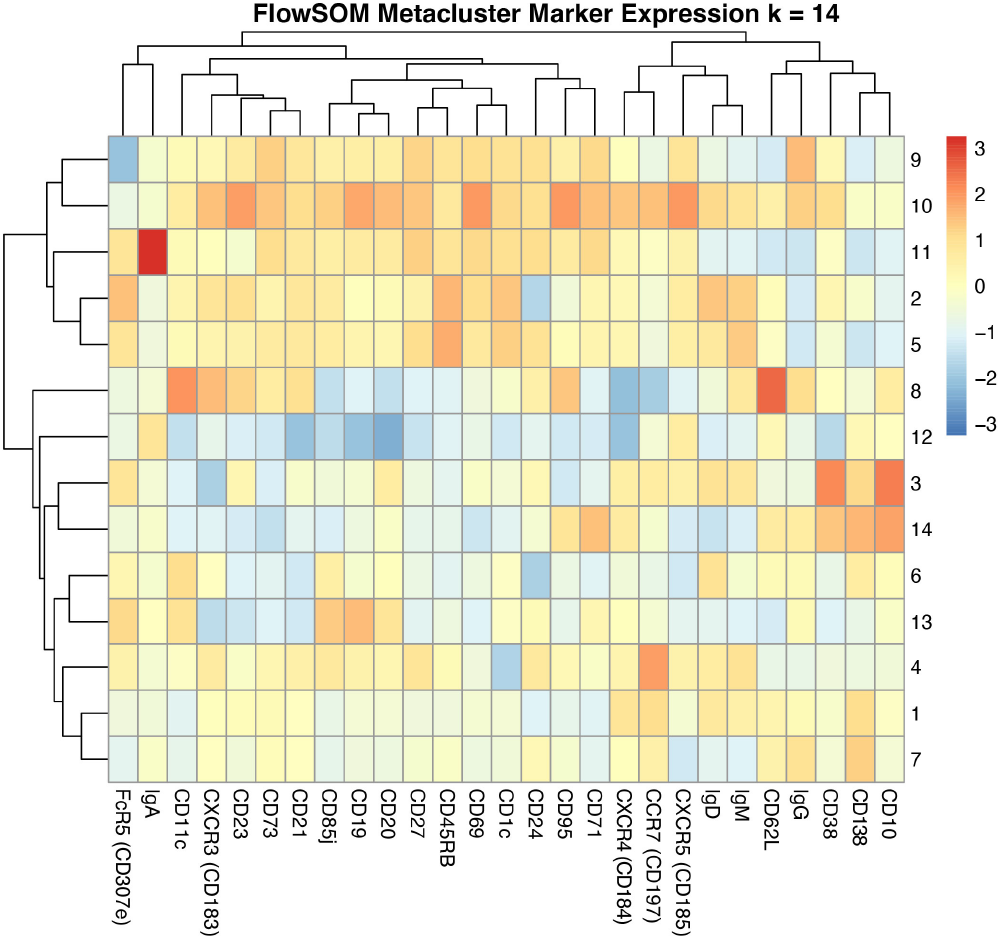
Heatmap of marker expression for metaclusters for *k* = 14

**Figure 10b.**
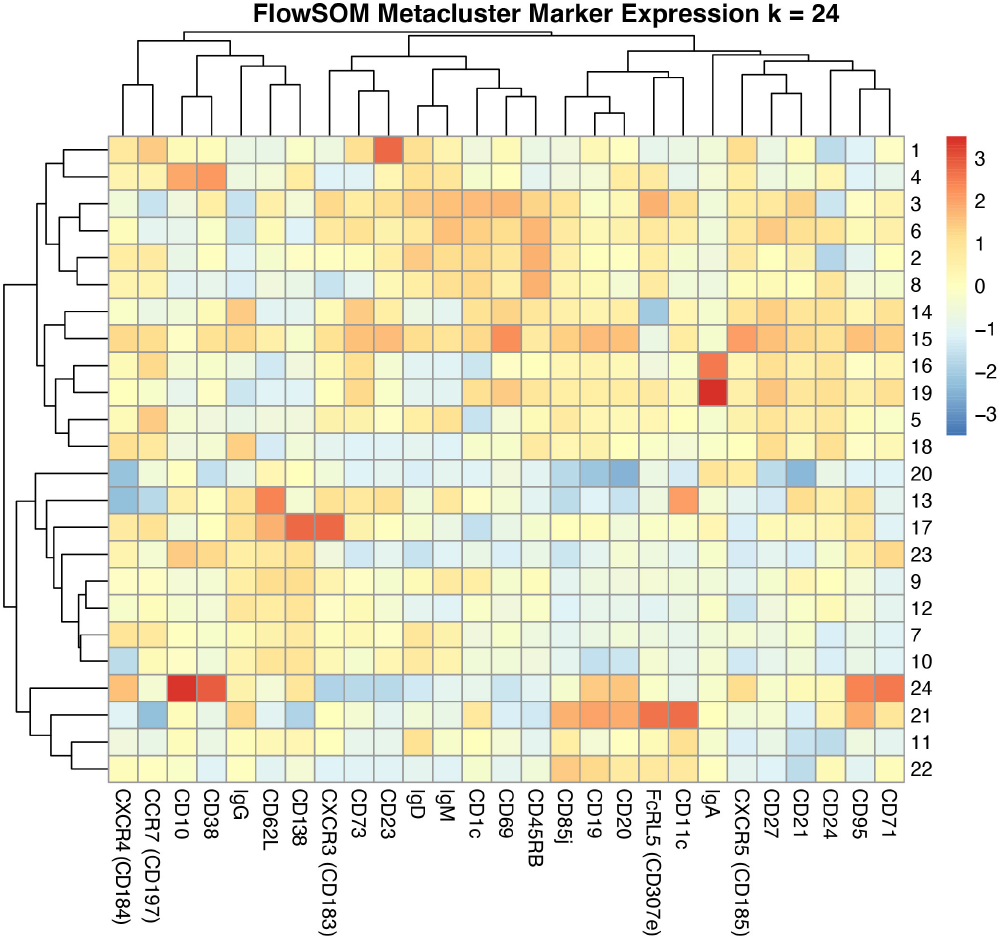
Heatmap of marker expression for metaclusters for *k* = 24

**Figure 10c.**
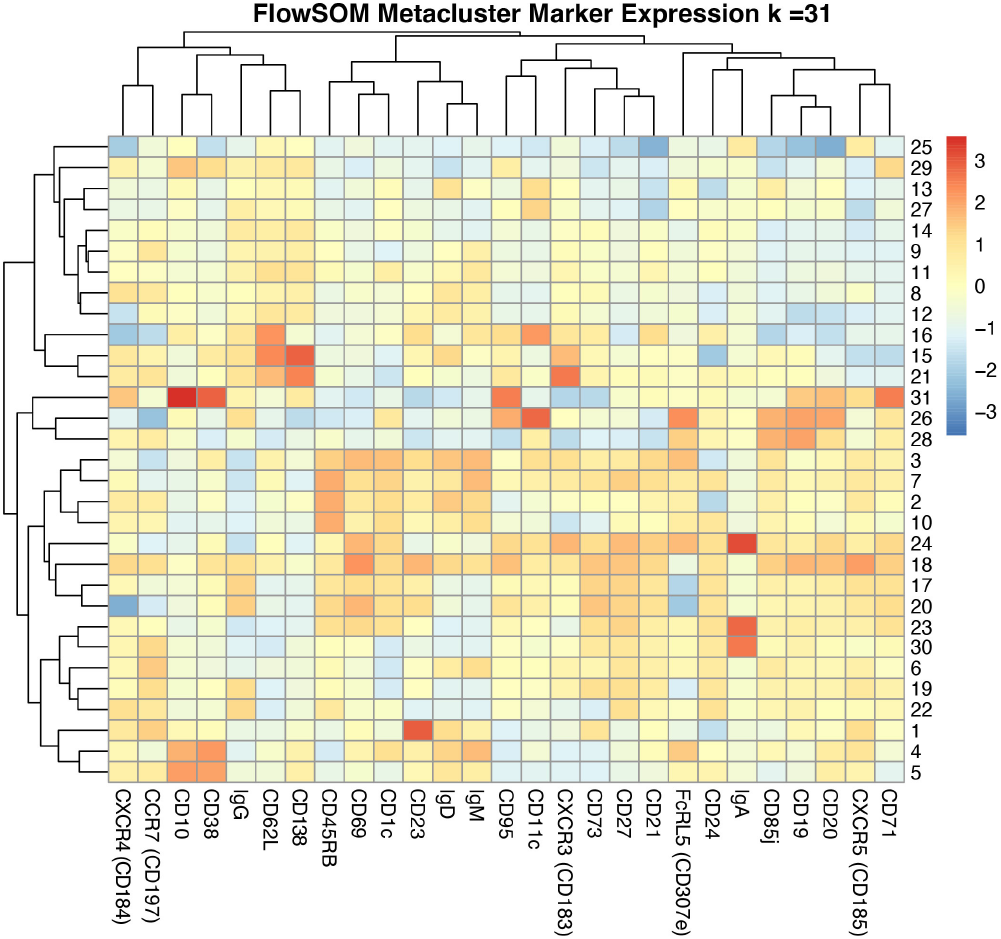
Heatmap of marker expression for metaclusters for *k* = 31

**Figure 10d.**
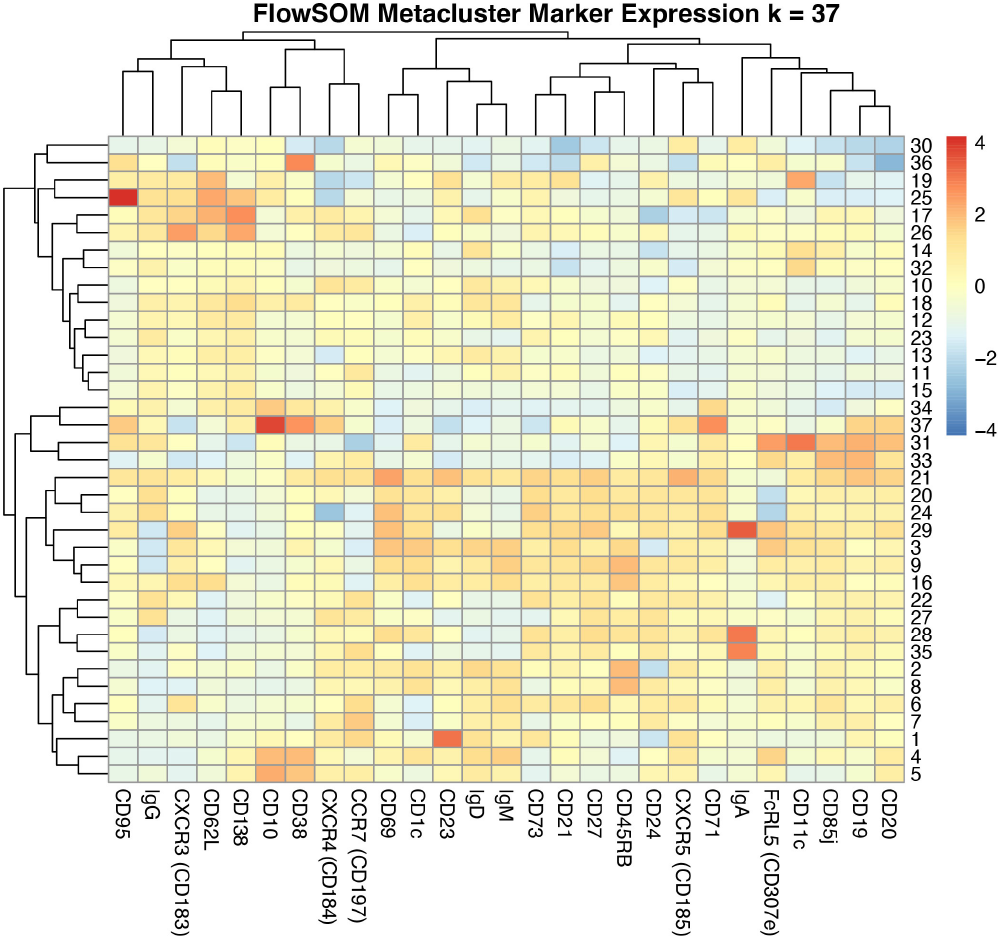
Heatmap of marker expression for metaclusters for *k* = 37

**Figure 11a.**
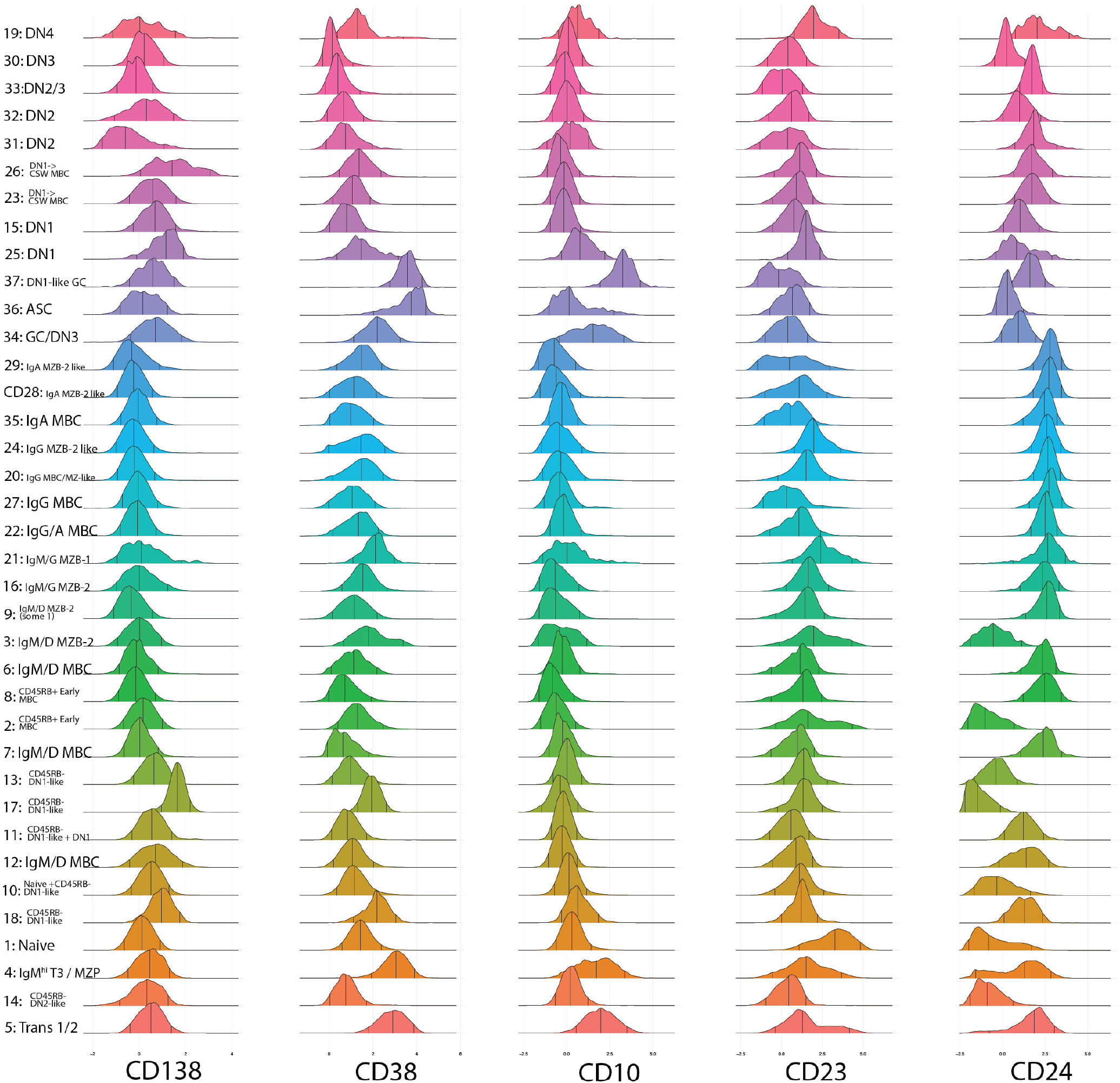
Flourescence expression density plots by metacluster for *k* = 37

**Figure 11b.**
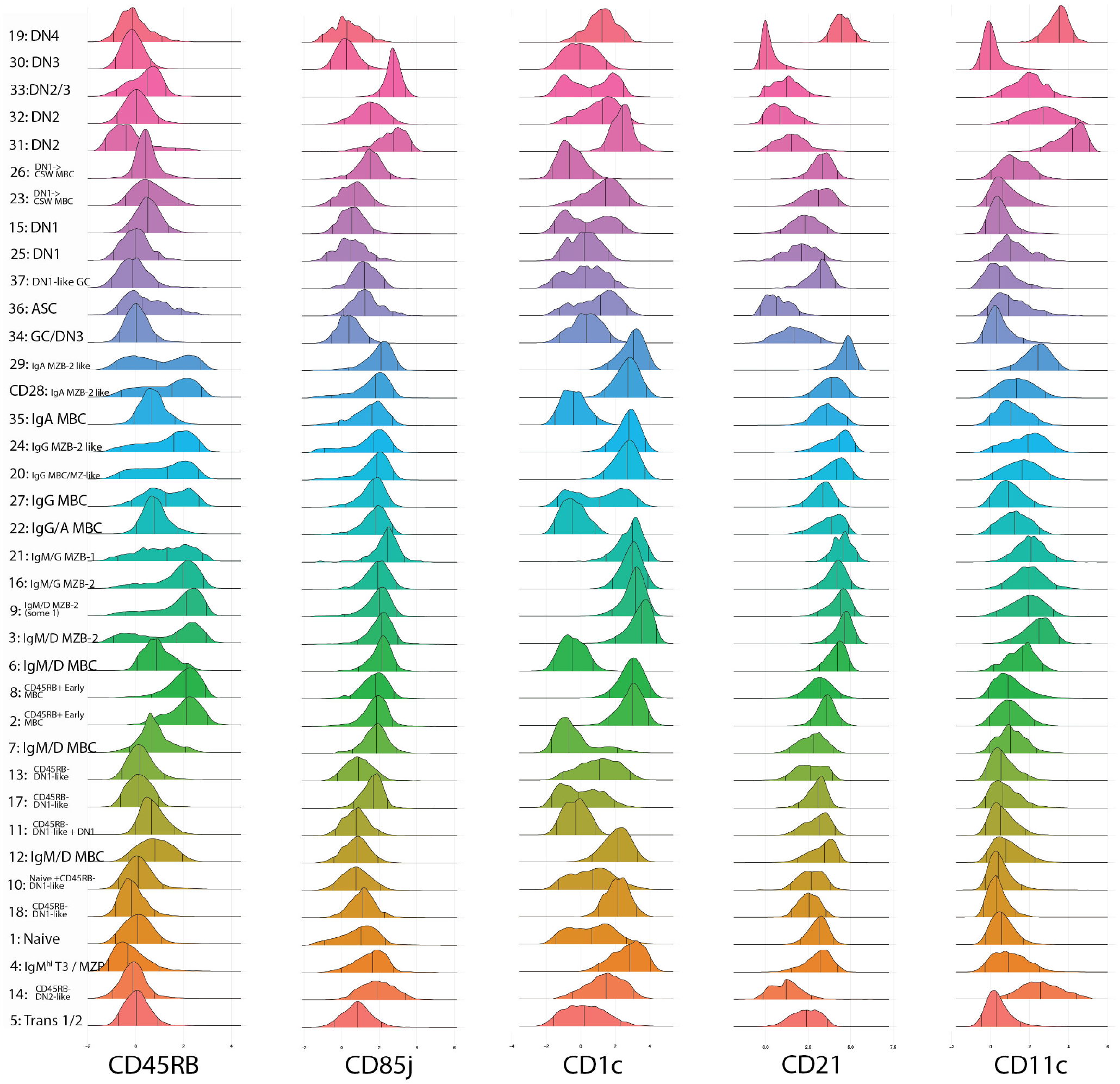
Flourescence expression density plots by metacluster for *k* = 37

**Figure 11c.**
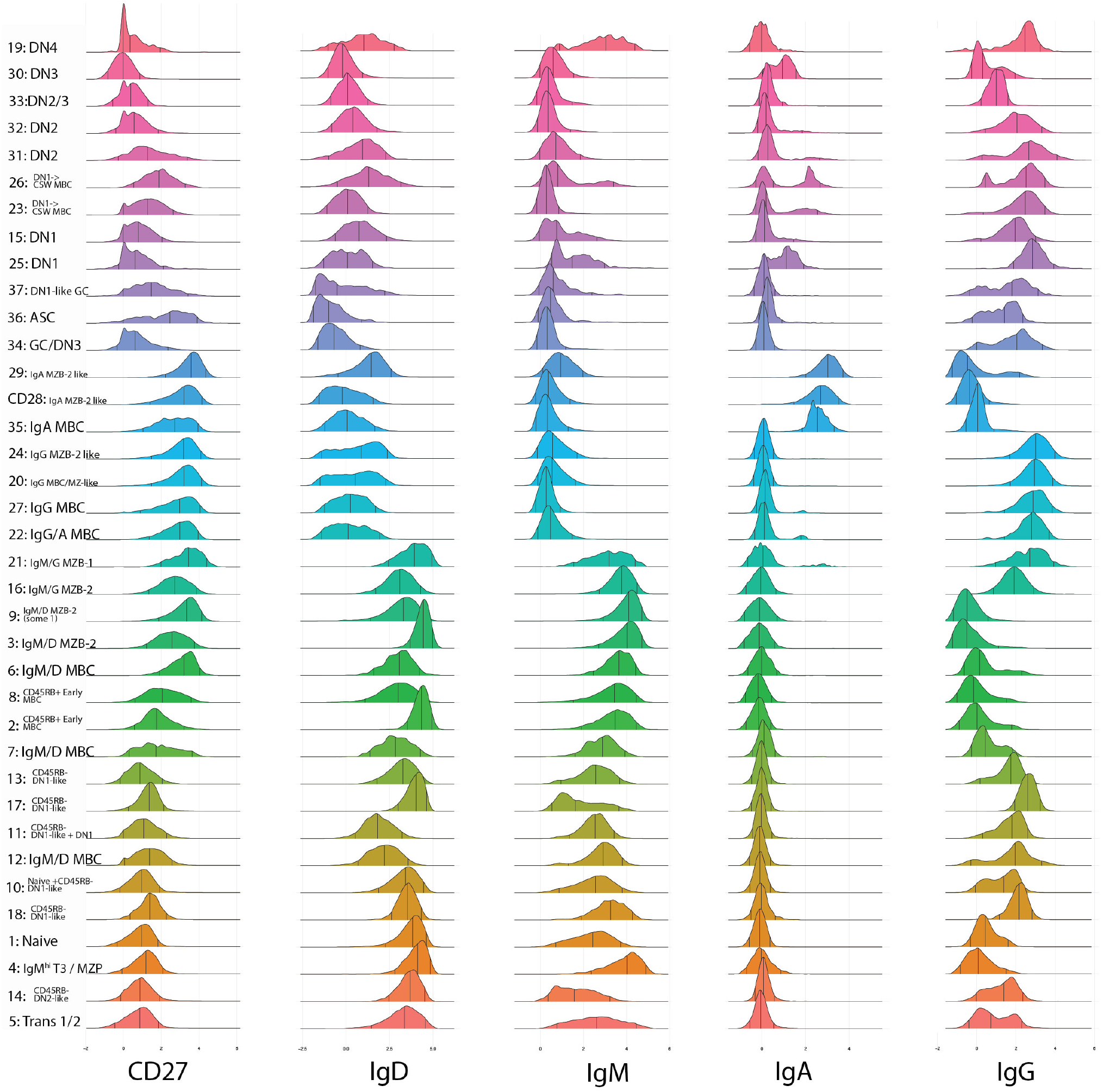
Flourescence expression density plots by metacluster for *k* = 37

**Figure 11d.**
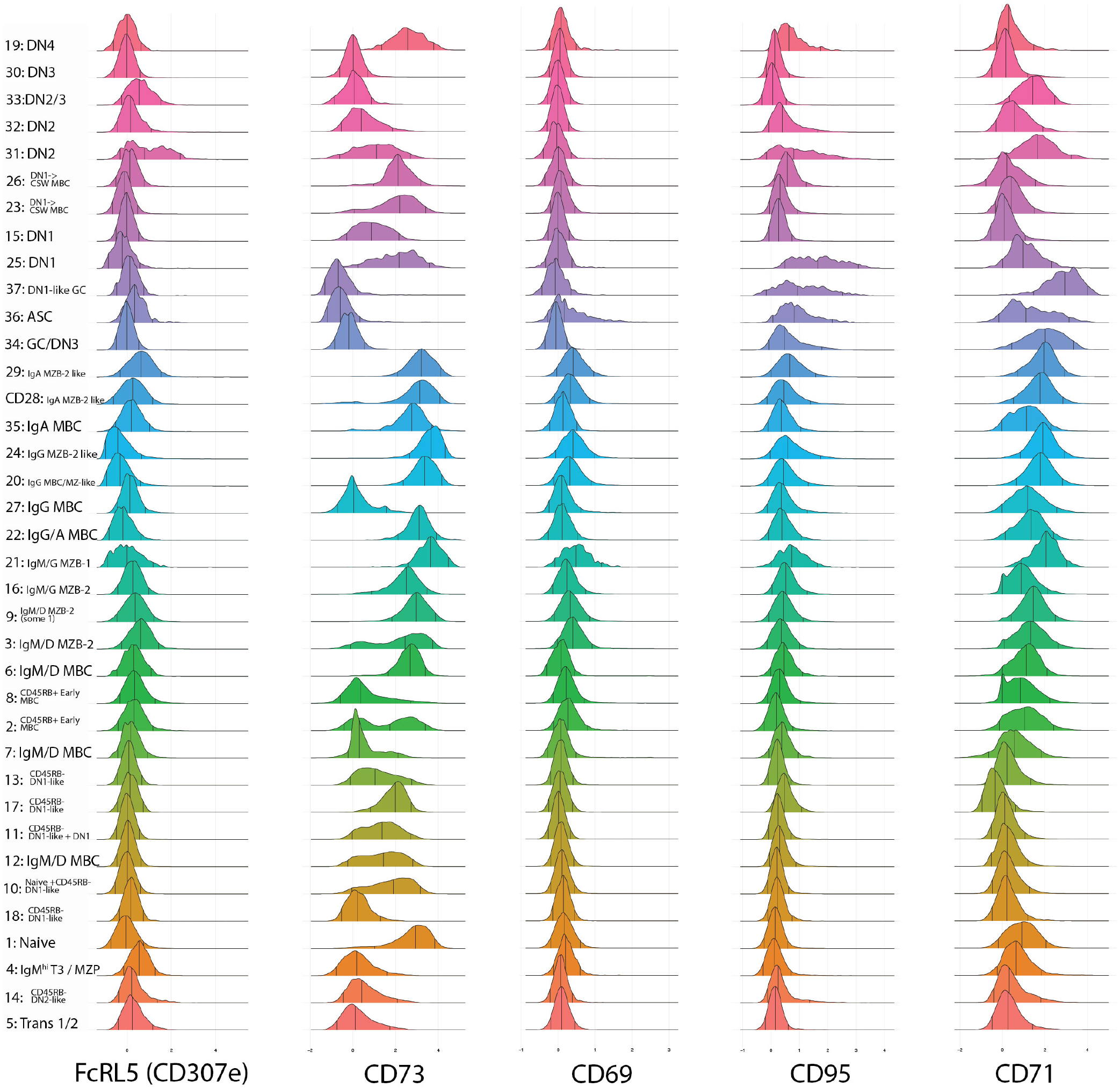
Flourescence expression density plots by metacluster for *k* = 37

**Figure 11e.**
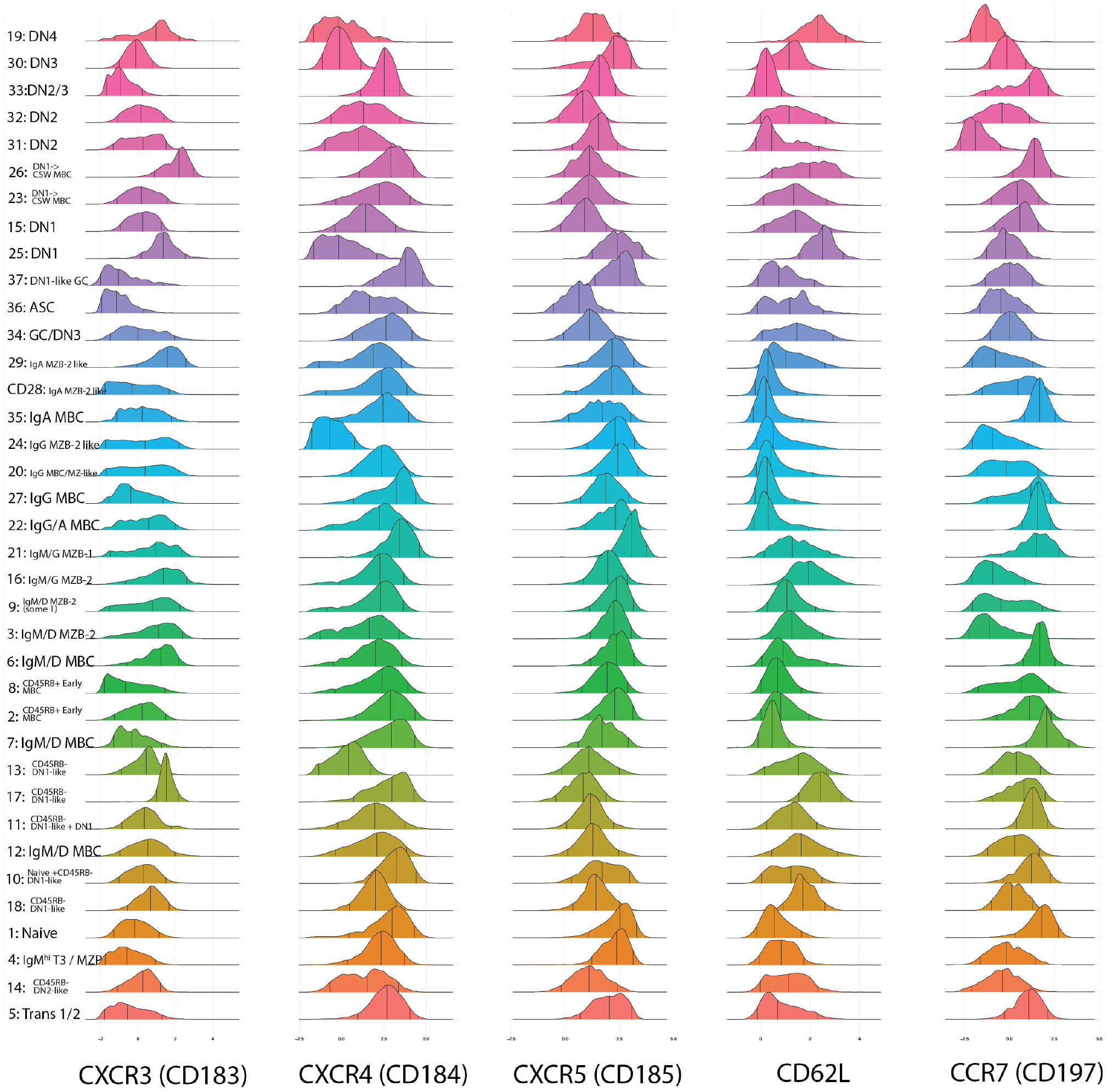
Flourescence expression density plots by metacluster for *k* = 37

**Figure 11f.**
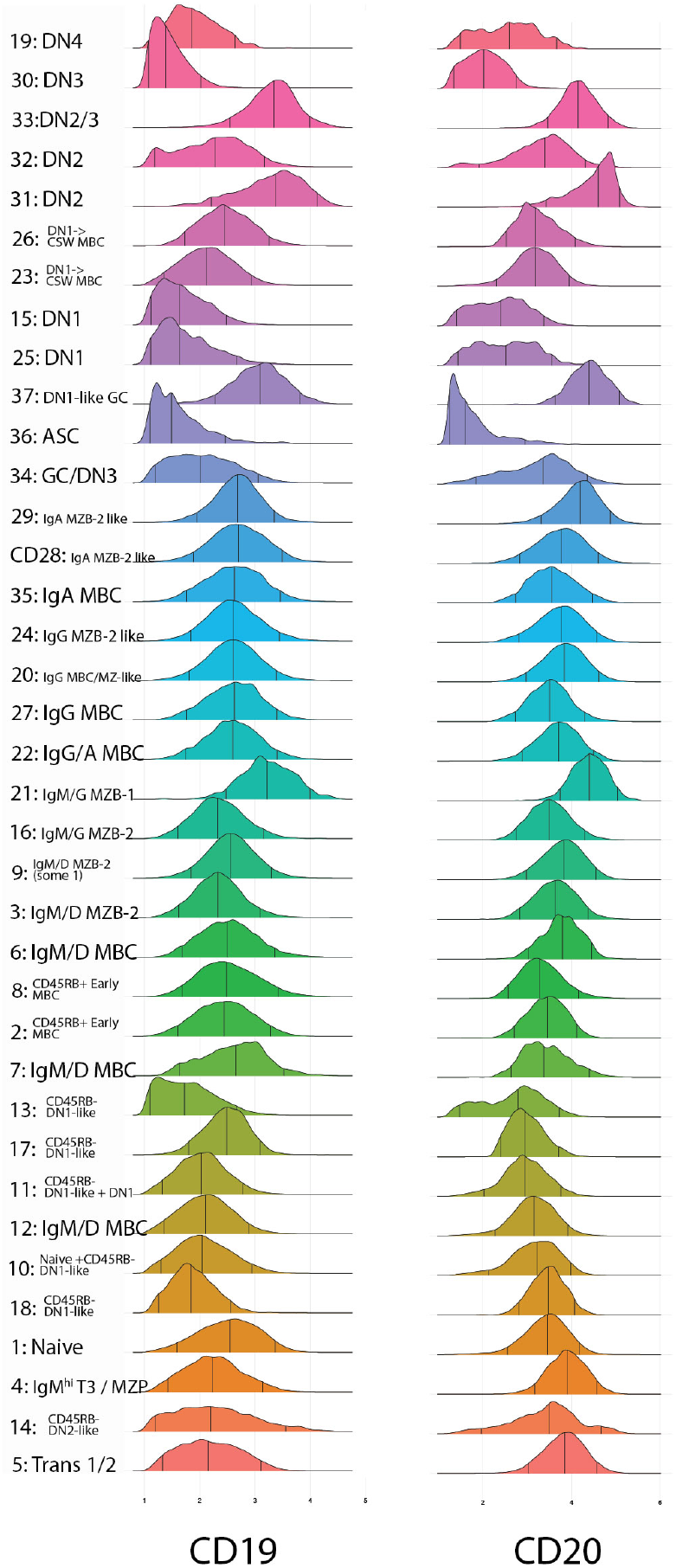
Flourescence expression density plots by metacluster for *k* = 37

**Figure 12a.**
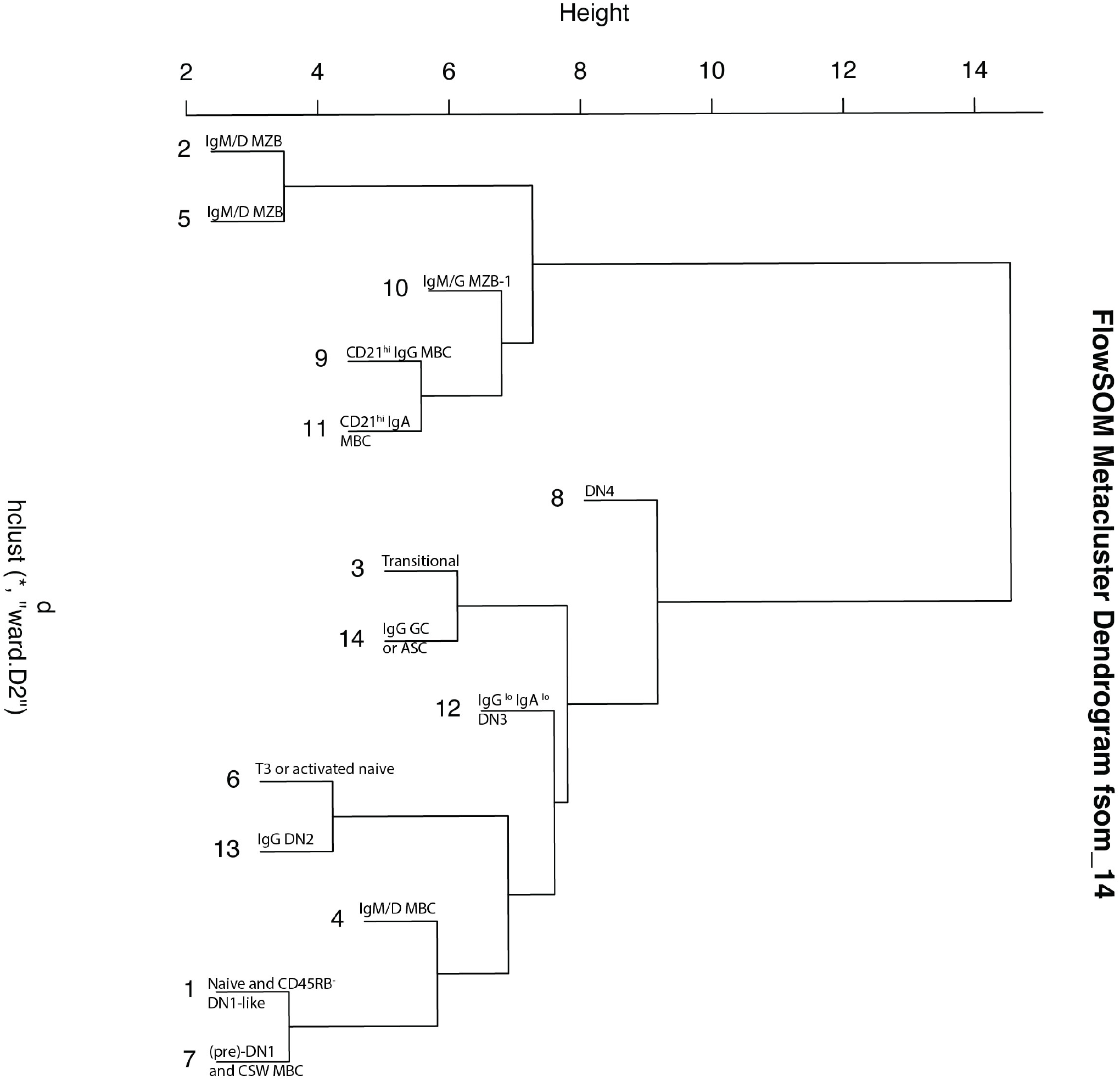
Dendrogram for *k* = 14 with metacluster label.

**Figure 12b.**
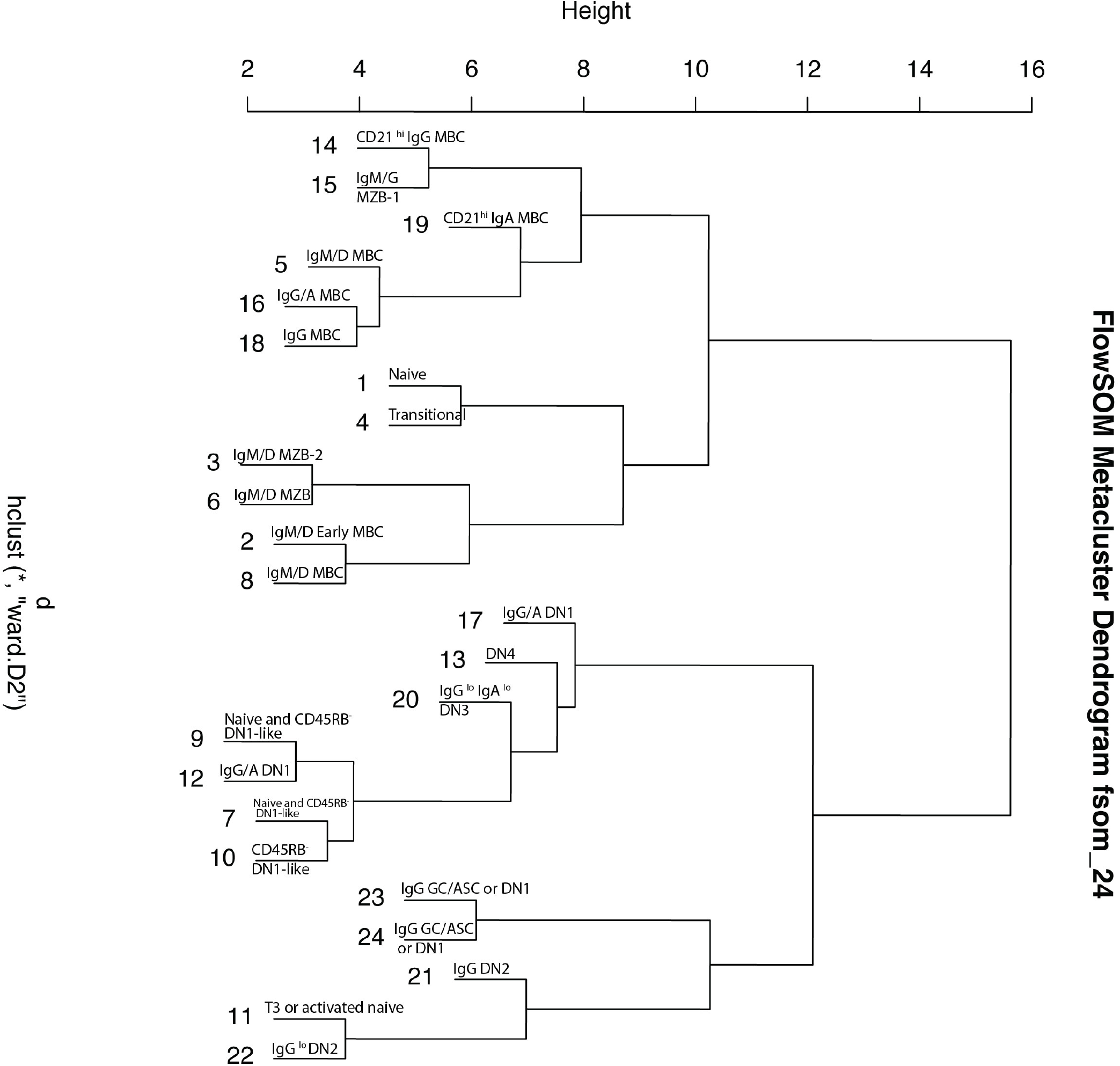
Dendrogram for *k* = 24 with metacluster label.

**Figure 12c.**
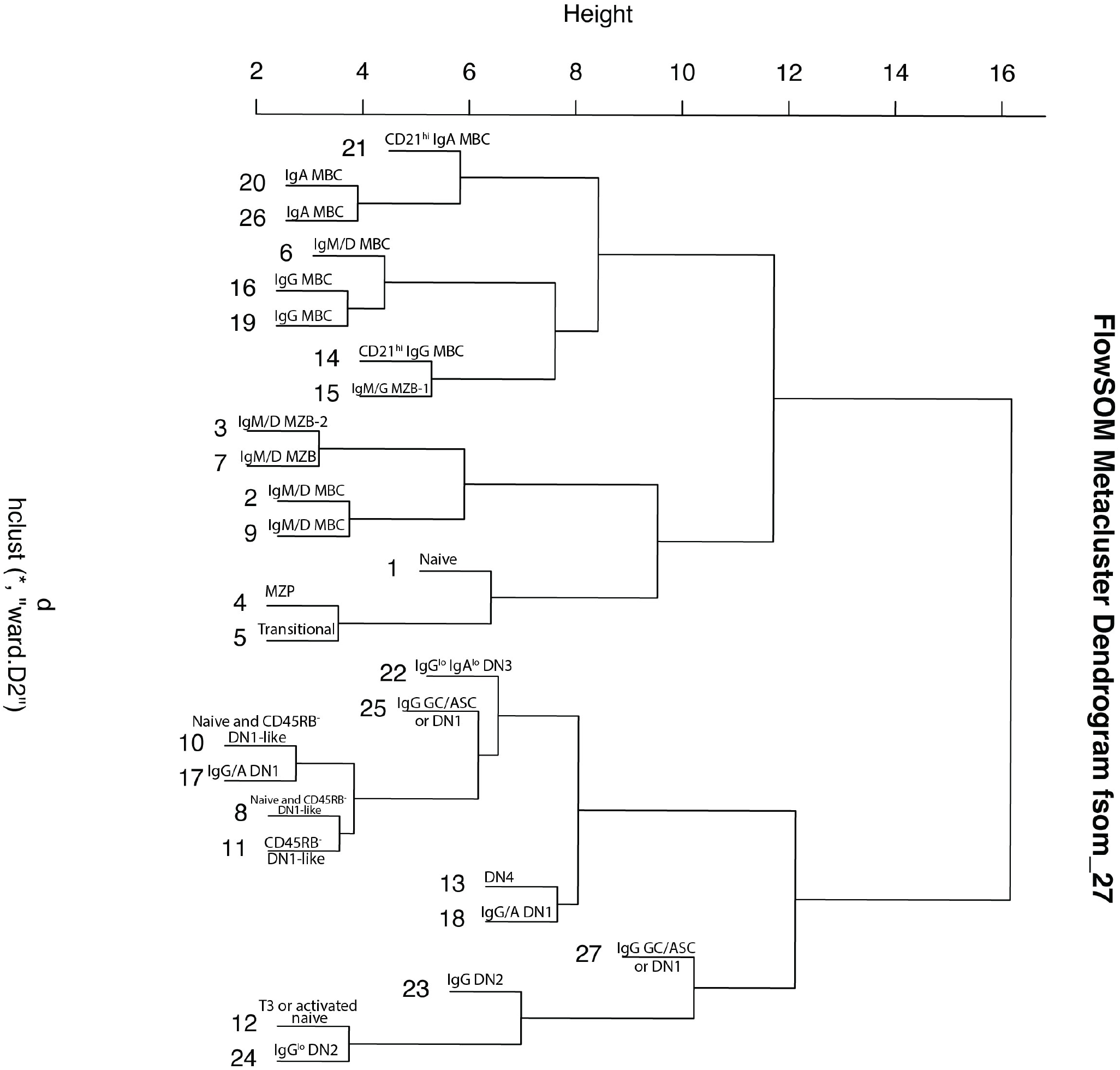
Dendrogram for *k* = 27 with metacluster label.

**Figure 12d.**
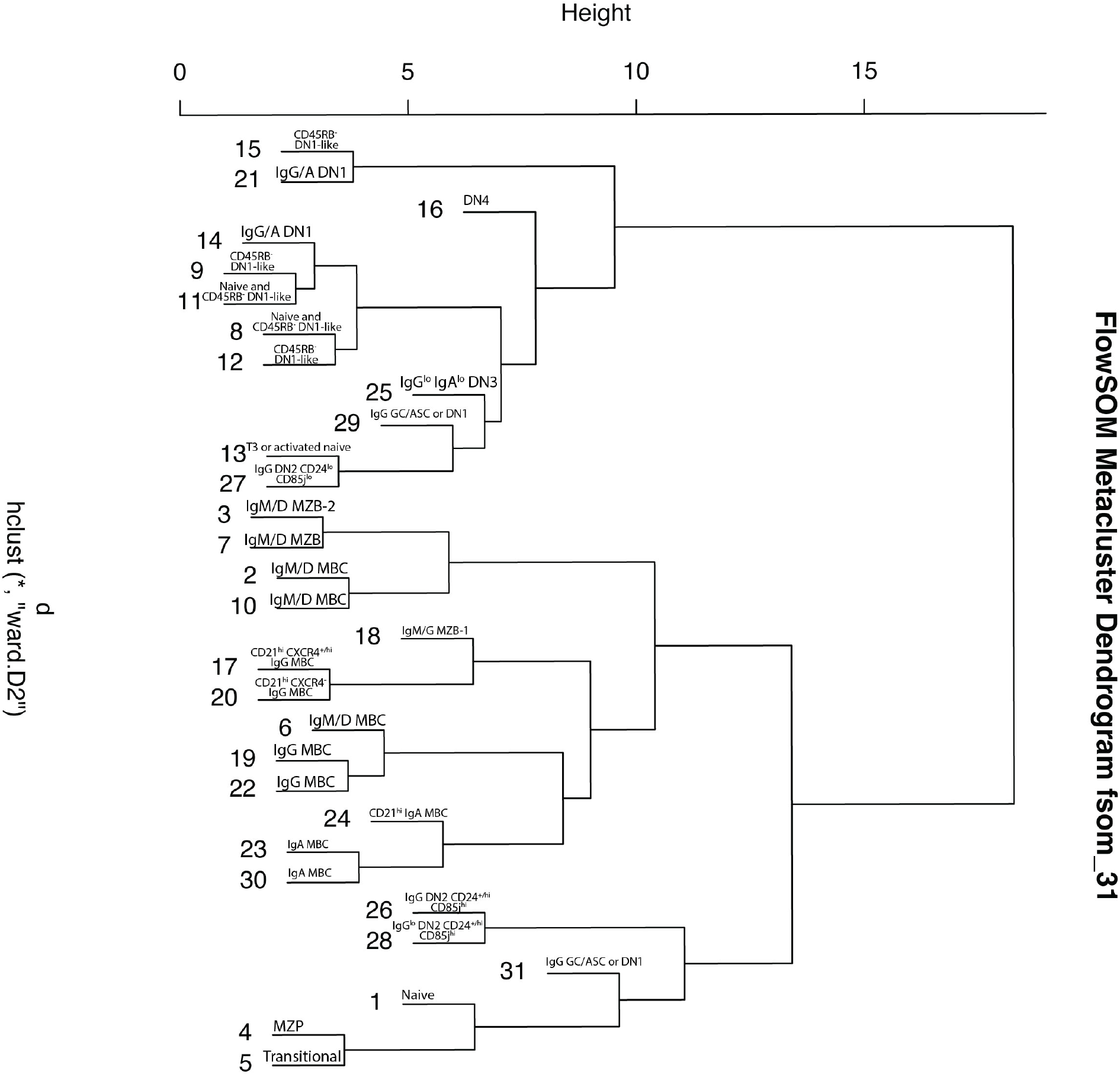
Dendrogram for *k* = 31 with metacluster label.

**Figure 12e.**
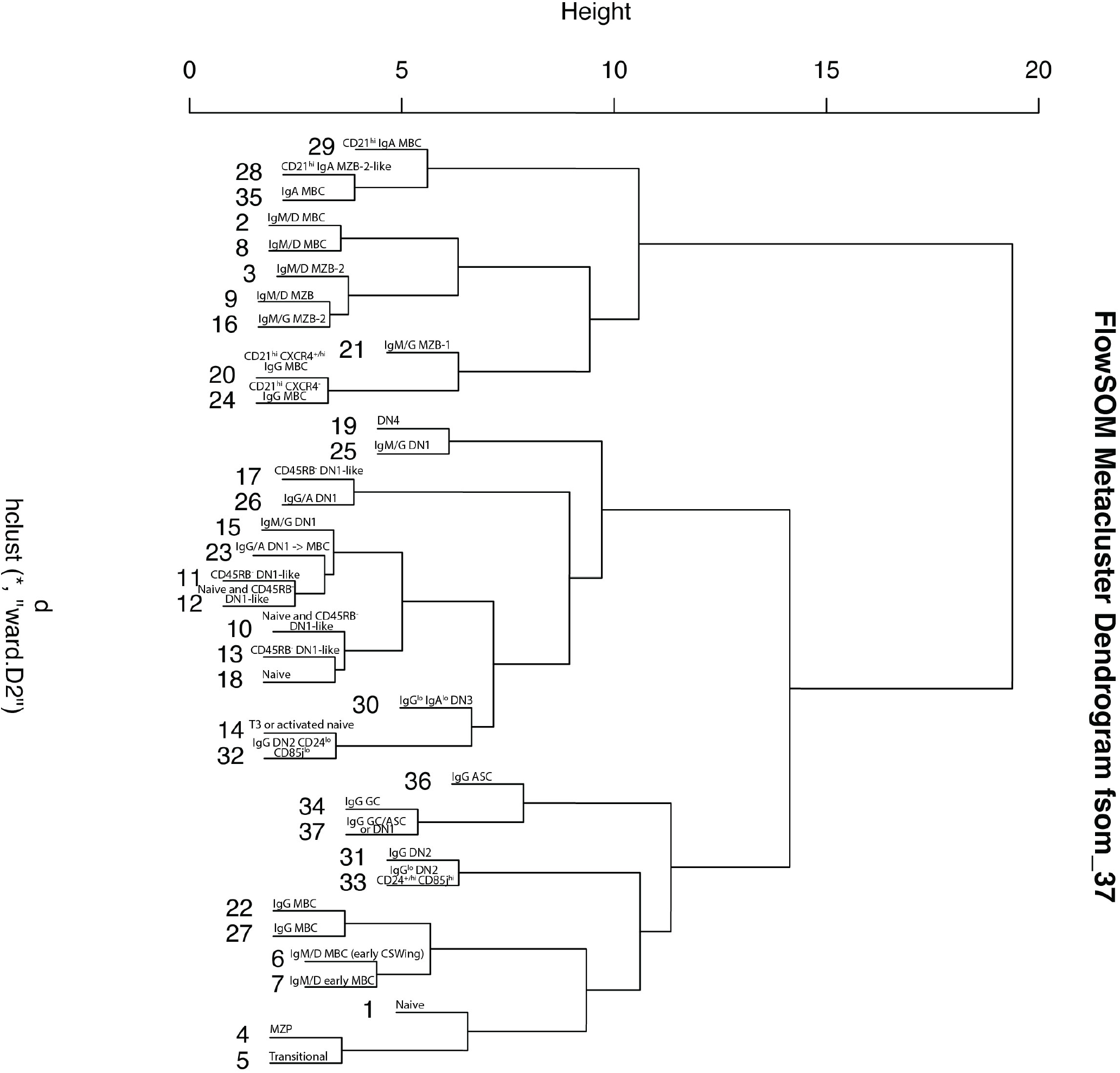
Dendrogram for *k* = 37 with metacluster label.

**Figure S13.**
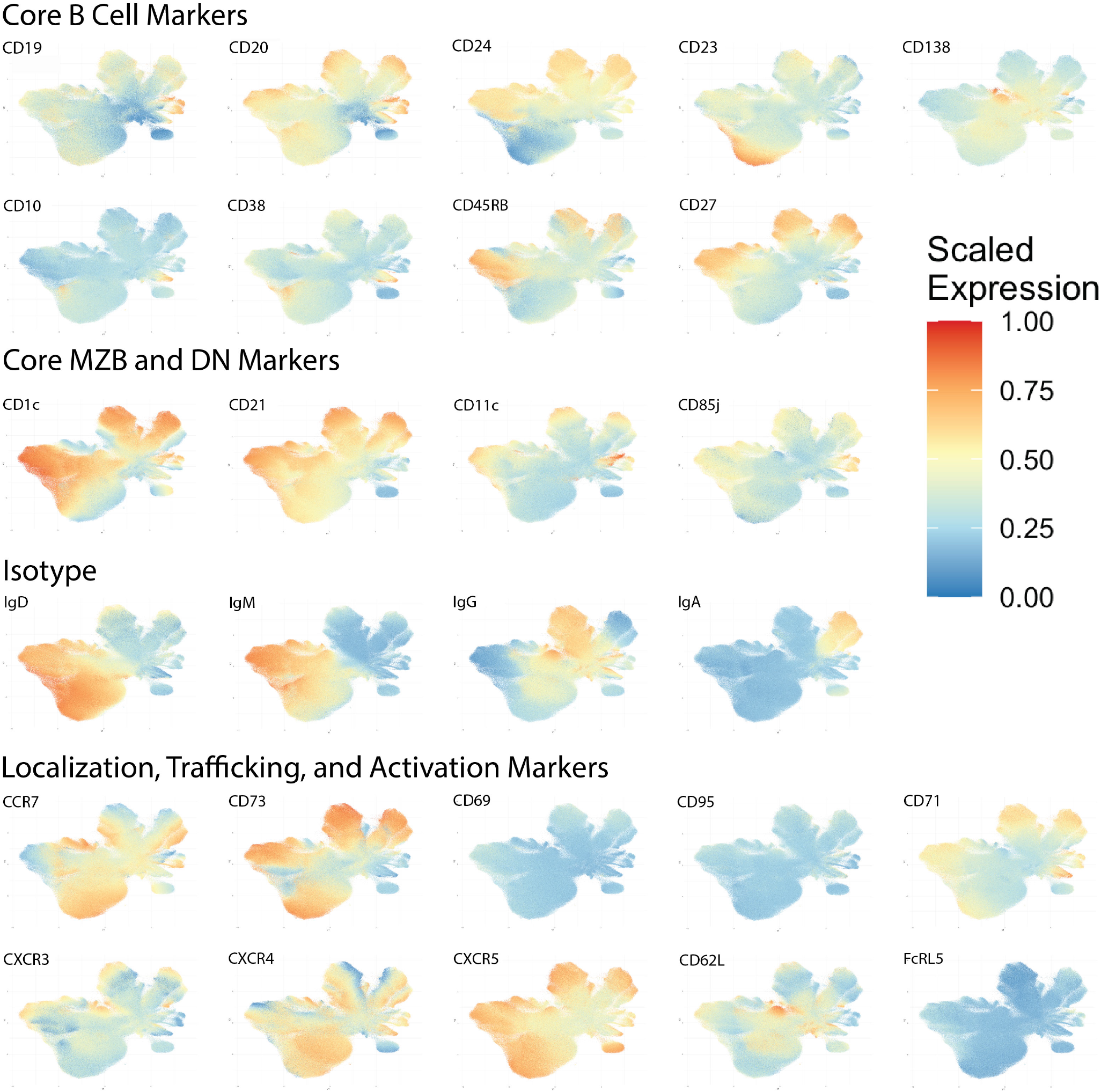
Fluorescent marker expression projected with a heatmap scale onto the UMAP from Figure 6

**Figure 14a.**
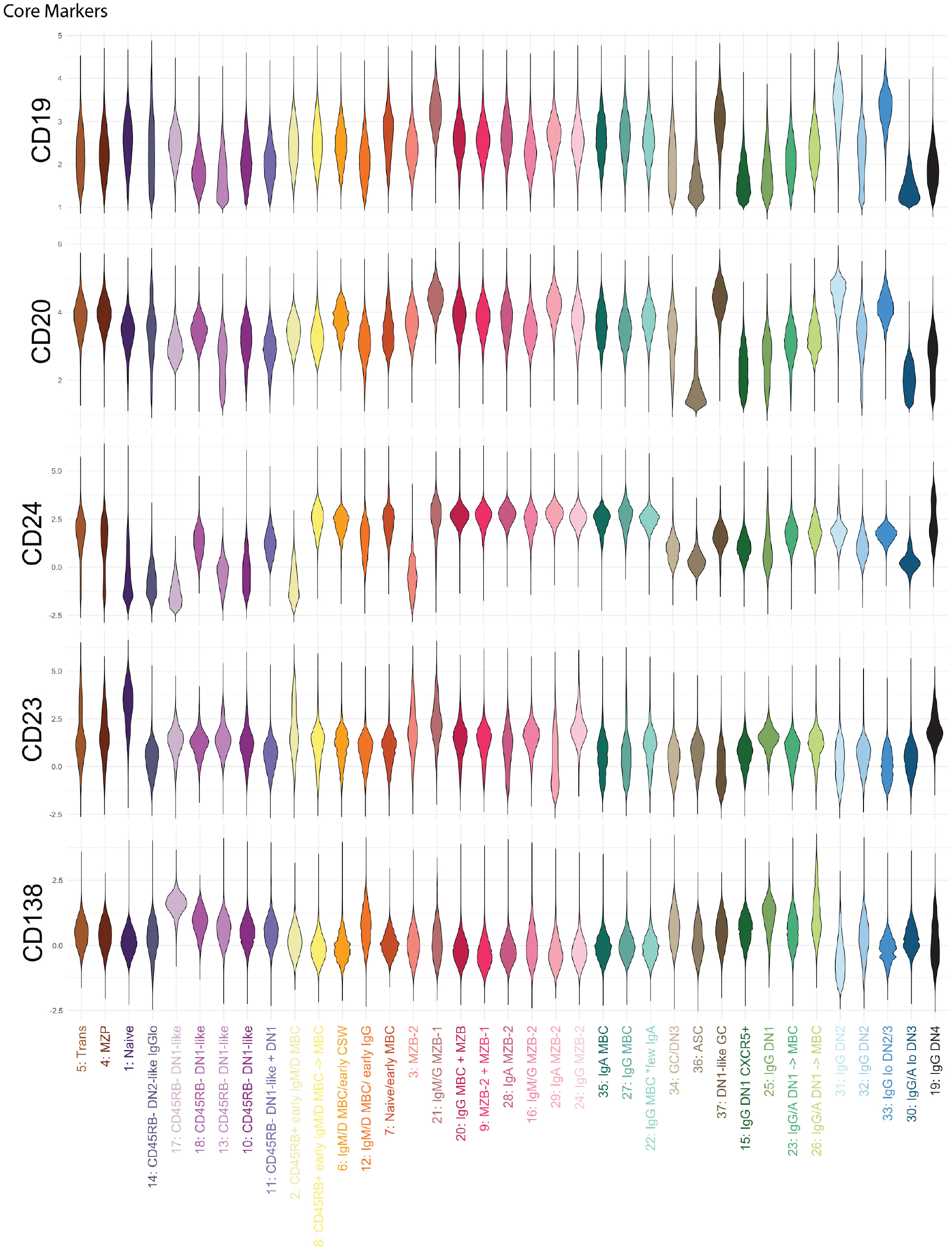
Violin plots of marker expression by cluster for k=37.

**Figure 14b.**
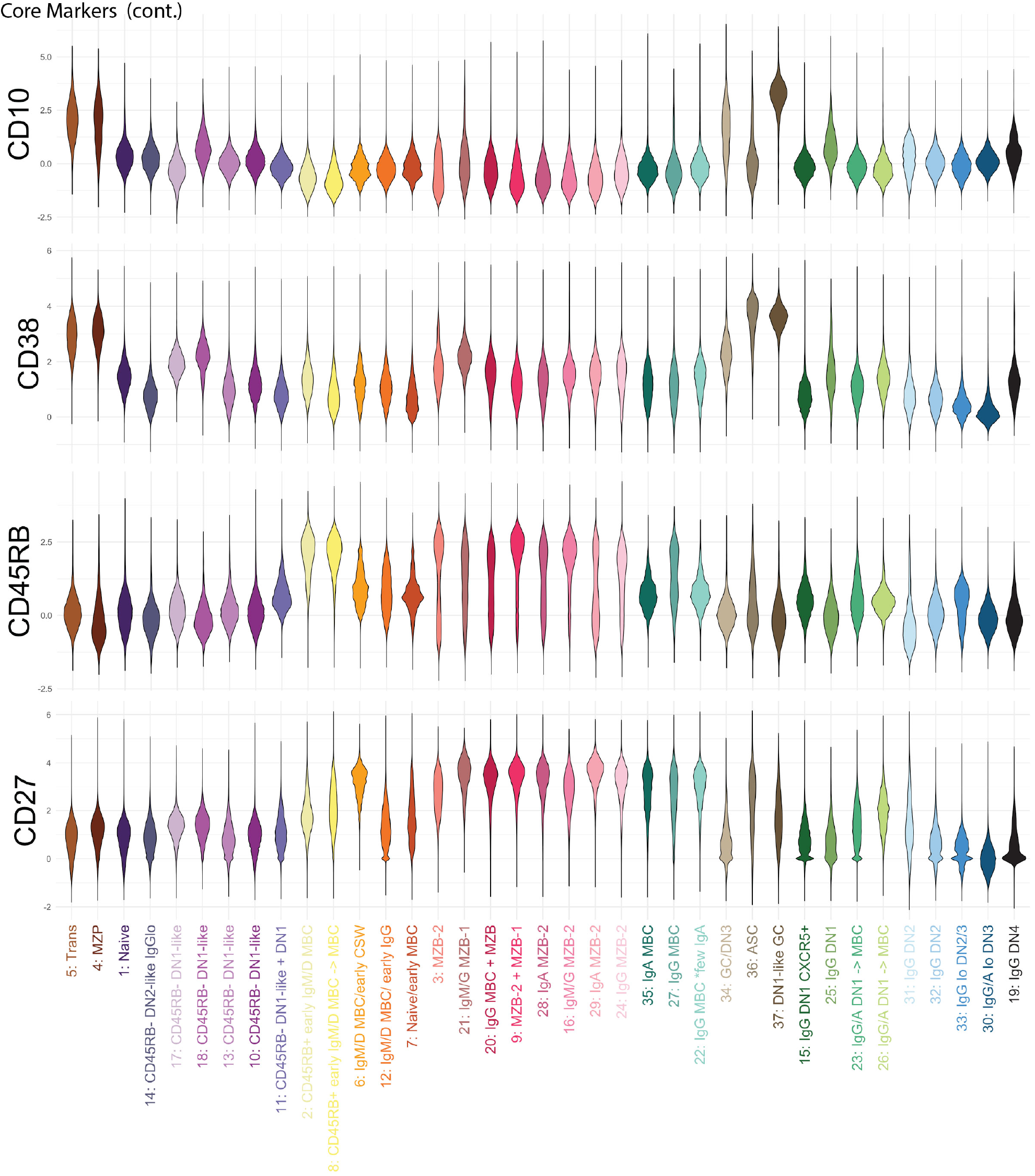
Violin plots of marker expression by cluster for k=37.

**Figure 14c.**
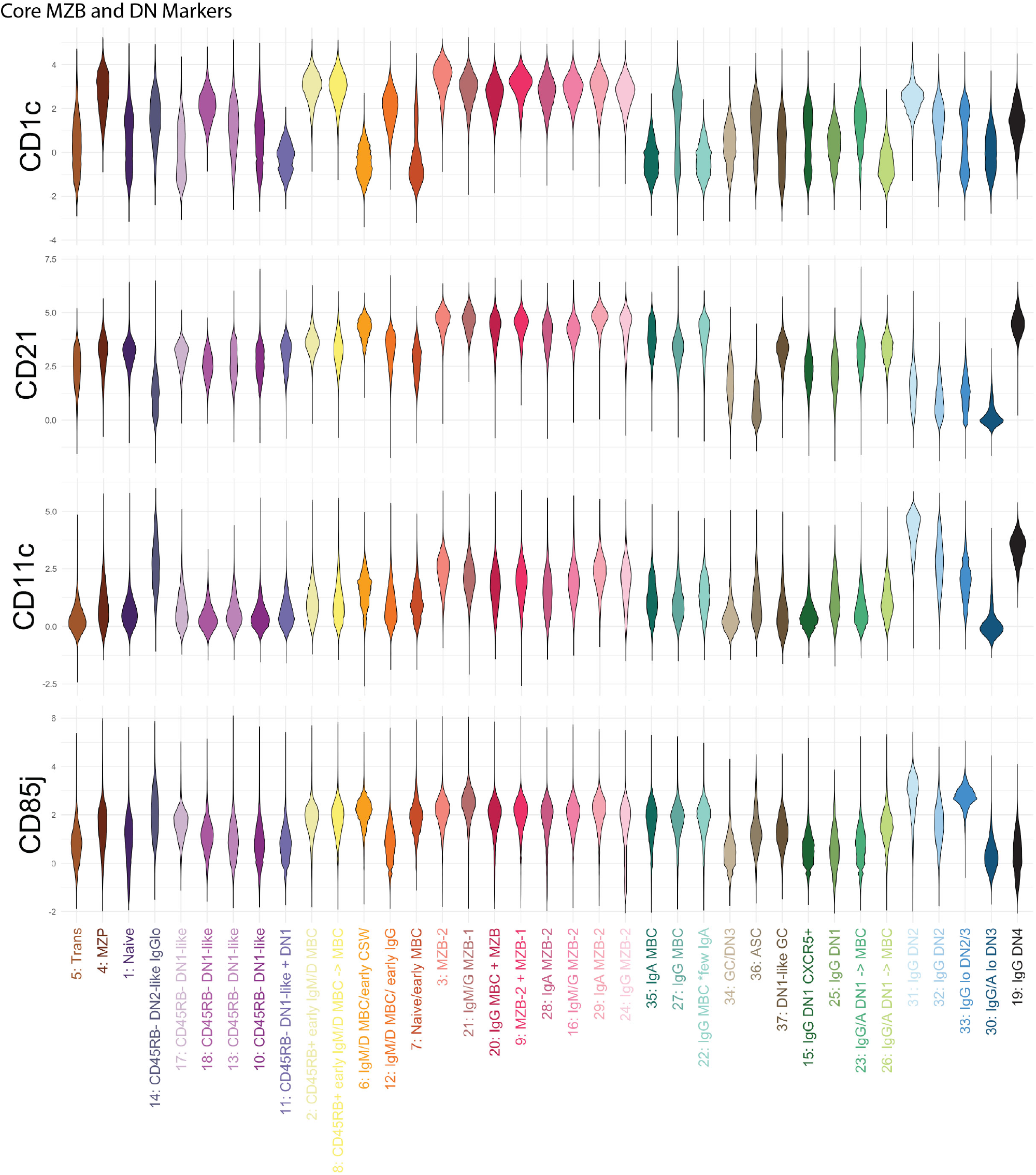
Violin plots of marker expression by cluster for k=37.

**Figure 14d.**
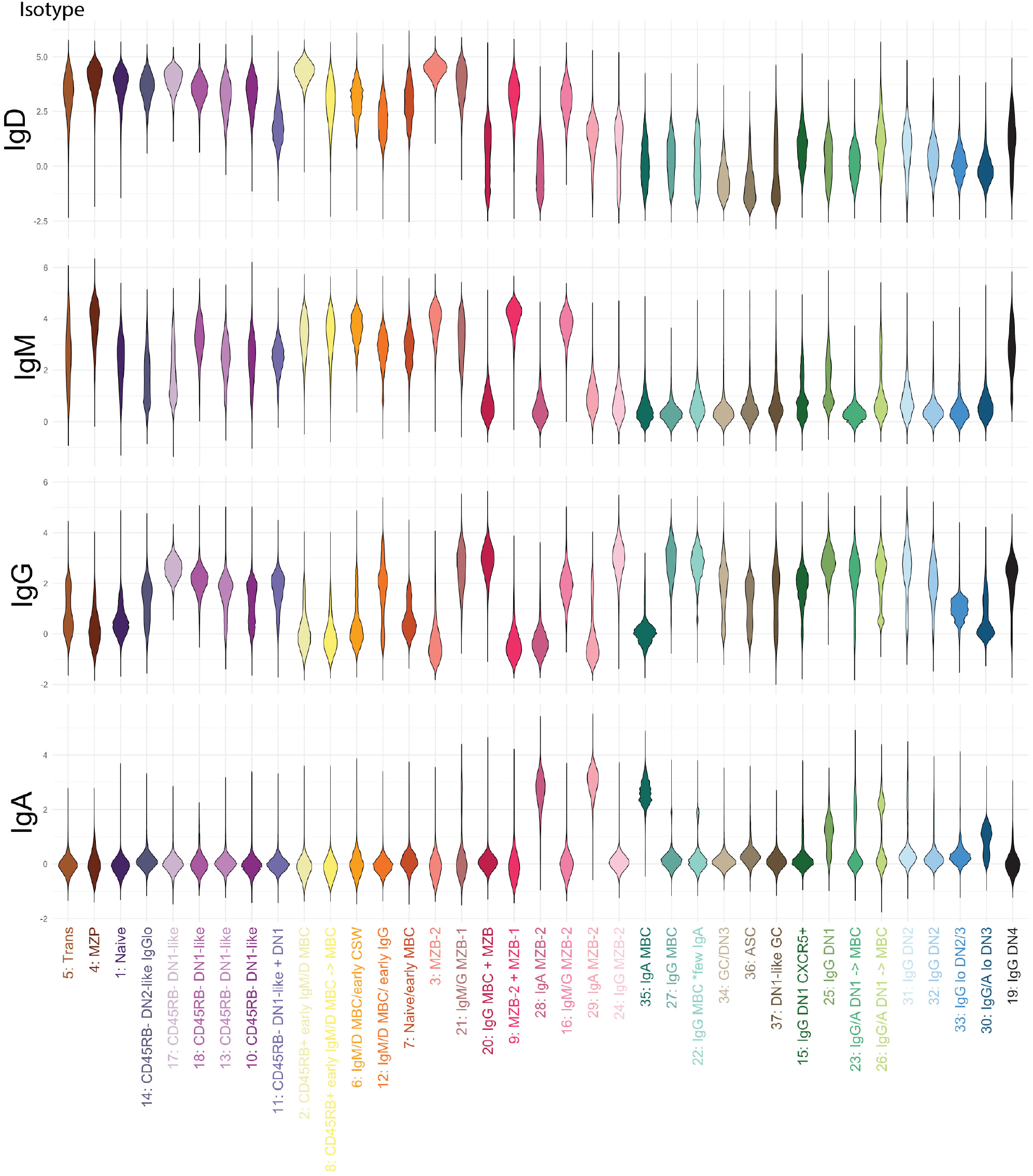
Violin plots of marker expression by cluster for k=37.

**Figure 14e.**
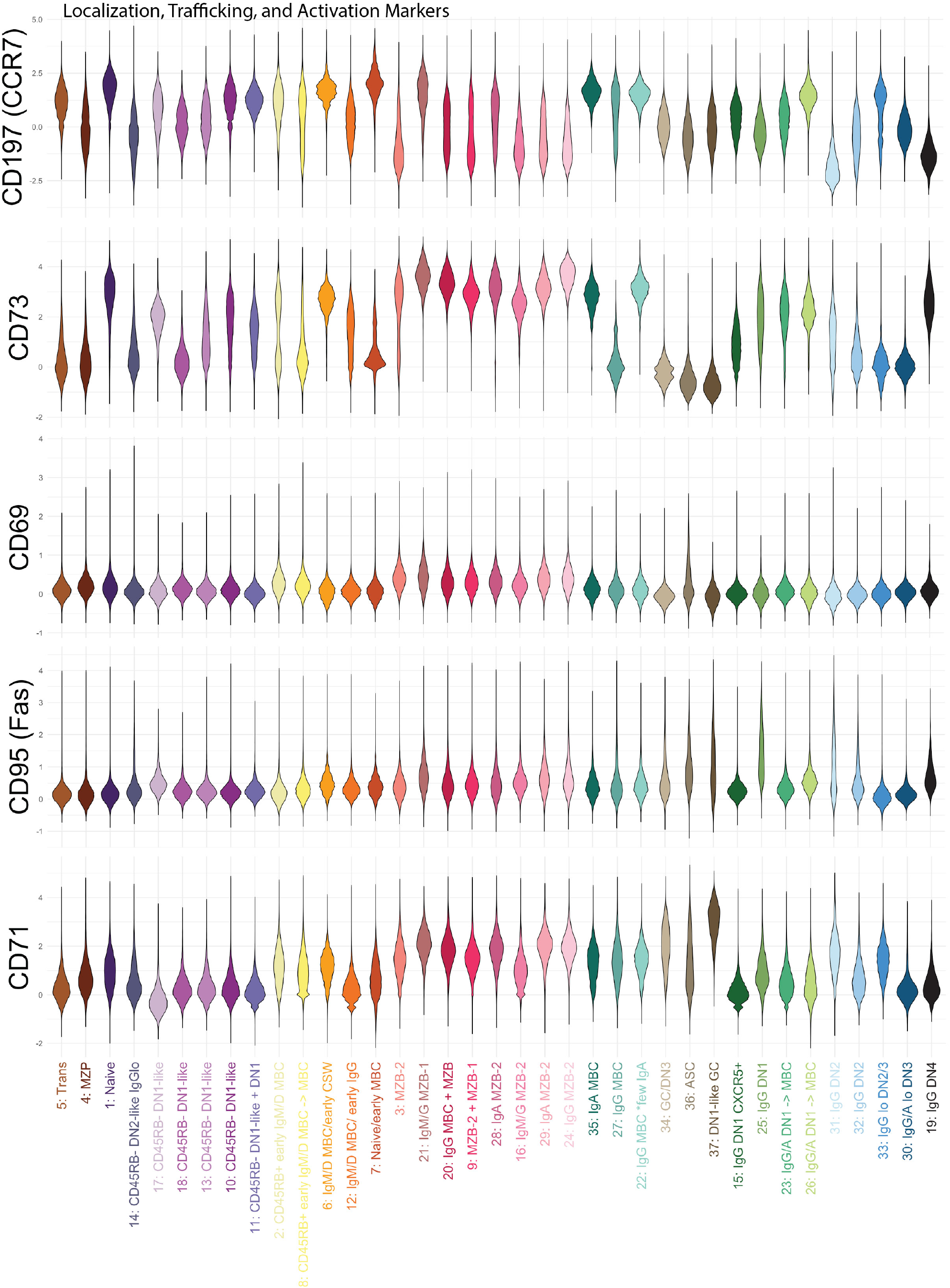
Violin plots of marker expression by cluster for k=37.

**Figure 14f.**
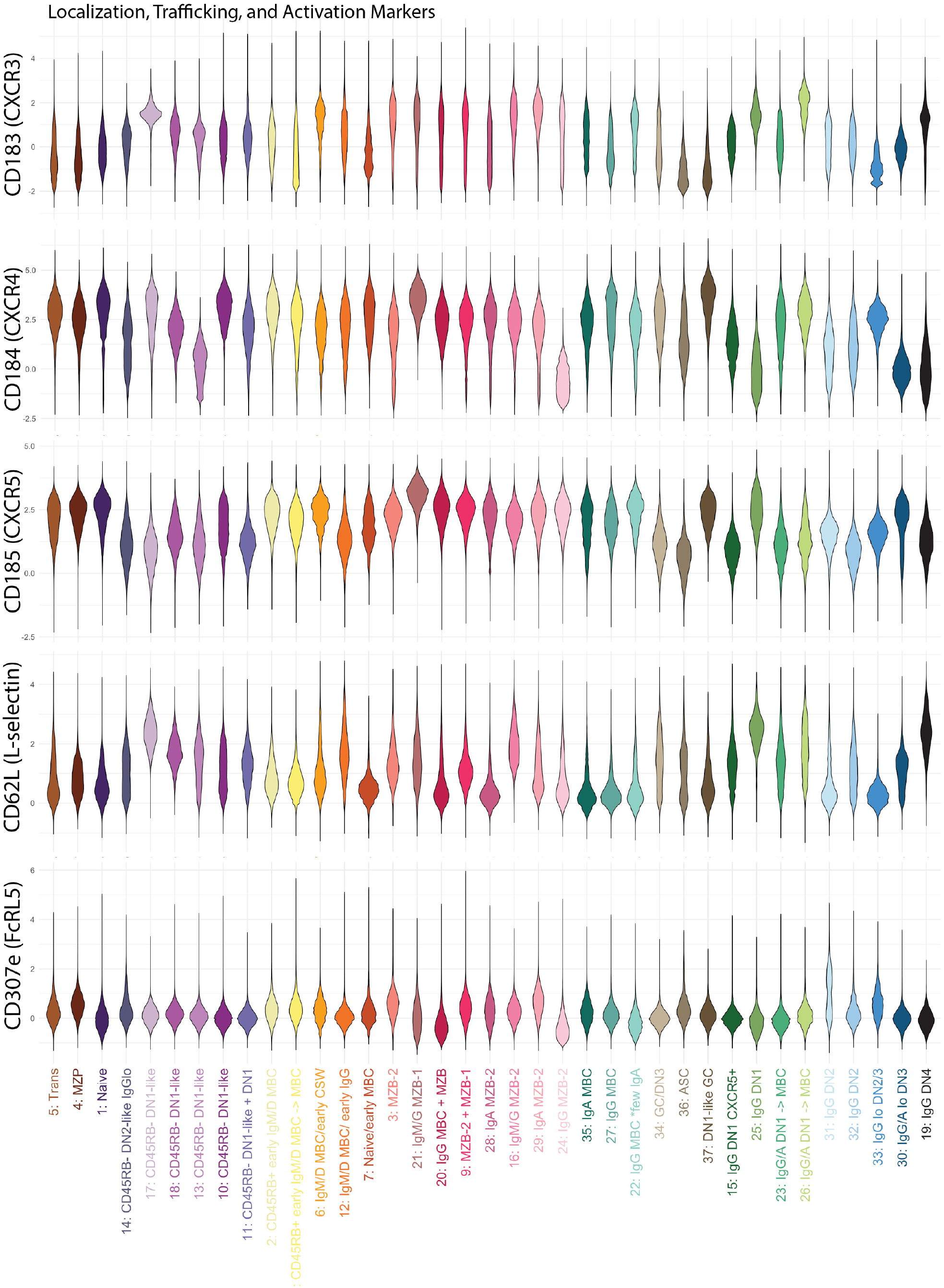
Violin plots of marker expression by cluster for k=37.

## Notes

### Competing Interest Statement

The authors have declared no competing interest.

## REFERENCES

1. Pabst R. enClinical relevance of the compartments and lymphocyte subsets in the human spleen. Cell and Tissue Research 402 (2025) 109–119. doi:10.1007/s00441-025-04001-0.

2. Bagnara D, Squillario M, Kipling D, Mora T, Walczak AM, Da Silva L, et al. A reassessment of igm memory subsets in humans. The Journal of Immunology 195 (2015) 3716–3724. doi:10.4049/jimmunol.1500753.

3. Kibler A, Budeus B, Homp E, Bronischewski K, Berg V, Sellmann L, et al. Systematic memory b cell archiving and random display shape the human splenic marginal zone throughout life. J Exp Med 218 (2021). doi:10.1084/jem.20201952.

4. Gates AJ, Wood IB, Hetrick WP, Ahn YY. Element-centric clustering comparison unifies overlaps and hierarchy. Sci Rep 9 (2019) 8574. doi:10.1038/s41598-019-44892-y.

5. Siu JH, Pitcher MJ, Tull TJ, Velounias RL, Guesdon W, Montorsi L, et al. enTwo subsets of human marginal zone B cells resolved by global analysis of lymphoid tissues and blood. Science Immunology 7 (2022) eabm9060. doi:10.1126/sciimmunol.abm9060.

6. Zhao Y, Uduman M, Siu JHY, Tull TJ, Sanderson JD, Wu YCB, et al. enSpatiotemporal segregation of human marginal zone and memory B cell populations in lymphoid tissue. Nature Communications 9 (2018) 3857. doi:10.1038/s41467-018-06089-1.

7. Weller S, Braun MC, Tan BK, Rosenwald A, Cordier C, Conley ME, et al. Human blood IgM “memory” B cells are circulating splenic marginal zone B cells harboring a prediversified immunoglobulin repertoire. Blood 104 (2004) 3647–3654. doi:10.1182/blood-2004-01-0346.

8. Colombo M, Cutrona G, Reverberi D, Bruno S, Ghiotto F, Tenca C, et al. enExpression of Immunoglobulin Receptors with Distinctive Features Indicating Antigen Selection by Marginal Zone B Cells from Human Spleen. Molecular Medicine 19 (2013) 294–302. doi:10.2119/molmed.2013.00069.

9. Descatoire M, Weller S, Irtan S, Sarnacki S, Feuillard J, Storck S, et al. Identification of a human splenic marginal zone B cell precursor with NOTCH2-dependent differentiation properties. Journal of Experimental Medicine 211 (2014) 987–1000. doi:10.1084/jem.20132203.

10. Ettinger R, Sims GP, Robbins R, Withers D, Fischer RT, Grammer AC, et al. IL-21 and BAFF/BLyS Synergize in Stimulating Plasma Cell Differentiation from a Unique Population of Human Splenic Memory B Cells1. The Journal of Immunology 178 (2007) 2872–2882. doi:10.4049/jimmunol.178.5.2872.

11. Chung MKY, Gong L, Kwong DL, Lee VH, Lee AW, Guan X, et al. enFunctions of double-negative B cells in autoimmune diseases, infections, and cancers. EMBO Molecular Medicine 15 (2023) e17341. doi:10.15252/emmm.202217341.

12. Knox JJ, Buggert M, Kardava L, Seaton KE, Eller MA, Canaday DH, et al. enT-bet+ B cells are induced by human viral infections and dominate the HIV gp140 response. JCI Insight 2 (2017) e92943. doi:10.1172/jci.insight.92943.

13. Tangye SG. enDo multiple subsets of CD11c+ B cells exist? You (T)-Bet! Journal of Allergy and Clinical Immunology 152 (2023) 607–609. doi:10.1016/j.jaci.2023.07.004.

14. Jenks SA, Cashman KS, Zumaquero E, Marigorta UM, Patel AV, Wang X, et al. Distinct effector B-cells induced by unregulated Toll-like receptor 7 contribute to pathogenic responses in Systemic Lupus Erythematosus. Immunity 49 (2018) 725–739.e6. doi:10.1016/j.immuni.2018.08.015.

15. Woodruff MC, Ramonell RP, Nguyen DC, Cashman KS, Saini AS, Haddad NS, et al. enExtrafollicular B cell responses correlate with neutralizing antibodies and morbidity in COVID-19. Nature Immunology 21 (2020) 1506–1516. doi:10.1038/s41590-020-00814-z.

16. Nitschke E, Dang VD, Rincon-Arevalo H, Szelinski F, Ritter J, Schrezenmeier E, et al. EnglishPhosphatidylcholine-specific B cells are enriched among atypical CD11chigh and CD21low memory B cells in antiphospholipid syndrome. Frontiers in Immunology 16 (2025). doi:10.3389/fimmu.2025.1585953.

17. Allard-Chamard H, Kaneko N, Bertocchi A, Sun N, Boucau J, Kuo HH, et al. enExtrafollicular IgDCD27CXCR5CD11c DN3 B cells infiltrate inflamed tissues in autoimmune fibrosis and in severe COVID-19. Cell Reports 42 (2023) 112630. doi:10.1016/j.celrep.2023.112630.

18. Hopp CS, Skinner J, Anzick SL, Tipton CM, Peterson ME, Li S, et al. Atypical B cells upregulate co-stimulatory molecules during malaria and secrete antibodies with T follicular helper cell support. Science immunology 7 (2022) eabn1250. doi:10.1126/sciimmunol.abn1250.

19. Reyes RA, Batugedara G, Dutta P, Reers AB, Garza R, Ssewanyana I, et al. Atypical b cells consist of subsets with distinct functional profiles. iScience 26 (2023) 108496. doi:10.1016/j.isci.2023.108496.

20. Montorsi L, Siu JHY, Spencer J. enB cells in human lymphoid structures. Clinical and Experimental Immunology 210 (2022) 240–252. doi:10.1093/cei/uxac101.

21. Palm AKE, Henry C. enRemembrance of Things Past: Long-Term B Cell Memory After Infection and Vaccination. Frontiers in Immunology 10 (2019) 1787. doi:10.3389/fimmu.2019.01787.

22. Stewart A, Ng JCF, Wallis G, Tsioligka V, Fraternali F, Dunn-Walters DK. Single-Cell Transcriptomic Analyses Define Distinct Peripheral B Cell Subsets and Discrete Development Pathways. Frontiers in Immunology 12 (2021) 602539. doi:10.3389/fimmu.2021.602539.

23. Ruffin AT, Casey AN, Kunning SR, MacFawn IP, Liu Z, Arora C, et al. Dysfunctional CD11cCD21 extrafollicular memory B cells are enriched in the periphery and tumors of patients with cancer. Science Translational Medicine 17 (2025) eadh1315. doi:10.1126/scitranslmed.adh1315.

24. Thorarinsdottir K, Camponeschi A, Gjertsson I, Mårtensson IL. enCD21/low B cells: A Snapshot of a Unique B Cell Subset in Health and Disease. Scandinavian Journal of Immunology 82 (2015) 254–261. doi:10.1111/sji.12339. eprint: https://onlinelibrary.wiley.com/doi/pdf/10.1111/sji.12339.

25. Martínez LE, Comin-Anduix B, Gü emes-Aragon M, Ibarrondo J, Detels R, Mimiaga MJ, et al. EnglishCharacterization of unique B-cell populations in the circulation of people living with HIV prior to non-Hodgkin lymphoma diagnosis. Frontiers in Immunology 15 (2024). doi:10.3389/fimmu.2024.1441994.

26. Budeus B, Kibler A, Küppers R. EnglishHuman IgM–expressing memory B cells. Frontiers in Immunology 14 (2023). doi:10.3389/fimmu.2023.1308378.

27. Perugino CA, Liu H, Feldman J, Marbourg J, Guy TV, Hui A, et al. Two distinct durable human class-switched memory B cell populations are induced by vaccination and infection. Cell Reports 44 (2025) 115472. doi:10.1016/j.celrep.2025.115472.

28. Sosa-Hernández VA, Torres-Ruíz J, Cervantes-Díaz R, Romero-Ramírez S, Páez-Franco JC, Meza-Sánchez DE, et al. B Cell Subsets as Severity-Associated Signatures in COVID-19 Patients. Frontiers in Immunology 11 (2020) 611004. doi:10.3389/fimmu.2020.611004.

29. Sanz I, Wei C, Jenks SA, Cashman KS, Tipton C, Woodruff MC, et al. enChallenges and Opportunities for Consistent Classification of Human B Cell and Plasma Cell Populations. Frontiers in Immunology 10 (2019) 2458. doi:10.3389/fimmu.2019.02458.

30. Velounias RL, Tull TJ. Human b-cell subset identification and changes in inflammatory diseases. Clinical and Experimental Immunology 210 (2022) 201–216. doi:10.1093/cei/uxac104.

31. DiLillo DJ, Palese P, Wilson PC, Ravetch JV. Broadly neutralizing anti-influenza antibodies require fc receptor engagement for in vivo protection. J Clin Invest 126 (2016) 605–10. doi:10.1172/JCI84428.

32. Vidal-Pedrola G, Naamane N, Cameron JA, Pratt AG, Mellor AL, Isaacs JD, et al. Characterization of age-associated b cells in early drug-naïve rheumatoid arthritis patients. Immunology 168 (2022). doi:10.1111/imm.13598.

33. Nellore A, Zumaquero E, Scharer CD, Fucile CF, Tipton CM, King RG, et al. enA transcriptionally distinct subset of influenza-specific effector memory B cells predicts long-lived antibody responses to vaccination in humans. Immunity 56 (2023) 847–863.e8. doi:10.1016/j.immuni.2023.03.001.

34. Crickx E, Chappert P, Sokal A, Weller S, Azzaoui I, Vandenberghe A, et al. enRituximab-resistant splenic memory B cells and newly engaged naive B cells fuel relapses in patients with immune thrombocytopenia. Science Translational Medicine 13 (2021) eabc3961. doi:10.1126/scitranslmed.abc3961.

35. Monaco G, Chen H, Poidinger M, Chen J, De Magalhães JP, Larbi A. enflowAI: automatic and interactive anomaly discerning tools for flow cytometry data. Bioinformatics 32 (2016) 2473–2480. doi:10.1093/bioinformatics/btw191.

36. Hammill D. Cytoexplorer: Interactive analysis of cytometry data (2021). R package version 1.1.0.

37. Azad A, Rajwa B, Pothen A. flowVS: channel-specific variance stabilization in flow cytometry. BMC Bioinformatics 17 (2016) 291. doi:10.1186/s12859-016-1083-9.

38. Van Gassen S, Gaudilliere B, Angst MS, Saeys Y, Aghaeepour N. CytoNorm: A Normalization Algorithm for Cytometry Data. Cytometry 97 (2020) 268–278. doi:10.1002/cyto.a.23904.

39. Van Gassen S, Callebaut B, Van Helden MJ, Lambrecht BN, Demeester P, Dhaene T, et al. Flowsom: Using self-organizing maps for visualization and interpretation of cytometry data. Cytometry Part A 87 (2015) 636–645. doi:10.1002/cyto.a.22625.

40. Charrad M, Ghazzali N, Boiteau V, Niknafs A. enNbClust: An R Package for Determining the Relevant Number of Clusters in a Data Set. Journal of Statistical Software 61 (2014) 1–36. doi:10.18637/jss.v061.i06.

41. Wilkerson MD, Hayes DN. ConsensusClusterPlus: a class discovery tool with confidence assessments and item tracking. Bioinformatics 26 (2010) 1572–1573. doi:10.1093/bioinformatics/btq170.

42. [Dataset] Melville J, Lun A, Djekidel MN, Hao Y, Eddelbuettel D, Bijl Wvd. uwot: The Uniform Manifold Approximation and Projection (UMAP) Method for Dimensionality Reduction (2025).

43. Tull TJ, Pitcher MJ, Guesdon W, Siu JH, Lebrero-Fernández C, Zhao Y, et al. Human marginal zone B cell development from early T2 progenitors. The Journal of Experimental Medicine 218 (2021) e20202001. doi:10.1084/jem.20202001.

44. Hanson CH, Henry B, Andhare P, Lin FJ, Pak H, Turner JS, et al. CD62L expression marks a functionally distinct subset of memory B cells. Cell reports 42 (2023) 113542. doi:10.1016/j.celrep.2023.113542.

45. Aghaeepour N, Simonds EF, Knapp DJHF, Bruggner RV, Sachs K, Culos A, et al. GateFinder: projection-based gating strategy optimization for flow and mass cytometry. Bioinformatics 34 (2018) 4131–4133. doi:10.1093/bioinformatics/bty430.

46. Hahne F, LeMeur N, Brinkman RR, Ellis B, Haaland P, Sarkar D, et al. enflowCore: a Bioconductor package for high throughput flow cytometry. BMC Bioinformatics 10 (2009) 106. doi:10.1186/1471-2105-10-106.

47. Vinh NX, Epps J, Bailey J. Information theoretic measures for clusterings comparison: Variants, properties, normalization and correction for chance. Journal of Machine Learning Research 11 (2010) 2837–2854.

